# PARAS: high-accuracy machine learning of substrate specificities in nonribosomal peptide synthetases

**DOI:** 10.1101/2025.01.08.631717

**Authors:** Barbara R. Terlouw, Chuan Huang, David Meijer, José D. D. Cediel-Becerra, Ruolin He, Marlene L. Rothe, Matthew Jenner, Shanshan Zhou, Yu Zhang, Christopher D. Fage, Yuta Tsunematsu, Gilles P. van Wezel, Serina L. Robinson, Fabrizio Alberti, Lona M. Alkhalaf, Marc G. Chevrette, Gregory L. Challis, Marnix H. Medema

**Affiliations:** Bioinformatics Group, Department of Plant Science, Wageningen University & Research, Wageningen, The Nether-lands; Institute of Biology, Leiden University, Leiden, The Netherlands; Department of Chemistry, University of Warwick, Coventry, United Kingdom; Department of Biochemistry and Molecular Biology, Biomedicine Discovery Institute, Monash University, Clayton, Australia; ARC Centre of Excellence for Innovations in Peptide and Protein Science, Monash University, Clayton, Australia; Department of Microbiology and Cell Science, University of Florida, Gainesville, Florida, USA; School of Life Sciences, University of Warwick, Coventry, United Kingdom; Graduate School of Bioagricultural Sciences, Nagoya University, Furo-cho, Chikusa, Nagoya, Aichi, Japan; Department of Environmental Microbiology, Swiss Federal Institute of Aquatic Science and Technology (EAWAG), Dübendorf, Switzerland; Department of Plant Pathology and Wisconsin Institute for Discovery, University of Wisconsin-Madison, Madison, WI, USA

**Keywords:** PARAS, PARASECT, NRPS adenylation domain prediction, intact protein mass spectrometry, XAI

## Abstract

Nonribosomal peptides are diverse natural products with important applications in medicine and agriculture. Bacterial and fungal genomes contain thousands of nonribosomal peptide biosynthetic gene clusters (BGCs) of unknown function, providing a promising resource for peptide discovery. Core structural features of such peptides can be inferred by predicting the substrate(s) of adenylation (A) domains in nonribosomal peptide synthetases (NRPSs). However, existing approaches to A domain prediction rely on limited datasets and often struggle with domains selecting large substrates or from less-studied taxa. Here, we systematically curate and computationally analyse 3,653 A domains and present two high-accuracy specificity predictors, PARAS and PARASECT. A type of A domain with unusually high L-tryptophan specificity was identified through the application of PARAS, and intact protein mass spectrometry to the corresponding NRPS showed it to direct the production of tryptopeptin-related metabolites in *Streptomyces* species. Together, these technologies will accelerate the characterisation of novel NRPSs and their metabolic products. PARAS and PARASECT are available at https://paras.bioinformatics.nl.

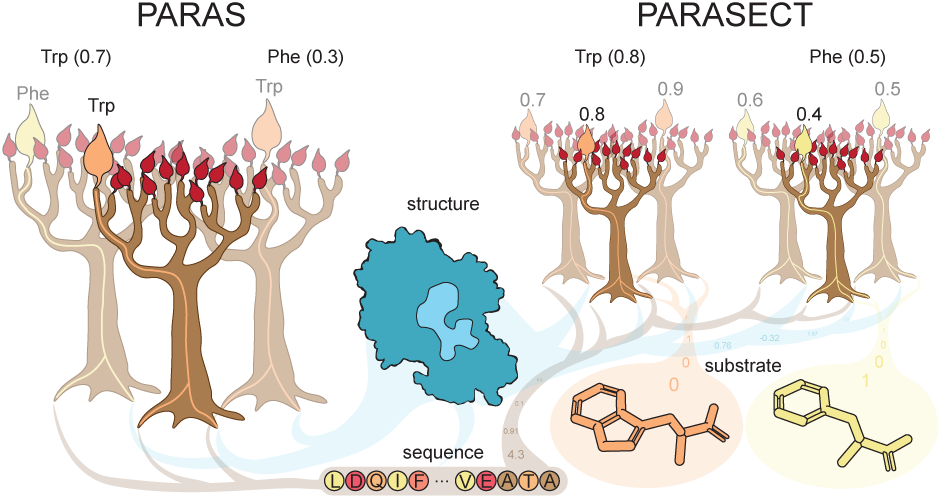

## INTRODUCTION

Bacteria and fungi produce a multitude of specialised metabolites, including numerous structurally and functionally diverse non-ribosomal peptides with important applications in medicine and agriculture. These include the antibiotic daptomycin^1^, the immunosuppressant cyclosporine^2^, the fungicide UK-2A^3^, and the herbicide Bialaphos^4^. Also, nonribosomal peptides such as lugdunin^5^, colibactin^6^ and thanamycin^7^ have been found to be key mediators of (host-)microbiome interactions.

Giant modular multienzymes called nonribosomal peptide synthetases (NRPSs) assemble these natural products. Each NRPS module contains an adenylation (A) domain, which selectively binds an amino (or other) acid and catalyses formation of an aminoacyl adenylate by reaction with ATP, and a peptidyl carrier protein (PCP) domain that is post-translationally modified via attachment of a coenzyme derived phosphopantetheine (PPant) arm to a conserved Ser residue. Nucleophilic attack of the thiol group at the terminus of the PPant arm on the activated carboxyl group of the aminoacyl adenylate results in formation of an aminoacyl thioester. In chain-elongating modules, condensation (C) domains catalyse peptide bond formation between aminoacyl thioesters attached to PCP domains in biosynthetically adjacent modules^8^. While ribosomally-biosynthesised peptides are assembled from the 20 proteinogenic amino acids, NRPSs can incorporate over 300 different building blocks^9^, including proteinogenic and non-proteinogenic amino acids, alpha-keto and alpha-hydroxy acids, various N-terminal acyl ‘caps’, and C-terminal amines and alcohols. This, combined with a large array of on- and post-NRPS tailoring domains/enzymes, and the ability of NRPSs to work together with type I modular polyketide synthases (PKSs), results in a wide range of modifications to the peptide scaffold and yields structurally diverse metabolites with immense functional potential.

In the late 1990s, Stachelhaus *et al.* and Challis and co-workers independently identified residues in A domain active sites predicted to control substrate selectivity. Based on the crystal structure of an A domain from the first module of gramicidin S synthetase A with its cognate L-Phe substrate and AMP bound, they converged on a core set of nine selectivity-conferring amino acid residues^10^, with Stachelhaus *et al.* proposing an additional tenth^11^. Stachelhaus *et al.* proposed that the selectivity-conferring residues constitute a “nonribosomal code” (sometimes referred to as the Stachelhaus code), likening it to the genetic code used as a blueprint for assembling proteins. Challis and co-workers, however, viewed this as an over-simplification, emphasising that these residues form a complex three-dimensional network of precisely positioned functional groups able to recognise diverse substrates using a combination of electrostatic, hydrophobic, and hydrogen bond interactions. This approach revealed additional nuances, such as distinct motifs for recognition of ornithine, *N*5-hydroxyornitione and *N*5-hydroxy-*N*5-formylornithine.

In a 2004 review, Challis and co-workers highlighted that recognition of substrates with smaller side chains than L-Phe likely requires fewer than nine or ten residues^12^. Models of two types of L-Pro-activating A domains with distinct sets of specificity conferring residues were used to illustrate this^12^. Subsequent X-ray crystallographic studies of an A domain with its cognate L-Thr substrate / ATP and the corresponding adenylate bound provided experimental confirmation of <9-10 residues required for selectivity^13^. Conversely, recognition of substrates with larger sides chains than L-Phe would be expected to require more than 10 residues. Taken together, these observations undermined the notion of a one-size-fits-all “nonribosomal code” for prediction of A domain substrate selectivity. Notwithstanding these limitations, Challis and co-workers applied their model for A domain substrate recognition to prediction of the amino acids selected by a cryptic NRPS encoded by the *S. coelicolor* genome^14^. Subsequent identification of the novel siderophore coelichelin as the product of this NRPS verified the accuracy of these predictions, heralding the dawn of genome mining as a new approach for discovery of novel nonribosomal peptides^15^. With (meta)genome sequencing costs at an all-time low, genome mining has now assumed a key role in natural product discovery^16,17^. Tools like antiSMASH^18,19^ have enabled researchers worldwide to perform millions of analyses to explore sequenced genomes for specialised metabolite BGCs.

Subsequently, other approaches to predict A domain substrate selectivity have been developed, including profile Hidden Markov Model (pHMM)-based methods^15,20^ and various machine learning algorithms trained on extended, 34-residue signatures, including support vector machines (SVMs)^21,22^, random forests^23,24^, and ensemble methods that use a combination of these approaches^25^. While methods using these extended 34-residue signatures have a better chance of capturing all residues that are involved in substrate recognition, the available tools deploying them for substrate prediction have a common limitation: a lack of high-quality training data. The SANDPUMA and AdenPredictor algorithms are based on the largest training set of only ∼1000 data points^24,25^. NRPSPredictor2, which is still employed by the widely used BGC annotation pipeline antiSMASH, was only trained on ∼550 data points, with 9 out of the 30 most common substrates being covered by <10 training data points. Also, evolutionarily independent clades of A domains with very different active site architectures can recognise identical substrates, highlighting a need for a large and diverse dataset that covers as many A domain clades as possible. Furthermore, current predictors only consider sequence features of the A domain and do not explore structural features of either the enzyme or the substrate. As such, these algorithms cannot learn from the three-dimensional configurations that ultimately govern substrate selectivity, nor can they use information about substrate similarity. Finally, current predictors were trained on datasets that contained data points labelled with multiple possible substrates, which can further conflate prediction accuracies.

Some A domains are naturally promiscuous, particularly those activating large hydrophobic substrates such as L-Phe, L-Leu, L-Ile, and L-Val. However, the ATP-pyrophosphate exchange assay commonly used to probe substrate selectivity has several shortcomings that misleadingly suggest many A domains are promiscuous. First, it does not permit competition between multiple substrates, as encountered by A domains *in cellulo*, to be assessed; only the measurement of their relative ability to adenylate isolated substrates. Second, it only measures the adenylation reaction, providing no means of determining whether the subsequent thiolation reaction also contributes to substrate selectivity^26^. NRPS condensation domains catalyse peptide bond formation between PCP-domain-bound amino-acyl (and peptidyl) thioesters. It is thus the overall selectivity of an A domain for formation of different aminoacyl thioesters that primarily dictates the composition of NRPS products. Current state-of-the-art algorithms are not true multi-label predictors, representing the selectivity of “promiscuous” A domains as unique multi-substrate strings corresponding to unique specificities, rather than substrate sets.

Here, we use sequence comparisons and AlphaFold^27^ structural models of thousands of A domains to reveal widespread independent evolution of domains with similar substrate selectivity. Motivated by our observations and powered by large-scale training data curation, we developed two fast and accurate machine learning frameworks: PARAS (Predictive Algorithm for Resolving Adenylation domain Selectivity) and PARASECT (Predictive Algorithm for Resolving Adenylation domain Selectivity by featurising Enzyme and Compound in Tandem). These algorithms overcome challenges with previously developed predictors, using an expanded and manually curated training set three times the size of previous datasets, providing options for structure-guided feature extraction, integrating substrate features, and accommodating reliable multi-label predictions. We benchmark PARAS and PARASECT against AdenPredictor, SANDPUMA, and NRPSPredictor2, showing a substantial increase in predictive accuracy on independent holdout data (improving by 20%, 27%, and 22%, respectively). Feature importance analysis shows how different residues are predictive for specificity of A domains accepting different substrates, but also for independently evolved clades of A domains that accept the same substrates. Subsequently, we showcase how PARAS and PARASECT accurately predict L-Trp as the substrate of an unusual A domain in a novel NRPS shown to assemble metabolites related to the immunomodulator tryptopeptin A. Intact protein mass spectrometry (MS) analyses of aminoacyl thioester formation on the downstream *holo*-PCP domain associated with this A domain confirm it has high specificity for L-Trp and highlight the importance of competition experiments for developing an accurate picture of physiologically relevant A domain substrate selectivity.

## RESULTS

### An expanded and highly curated training set of A domain specificities

The performance of predictive models largely relies on the size of the dataset on which they are trained. For this reason, we collated an A domain dataset from three sources: MIBiG 3.0^28^, the SANDPUMA/AdenPredictor (SP/AP) training set^24,25^, and the NRPSPredictor2 training set^22^. As fungal domains were underrepresented in the resulting dataset (252 compared to 2900 bacterial domains), we annotated an additional 494 fungal domains from MIBiG 4.0^29^ and literature^30–51^. The resulting combined dataset counted 3,653 A domains from diverse phyla (Figure 1a), recognising 278 different substrates (Figure 1b). This accounts for 3.4 and 6.8 times as many labelled data points as were used for training SP/AP (1089 data points) and NRPSPredictor2 (534 data points), respectively. Data were extensively curated and validated based on detailed phylogenetic and literature analyses through which we discovered and corrected misannotations in the SP/AP and NRPSPredictor2 datasets (137 [12.6%] in SP/AP, 31 [5.8%] in NRPSPredictor2), which will likely have impacted the performance of these models.

**Figure 1.**
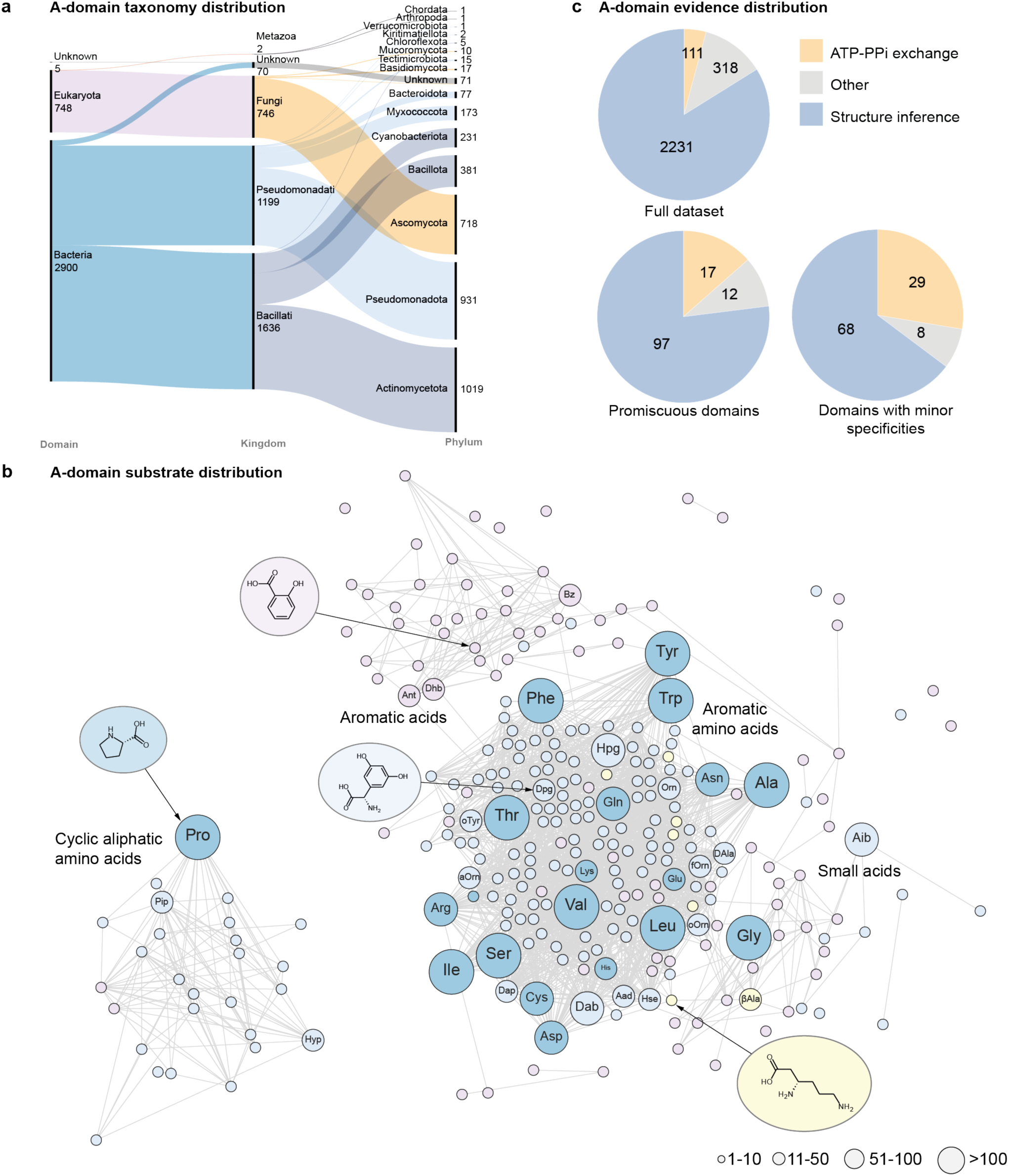
Overview of A domains used for training PARAS and PARASECT. **a.** Alluvial diagram showing the taxonomic distribution of all A domains for which taxonomy was obtained. **b.** Tanimoto similarity network (cutoff=0.46) of substrates activated by the A domains in the dataset. Node size represents the number of times a substrate is present in the dataset. Blue: ⍺-amino acids. Dark blue: proteinogenic amino acids. Yellow: β-amino acids. Pink: Other acids. The 34 substrates labelled with abbreviations are the substrates used for training validation and benchmarking models. **c.** Proportion of A domains annotated using ATP-pyrophosphate exchange assays across the full dataset, A domains with multiple major substrate specificity annotations, and A domains with minor substrate specificity annotations. Aad: 2-aminoadipic acid. Dab: 2,4-diaminobutyric acid. Aib: 2-aminoisobutyric acid. βAla: β-alanine. Dhb: 2,3-dihydroxybenzoic acid. D-Ala: D-alanine. Orn: ornithine. fOrn: N5-formyl-N5-hydroxyornithine. oOrn: N5-hydroxyornithine. aOrn: N5-acetyl-N5-hydroxyornithine. Hpg: 4-hydroxyphenylglycine. Dpg: 3,5-dihydroxyphenylglycine. oTyr: (R)-β-hydroxytyrosine. Pip: pipecolic acid. Ant: anthranilic acid. Sal: sali-cylic acid.

Metadata were available from the MIBiG database for a subset of these A domains, providing i) the evidence that was used to obtain each substrate annotation, and ii) A domain selectivity classified as major and minor substrate(s). The majority of annotations were based on structural inferences of the peptide scaffolds produced *in vivo*, and less frequently by high levels of activation (>80% activation compared to the predominant substrate) in A domain specificity assays, such as ATP-pyrophosphate exchange^28^. Of the 2,660 A domains with available metadata, 111 (∼4%) were elucidated using ATP-pyrophosphate exchange assays. For promiscuous A domains with multiple major substrates, this proportion was substantially higher (17/126; ∼13%). For promiscuous A domains with one or more minor substrates in addition to their major substrate, this amounted to an even greater proportion of ∼28% (29/105; Figure 1c). These differences suggest that A domain promiscuity is overestimated by the ATP-pyrophosphate exchange assay, which can report false positives (as discussed in the introduction), with many activated substrates never being transferred to the cognate PCP domain *in cellulo* and therefore never being incorporated into the NRP scaffold. Thus, we chose to remove all minor substrates from our dataset.

### Large-scale analysis of structural models reveals independent evolution of A domain substrate specificity

We next investigated other shortcomings of current state-of-the-art A domain predictors. We observed that they disproportionately made errors in the classification of large proteinogenic amino acids such as phenylalanine, tryptophan, and lysine^22,24^ (F1-score of 0.688, 0.320 and 0.400 in NRPSPredictor2, respectively; most common mispredictions in AdenPredictor). We hypothesised that due to the size and degrees of rotational freedom of the side chains of large substrates, parallel evolution might have given rise to A domain active site pockets that select identical substrates using different active site architectures.

To explore this hypothesis, we constructed AlphaFold models for 3,254 A domains and explored the diversity of their active site pockets using principal component analysis on their three-dimensional voxel grids (see Online Methods). We observed that A domains selecting substrates for which current predictors perform well (e.g., threonine, and serine), cluster together in one or a few major groups, indicating high active site similarity (Figure 2a). In contrast, A domains selecting large substrates (e.g., phenylalanine, tryptophan, and lysine), were predicted to have divergent active site pockets (Figure 2b), suggesting multiple active site architectures for these substrates, possibly resulting from parallel evolution. This was further supported by phylogenetic analysis of the 34 amino acid extended active site signatures, which demonstrates that A domains recognising large substrates are dispersed across multiple different monophyletic clades, while 41% of Thr-incorporating A domains fall into a single monophyletic clade and 75% of Thr-incorporating A domains fall into the largest five clades (Figure 2c).

**Figure 2.**
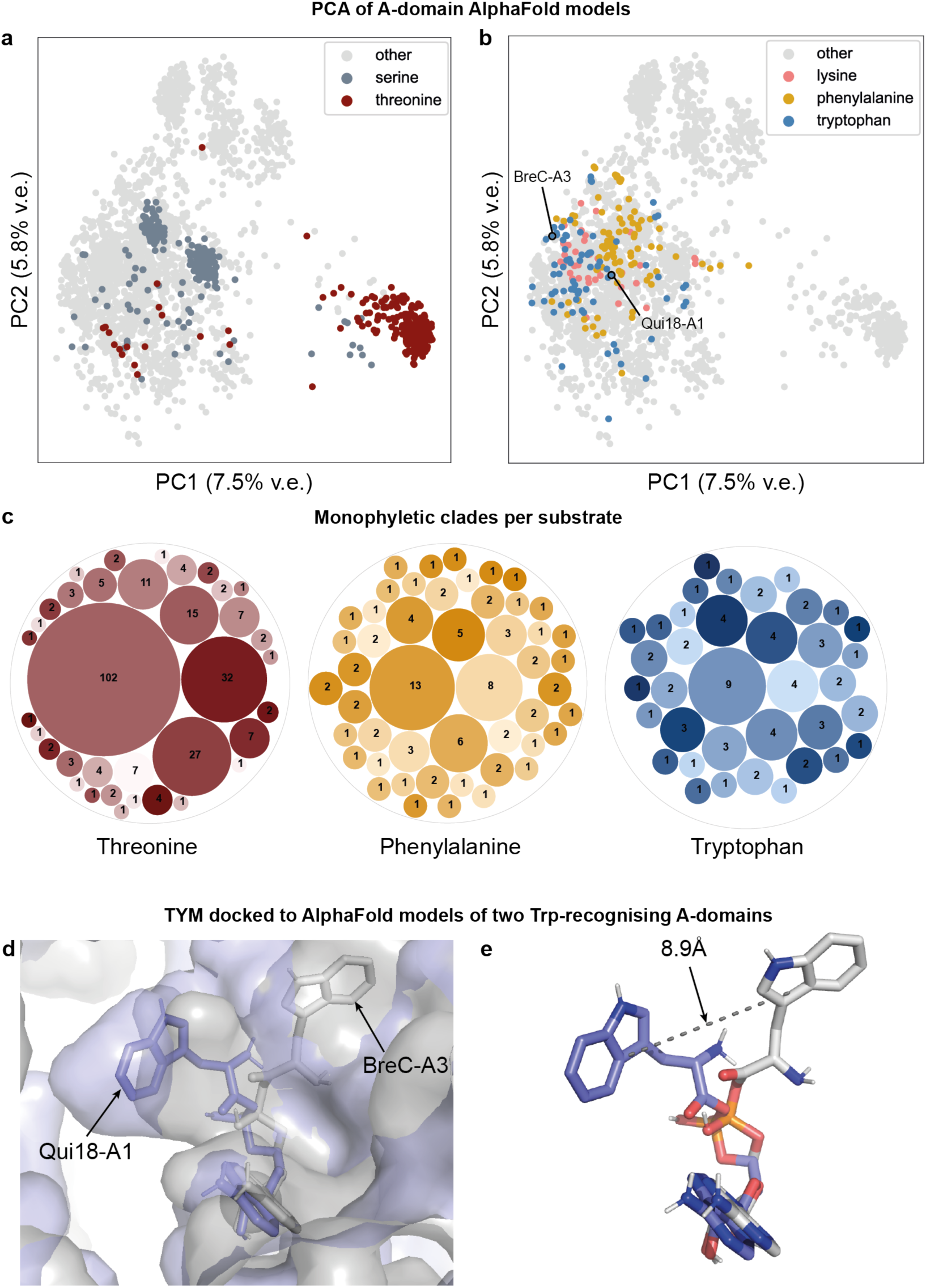
Large-scale structure and sequence analysis of A domain active site diversity. **a.** PCA of A domain active sites based on 3,254 AlphaFold models, with those incorporating the polar substrates Ser and Thr highlighted, and **b.** A domains incorporating the large substrates Phe, Trp, and Lys highlighted. Trp-incorporating domains Qui18-A1 and BreC-A3 are indicated. v.e.: Variance explained **c.** Bubble representation of the number per monophyletic clade for Thr, Trp, and Phe-incorporating A domains, based on a phylogenetic tree of the 34 amino acid extended active site signatures. Each bubble represents one monophyletic clade, with the size of each bubble proportional to the number of A domains in the monophyletic clade. **d.** Comparison of TYM docked to AlphaFold-modelled Qui18-A1 (blue) and BreC-A3 (grey). The left pocket is smaller in BreC-A3 (grey) than in Qui18-A1 (blue), making it unlikely for the tryptophan substrate to fit into this pocket in BreC-A3. Instead, the tryptophan moiety of TYM docked to BreC-A3 localises to a pocket on the right, leading to a 8.9 Å shift (d; dashed line) of the indole C3 atom in the bound TYM relative to the Qui18-A1 docked structure.

To visually illustrate the nature of active site pocket differences that may be observed between A domains in phylogenetically distinct branches, we compared the predicted active sites of two Trp-incorporating A domains. We chose the A domain from Qui18 (GenPept accession AET98916.1; Qui18-A1), an NRPS involved in quinomycin biosynthesis in *Streptomyces griseovariabilis subsp. bandungensis*^52^; and the third A domain from BreC (GenPept accession ATY37608.1; BreC-A3), an NRPS involved in the biosynthesis of brevicidine in *Brevibacillus laterosporus*^53^. The 10-residue active site signatures of these A domains are highly divergent: DAWTVTGVGK for Qui18-A1 and DPTQAGEVVK for BreC-A3, only sharing three common residues, two of which are the highly conserved Asp and Lys residues that form salt bridges with the backbone ammonium and carboxylate groups in the majority of A domains. Their extended 34-residue active site signatures, which are used by NRPSPredictor2, SANDPUMA, and AdenPredictor, are also highly dissimilar, with only 10 out of 34 residues in common and a pairwise alignment bitscore of 31.0, compared to a score of 184.0 for the two most similar tryptophan-recognising A domains in our dataset. The distance between both domains in a principal components analysis (PCA; Figure 2b), as well as docking of Trp-adenylate (TYM) to the AlphaFold models for both A domains predicts that these differences are manifest in their 3D structures, further suggesting A domains can bind the side chains of large substrates using architecturally diverse pockets (Supplementary Discussion 1; Figure 2d, 2e; Figure S23).

This underscores the need for a sufficiently varied training dataset to cover A domains spanning different phylogenetic branches.

### Improving A domain substrate prediction: structure-based approaches and substrate featurisation

In addition to increasing the size of our training set, extensive data curation, and eliminating unreliable annotations of substrate promiscuity, we explored two avenues for improving A domain substrate prediction: including featurising the A domain and substrate in tandem to allow multi-label predictions (Figure 3a,c) and using protein structure-based features (Figure 3b).

**Figure 3.**
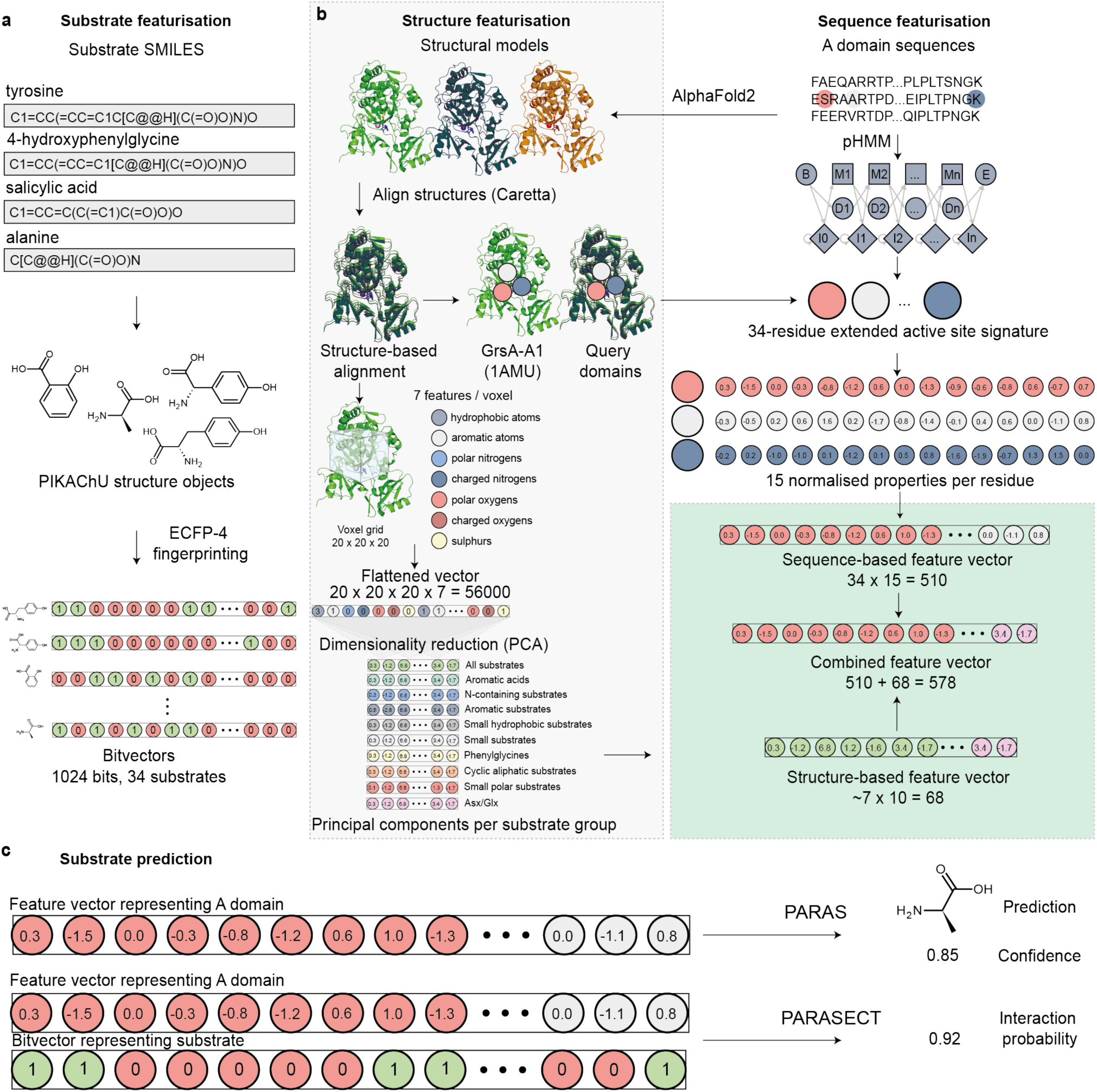
Overview of PARAS and PARASECT workflow. **a.** Feature extraction for substrates, used in PARASECT models. SMILES strings of substrates are first processed and converted into PIKAChU structure graphs. Then, Extended Connectivity Fingerprinting (ECFP-4) is used to generate bit vectors that represent the substrate structure. **b.** Feature extraction for sequence-based, structure-based and combined models. In structure featurisation, structural models are generated for A domain sequences using AlphaFold2, and the resulting structural models are aligned using structure-based alignment. Then, either sequence-based features can be extracted from this structure-guided alignment based on the alignment with the reference GrsA-A1 crystal structure (then proceeding again with the sequence-based workflow), or structure-based features can be extracted using a 3D voxel grid, where physicochemical properties of coordinates across the substrate-binding pocket are encoded before reducing their dimensionality with principal component analysis. In sequence-based featurisation, sequence-based features are directly extracted through alignment to a pHMM along with the GrsA-A1 reference sequence to extract a 34-residue active site signature. For each of the 34 residues, 15 normalised physicochemical properties are computed and encoded in a feature vector. Sequence-based feature vectors can either be used on their own (as by default, to optimise speed) or combined with structure-based feature vectors. **c.** PARAS and PARASECT input and output. The PARAS algorithm is a multi-label classifier that predicts a single substrate for a query A domain, with a given confidence level. In contrast, PARASECT computes interaction probabilities for each and every substrate separately, and these can then be ranked to identify the most likely substrate(s).

Prior to training all models, we made two train-test splits: one by stratifying on substrate, and the other by stratifying on taxonomy at family level. The latter enables us to assess how well our algorithms will perform on sequences that are phylogenetically distinct from any training data. The average protein percent identity of a test data point to its closest relative in the training set was 75%±18% for the class-based split and 59%±9% for the phylogeny-based split. In both cases, the ratio of training data to test data was approximately 3:1 (Figure S1).

With our expanded training set, we explored how different feature selection and extraction methods affect model performance. Models were trained and validated in four steps to select appropriate parameters, and to facilitate substrate-specific analysis and fair comparison to existing A domain predictors (Online Methods). For feature selection, we investigated the amino acid sequence features also used by NRPSPredictor2, AdenPredictor, and SANDPUMA; structural A domain features derived from 3D AlphaFold 2 models; residue-specific Evolutionary Scale Model (ESM)2 embeddings^54^; and Morgan fingerprints^55^ of the substrates, which convert chemical structures into bitvectors representing their substructures. We trained random forest models based on all possible feature combinations (Figure S2). These fell into two model categories: (i) those that only use an A domain as input to predict its most likely substrate (PARAS), and (ii) those that use both an A domain and a putative substrate as input to predict the probability that the A domain selects that substrate (PARASECT).

Importantly, PARAS is only trained on the first-listed substrate, which always corresponds to the substrate incorporated into its validated natural product. Therefore, PARAS does not suffer from potential inaccuracies in the annotation of promiscuous A domains that arise from ATP-pyrophosphate exchange assay data. In contrast, PARASECT was trained on all major substrates listed for each A domain, and is therefore more sensitive to these inaccuracies. We limited the likelihood of such misannotations impacting PARASECT model performance by excluding minor substrates, which reduced the percentage of promiscuous annotations relying on ATP-pyrophosphate exchange data to 13%.

### Structure-based feature extraction provides slight improvements for taxonomic outliers at the expense of speed

Because the shape of an A domain’s active site determines its substrate selectivity ^10,11,56^, we expected that models based on structural features would perform well. However, we observed that models trained on sequence features substantially outperformed those trained on structural features. Using both sequence and structural features did not significantly improve performance compared to using sequence features only (Figure S2), suggesting that there is little extra information contained in the structural features that the model cannot interpret from sequence features. However, feature importance analyses show that in models combining sequence and structural features, many of structural features were among the most frequently used in the model, indicating that certain structural information can be informative (Supplementary Discussion). We also explored residue-specific ESM2 embeddings, which also contain structural information. While these increased predictive power compared to AlphaFold features, they underperformed compared to sequence features by 3-5% (Figure S3).

During cross-validation, we noticed that our sequence-based PARASECT model performed slightly better on bacterial data when fungal data was not included. Conversely, bacterial data did improve performance on fungal data (Figure S4). For this reason, we trained two PARASECT models: one trained on bacterial sequences only (to be used on bacterial data), and one trained on all sequences (to be used on metagenome and fungal data).

Our best model (PARAS, sequence-based) achieved top accuracies of ∼88% (∼92% bacterial, ∼74% fungal) on substrate-stratified data, and ∼83% (∼89% bacterial, ∼48% fungal) on taxonomy-stratified data (Figure S3, S10).

Runtime is an important consideration for an A domain predictor: SANDPUMA was implemented in antiSMASH 4.0 and subsequently removed in antiSMASH 5.0 due to a computational cost of 2-5 minutes per A domain. The inclusion of structural features in PARAS and PARASECT would increase their runtime by ∼20 minutes per A domain, as AlphaFold models would have to be constructed prior to running the model. ESM2 embeddings are less costly, but still require ∼20-30 seconds per domain on a CPU, or a dedicated GPU for computing embeddings more rapidly. As neither structural features nor ESM2 embeddings substantially improved sequence-based PARAS and PARASECT models, which take only a few milliseconds per prediction (Table S2), we opted for sequence-based models for benchmarking.

We also examined structure-informed methods that do not require the modelling of A domains that the user wants to query or the computation of ESM2 features, including structure-guided profile alignment for active site extraction. Compared to extraction using sequence-based profile alignments, as used by SANDPUMA, this method yielded active sites with fewer gaps (Figure S5), which in turn led to models with slightly increased performance (Table S1). However, extracting the active sites using the A domain pHMM from NRPSPredictor2 is much faster (milliseconds rather than seconds per domain), and led to even fewer gaps overall (163 vs 673). Comparison of different active site extraction methods is discussed in more detail in our Supplementary Discussion.

For 35 A domains, structure-guided profile alignment yielded active sites with fewer gaps than pHMM-based extraction. Of these, 17 recognise non-amino acid substrates, 2 tether aspartate to the PCP domain by the side chain carboxyl group, and 14 belong to kingdoms and phyla that are underrepresented in our dataset, including Fungi, Myxococcota, Cyanobacteriota, and various uncultured bacteria from environmental samples. This suggests that structure-guided profile alignment might be better suited for extracting the active sites of A domains recognising non-amino acid substrates and of A domains that are phylogenetically distinct from those in our dataset. For example, for fungal A domains, Heard and Winter demonstrated that structural modeling enhanced substrate preference prediction^57^.

The limitations of current predictors for fungal A domains or non-amino acids are understandable, as the pHMM developed by Rausch *et al.*^21^ was based on a phylogenetically restricted set of A domains that recognise amino acid substrates. Therefore, we added an option to our command line models and our webpage to select either method for sequence feature extraction (Figure S6).

### PARASECT allows predicting multiple or previously unseen A domain substrates

Our PARASECT models provide a novel method of A domain selectivity prediction: rather than solely using A domain features to predict a substrate, PARASECT considers both the A domain and the structure of the substrate and predicts if the domain is compatible with that substrate. This has a number of advantages.

First, the model can learn from the similarity and differences between substrates during training, e.g., how the presence of an aromatic ring in the substrate combined with hydrophobic residues in an A domain active site affects selectivity. We observe that PARASECT’s top predictions (i.e., those substrates with the highest predicted interaction probabilities with an A domain) are often structurally similar, in contrast to PARAS, indicating that the PARASECT model indeed incorporates substrate similarity to make classifications. We confirmed this with feature inference analysis of our cross-validation models, which showed six substrate features as the most informative in the model (Figure S7). Also, as multiple top predictions may be biologically relevant, PARASECT provides a means to obtain a range of substrate predictions for an A domain among which the true substrate is likely present.

Second, PARASECT can be queried with substrates not present in the model’s training data. For example, we used PARASECT to predict the interaction between the A domain in module 6 of the calcium-dependent antibiotic (CDA) NRPS (MIBiG accession BGC0000315), which canonically incorporates 4-hydroxyphenylglycine. This domain is also able to incorporate 4-fluorophenylglycine in mutasynthesis experiments^58^. PARASECT predicted an interaction probability of 54.1%, returning 4-fluorophenylglycine as the second most likely substrate to be incorporated after 4-hydroxyphenylglycine (Figure S9). Notably, PARASECT’s training set does not include fingerprints of substrates containing F atoms. This demonstrates a potential application of PARASECT in targeting A domains for variant creation via precursor-directed biosynthesis. To make this feature available to users, we have included an option to upload substrates in SMILES format^59^ in our web application.

Finally, PARASECT can train on and make predictions for promiscuous A domains that incorporate multiple substrates, because the model is trained on and returns predictions for A domain/substrate pairs. The general applicability of this feature greatly depends on the quality of annotations for promiscuous A domains. While we aimed to reduce the number of false positives by excluding minor substrates from our dataset, generating less biased and higher quality data in the future will facilitate more reliable A domain promiscuity predictions.

### PARAS and PARASECT outperform existing predictors

To demonstrate the improvement of PARAS and PARASECT over other A domain selectivity predictors, we benchmarked both tools against NRPSPredictor2, AdenPredictor, and SANDPUMA. As SANDPUMA is an ensemble of multiple predictive algorithms, we also assessed these individually. In this comparison, we also included the deep learning tools NRPSTransformer60 and DeepAden61, which were published after initial deposition of our manuscript as preprint and developed in parallel. For testing, we used a bacterial (151 domains, collated by the authors of NRPSTransformer and curated for this work) and a fungal (130 domains) benchmarking set (https://zenodo.org/records/17404295), both consisting of A domains that none of the tools were trained on.

PARAS and/or PARASECT outperformed all other algorithms for both bacterial and fungal domains(Figure 4a,b). For fungal domains, this difference was substantial, with observed accuracies of ∼70% for PARAS and ∼66% for PARASECT compared to the next-best algorithm AdenPredictor (∼23%, MWU, p<1e-16; Figure 4a). Note that the poor performance of NRPSTransformer (∼13%) is expected, as it was not trained on any fungal A domains. For bacterial systems, PARAS, PARASECT, and NRPSTransformer all performed comparably, with PARAS performing best (∼77%), closely followed by PARASECT (∼75%) and NRPSTransformer (∼75%, MWU, p=1.2e-4 vs PARAS, p=5.9e-4 vs PARASECT; not significant; Figure 4a). Overall, we observe that model performance correlates strongly with the number of training points in the dataset (R2fungal=0.95, R2bacterial=0.80, Figure 4c). In contrast, model and featurisation choice are less important: NRPSTransformer, PARAS, and PARASECT, which use vastly different approaches, all perform comparably on bacterial data, and have very similar training set sizes. SANDPUMA (phylogeny ensemble model), NRPSPredictor2 (SVM), AdenPredictor (RF), and DeepAden (GNN), which were all trained on far smaller datasets, perform similarly to one another as well (Figure 4c). This demonstrates that, on A domain data, classical machine learning models such as random forests can perform just as well as if not better than deep learning models.

**Figure 4.**
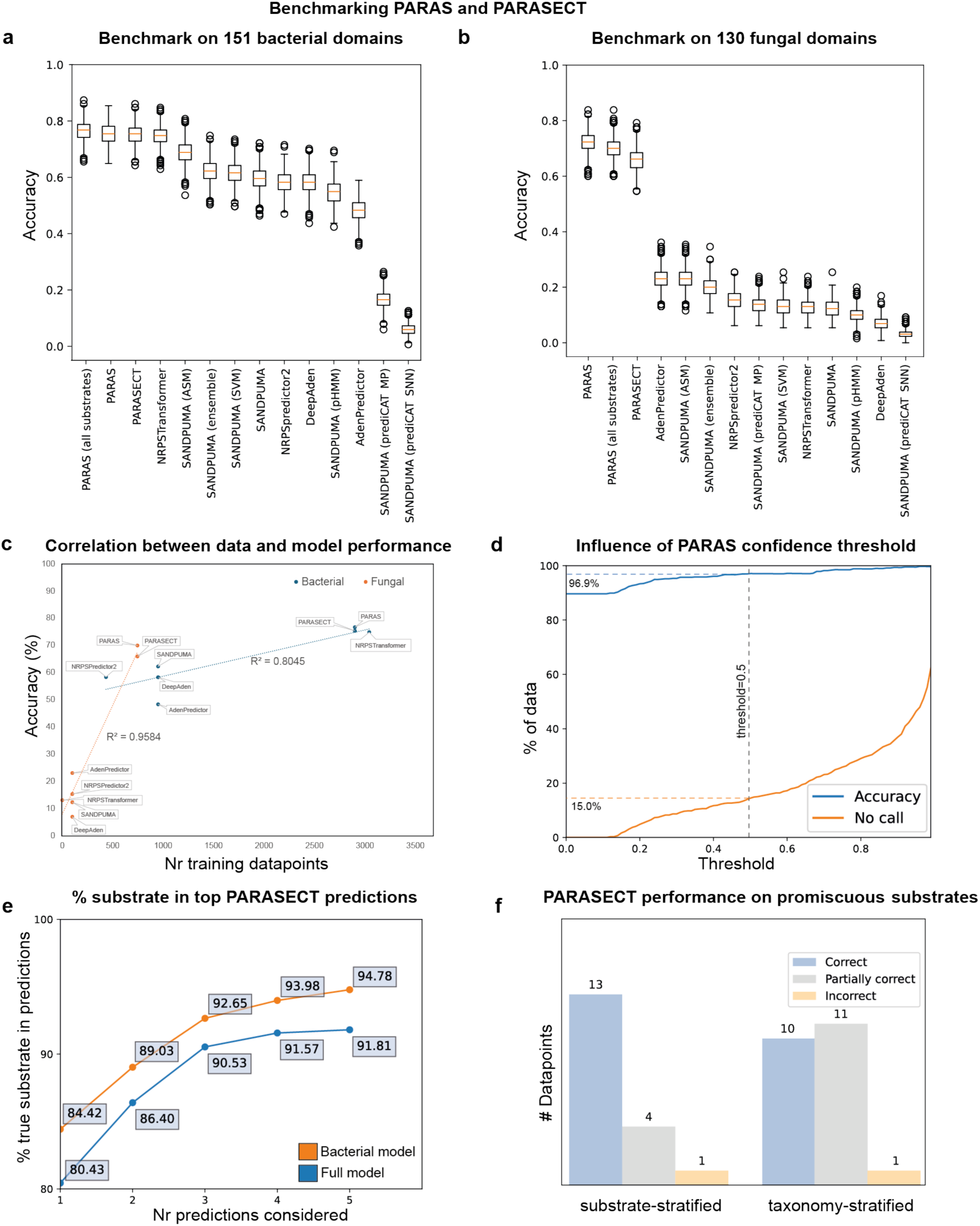
Benchmarking PARAS and PARASECT. **a.** Performance comparison on bacterial domains. **b.** Performance comparison on fungal domains. **c.** Correlation between tool accuracy on fungal and bacterial domains based on the number of fungal/bacterial domains in the dataset of each tool, showing that data and not model choice is the dominant factor in model performance. **d.** PARAS accuracy and number of data points for which no prediction was made across different confidence threshold cutoffs. **e.** % of times PARASECT trained on a taxonomy-stratified dataset predicts the correct substrate among its top 1-5 predictions. Orange: bacterial PARASECT model predictions on bacterial domains. Blue: full PARASECT model predictions on all domains. **f.** PARASECT performance on promiscuous substrates. For data points labelled as ‘correct’, the substrates selected by the A domain and the top x substrates predicted for that domain, where x is the number of substrates selected by that domain, match exactly. ‘Partially correct’ data points are A domains for which there is an overlap of at least one substrate between the actual x substrates and the top x predicted substrates. For data points labelled ‘incorrect’, there is no overlap between the actual substrates and the top x predicted substrates.

As prediction speed is also relevant, especially for integration into tools hosted by public web servers such as an-tiSMASH and for user convenience, we performed a speed benchmark on all A domain predictors. NRPSPredictor, PARAS and PARASECT, AdenPredictor, and NRPSTransformer were the fastest algorithms (Table S2), achieving speeds of 0.002-∼0.03s/domain. However, NRPSTransformer only achieves these speeds when run on a GPU from the command line. The speeds achieved on the NRPSTransformer web server are much lower, with the five A domains encoded by dptA (BGC0000336) taking ∼181s to complete, averaging ∼36s per domain. Additionally, due to their simple architecture, PARAS/PARASECT models take less than a minute to retrain.

While many current A domain specificity predictors enforce a cutoff to only return highly confident predictions, we chose not to implement such a cutoff for PARAS. As PARAS already performs very well at low confidence thresholds and many correct predictions would be missed when implementing a threshold (Figure 4c), we felt the community would benefit more from our raw model. Still, PARAS/PARASECT always return a confidence score alongside their predictions so that the user can consider this in the context of the biological system they are studying.

When querying PARASECT with sequences distantly related to the training data (e.g., from less-well-studied organisms), it is important to note that its feature to provide prediction probabilities for multiple possible substrates can come in very useful: in the taxonomy-stratified dataset, when the top two predictions are considered, the accuracy of the model improves by nearly 10% (Figure 4e).

However, PARASECT frequently gives high interaction scores to substrates structurally similar to the actual substrate, even when these other substrates are not selected by the A domain (Figure S8), leading to a slight drop in overall performance compared to PARAS (Figure 4a). At the same time, PARASECT does also allow correct identification of real substrate promiscuity of bacterial domains: out of 18 promiscuous A domains in our substrate-stratified test set, for 13 the top ‘x’ PARASECT predictions exactly matched the set of substrates incorporated by the A domain, where ‘x’ is the number of different substrates incorporated. For 4, there was at least 1 substrate overlap between the top ‘x’ PARASECT predictions and the actual substrate selectivities (Figure 4f).

We also assessed the substrate-specific prediction accuracy for each of the 37/38 substrates that PARAS and PARASECT were trained on. As our computational modelling and docking indicated that active site architectures of domains recognising large substrates may vary substantially across phylogenetic clades, we were particularly interested in performance differences for large substrates. Comparing the test sets obtained through substrate-based and taxonomy-based stratification (i.e., testing on sequences relatively closely versus distantly related to the training data), we indeed observed a large difference in performance across these two datasets for various large substrates, including l-Arg and l-Trp. While PARASECT correctly classified 71-79% of the data points for both substrates in the substrate-stratified test set (F1: 0.71/0.87, respectively), this dropped to 57-71% in the taxonomy-stratified test set (F1: 0.31/0.59, respectively; Figure S11, Table S7). Other hard-to-predict substrates were underrepresented ones (such as pipecolic acid, d-Ala, homoserine, and 2-aminoadipic acid) as well as l-Val and l-Ile, whose active sites frequently clade together. This illustrates the necessity of a large and diverse training set to cover as much phylogenetic breadth as possible. After model validation, cross-validation, and benchmarking, we used our full dataset to train four sequence-based models that we made available to the community on our web application (paras.bioinformatics.nl): a PARAS model and a PARASECT model trained on all 37/38 substrates for which at least 10 examples exist in our dataset; an additional PARAS model trained on all 3,653 A domains; and a PARASECT model trained on all 2,900 bacterial domains (30 substrates). These models are also available through direct outlinks in the version 8 release of antiSMASH.

To fully leverage the retrainability of PARAS/PARASECT, we have included a data submission portal on our web page, paras.bioinformatics.nl/data_annotation, where users can annotate new data points or correct existing ones. Entries are automatically detected from protein sequence and checked against a SQLite database of annotated A domain, protein, taxonomy, and substrate data (Table S4). Human error is minimised by providing PARAS predictions for each entry, enforcing literature referencing, and reviewer curation through GitHub. Users can optionally submit their ORCID ID, such that their contributions can be tracked. This facilitates rapid retraining of PARAS and PARASECT with the most recent A domain data, providing a continually maintained high-quality A domain dataset and frequently updated state-of-the-art models to the community.

### Characterisation of an unusual tryptophan-incorporating A domain in the tryptopeptin NRPS

To further validate PARAS and PARASECT, particularly in the context of A domains incorporating amino acids with large hydrophobic side chains, we selected tryptopeptin A^62^, a biosynthetically uncharacterised member of the peptidyl epoxyketone family of proteasome inhibitors with an unusual tryptophan-derived α,β-epoxyketone pharmacophore. We hypothesised that tryptopeptin is assembled following similar biosynthetic logic to eponemycin and TMC-86A^63,64^. Thus, an NRPS would catalyse the successive condensation of a 3-methylbutyryl thioester with L-Val, L-*allo*-Thr and L-Trp to give rise to an N-acyl-tripeptidyl thioester, followed by polyketide synthase-catalysed net elongation of the Trp residue with propanoate, and conversion of the resulting α-methyl-β-ketoacid to the corresponding epoxyketone by a flavin-dependent decarboxylative-desaturase / monooxygenase.

Using ClusterTools^65^, we searched the Prokaryotic RefSeq Representative Genomes repository at NCBI for gene clusters containing a homologue of the gene encoding the eponemycin epoxyketone synthase EpnF (protein ID AHB38508.1) and genes encoding at least one A, C, and ketosynthase domain. Among the hits, a BGC (GenBank accession: NZ_MAXF01000131) in the draft genome sequence of *Streptomyces sparsogenes* ATCC 25498 (DSM 40356) contained genes encoding a putative epoxyketone synthase (TtpC), a tri-modular NRPS (TtpD), a PKS module (TtpE) containing a putative C-methyl transferase domain, and a proteasome β-subunit (Figures 5a and 5b). Detailed manual annotation of the enzymes encoded by this BGC indicated that it likely directs the biosynthesis of tryptopeptin-like metabolite(s) because (i) the 9-residue active site signatures of the A domains in modules 1 and 2 of the NRPS in the BGC were predicted to select L-Val and L-Thr, respectively, and (ii) although the substrate of the A domain in module 3 of the NRPS could not be predicted from the 9-residue active site signature, the relative compactness and hydrophobicity of the residues suggested the preferred substrate is a large hydrophobic amino acid (Figure 5b).

**Figure 5.**
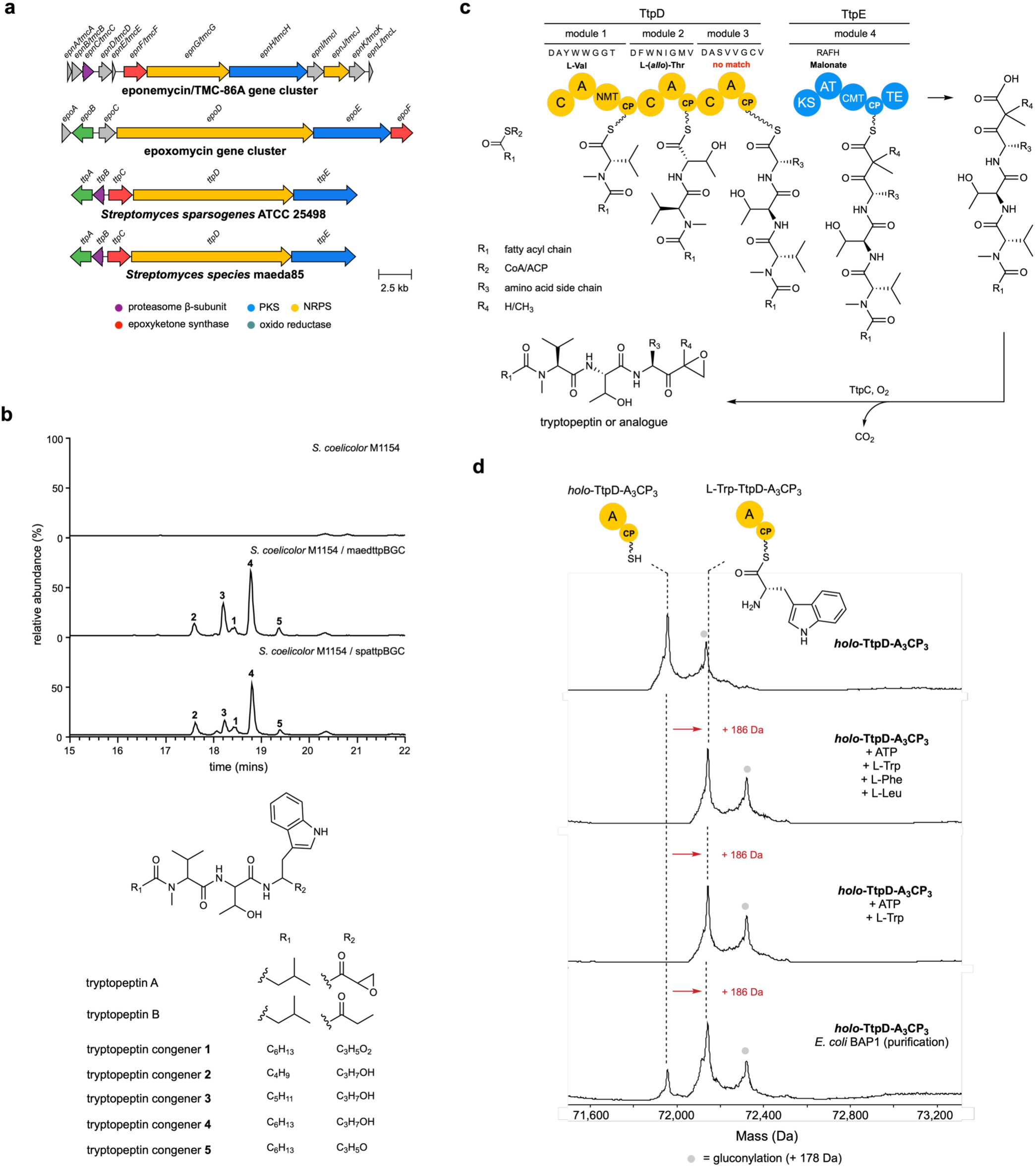
Experimental validation of the tryptopeptin biosynthetic gene cluster. **a.** Comparison of putative tryptopeptin BGCs in *Streptomyces sparsogenes* and *Streptomyces* sp. maeda85 with the eponemycin and epoxomycin BGCs. **b.** Proposed biosynthetic pathway for tryptopeptin A. **c.** Extracted ion chromatograms (EICs) from UHPLC-ESI-Q-TOF-MS analyses of culture extracts from *S. coelicolor* M1154, *S. coelicolor* M1154 containing pCAP1000maedttpBGC and pCAP1000spattpBGC corresponding to [M+H]^+^ for tryptopeptin-related metabolites 1, 2, 3, 4, and 5. The proposed planar structures are consistent with MS/MS analyses (Fig S11). **d.** Intact protein MS analysis of the purified *holo*-A-PCP di-domain from module 3 of TtpD incubated with an amino acid mixture, L-Trp, or loaded *in cellulo* using *E. coli* BAP1 cells, demonstrating that TtpD-A3 preferentially loads L-Trp *in vitro* and *in vivo*.

No tryptopeptin production was detected in *S. sparsogenes* DSM 40356. We therefore sequenced and assembled the genome of a previously identified tryptopeptin A producer *Streptomyces sp.* maeda85. Analysis of the genome sequence revealed a BGC (GenBank accession: PQ740899) with high similarity to the putative tryptopeptin BGC we identified in *S. sparsogenes* DSM 40356 (Figure 5a). UHPLC-ESI-Q-TOF-MS/MS analyses confirmed that *Streptomyces sp.* maeda85 produces tryptopeptin A (Figure S13).

To verify that the highly similar BGCs identified in the genomes of *S. sparsogenes* DSM 40356 and *Streptomyces* sp. Maeda85 direct the production of tryptopeptin, both were cloned from genomic DNA using our recently reported high efficiency yeast transformation-associated recombination capture vector pCAP1000^64^. The captured BGCs were transferred via conjugation from *E. coli* into *Streptomyces coelicolor* M1154^66^. Analyses of culture extracts from the trans-conjugants using UHPLC-ESI-Q-TOF-MS/MS identified five tryptopeptin-related metabolites, all containing an *N*-methyl-L-Val-L-(*allo*)-Thr-L-Trp core scaffold (Figure 5c; Figure S14, Supplementary Discussion).Notably, these metabolites were produced in similar relative quantities by strains expressing the two BGCs, neither of which directed tryptopeptin A production in *S. coelicolor*.

To confirm that the A domain in module 3 of TtpD incorporates L-tryptophan, we overproduced the corresponding A-PCP di-domain in *E. coli* BL21(DE3) and *E. coli* BAP1 as N-terminal His_6_ fusion proteins. These strains are expected to produce the *apo* and *holo* forms of the protein, respectively^67^. Production of the *apo-*A-PCP di-domain by *E. coli* BL21(DE3) was confirmed by UHPLC-ESI-Q-ToF-MS analysis of the purified protein (Figure S15, top row). The *apo* protein was converted to its *holo* form using Sfp and coenzyme A (Figure S15, second row) and the ability of the A domain to load L-Trp, L-Phe, L-Leu, L-Val, and L-His onto the downstream PCP domain was examined using UHPLC-ESI-Q-ToF-MS (Figure S15, rows 3-7). When the *holo*-A-PCP di-domain was incubated with the substrates in isolation, high levels of L-Trp and L-Phe loading were observed, along with some loading of L-Leu, very little loading of L-His and no loading of L-Val. However, when incubated with a mixture of L-Trp, L-Phe and L-Leu, the *holo*-A-PCP di-domain exclusively loaded L-Trp (Figure 5d), suggesting that this is the preferred substrate. We validated this by purifying the A-PCP di-domain from *E. coli* BAP1. UHPLC-ESI-Q-ToF-MS analysis showed this is a mixture of the unloaded and Trp-loaded *holo*-protein (Figure 5d), confirming L-Trp is the sole substrate loaded onto the PCP domain *in cellulo*.

While we could use the broad selectivity hypothesised by manual inspection of the active site of the A domain in module 3 of TtpD to link the BGC in both strains to production of tryptopeptin-related metabolites, this example demonstrates the need for selectivity predictors that can handle A domains with active site signatures that are divergent from previously characterised examples. Because PARAS and PARASECT were developed with these types of domains in mind, we were curious to see how our models performed on the A domain in module 3 of TtpD. We therefore predicted the substrate preference of the A domains in TtpD from *S. sparsogenes* DSM 40356 with PARAS and PARASECT, in addition to AdenPredictor, NRPSPredictor2 and SANDPUMA. PARAS and PARASECT both correctly predicted that this A domain incorporates L-Trp, while in contrast, AdenPredictor and NRPSPredictor2 were not able to identify L-Trp as the substrate, instead predicting L-Asp and L-Leu, respectively. The only other algorithm that correctly identified tryptophan was the SVM submodel of SANDPUMA, with the other submodels of this algorithm not able to give a confident prediction. Importantly, the A domain from module 3 of TtpD does not occur in the PARAS and PARASECT training sets. Indeed, PCA-based comparison of the AlphaFold model of this domain to models of other L-Trp-incorporating A domains shows that it is structurally distinct from domains in the training set, possibly indicating yet another binding mode for this substrate (Figure S16). This demonstrates the added value of PARAS and PARASECT over existing predictors and non-machine learning methods for the discovery and de-orphaning of natural products.

### Feature inference reveals residues involved in substrate recognition

In addition to being a suitable machine learning solution for small datasets, the random forest models used by PARAS and PARASECT have another advantage: their simple architecture based on combining multiple decision trees make their models interpretable. Therefore, we were able to use feature inference to gather clues about the structural basis for A domain substrate selection. We first looked at features that are used most frequently to make predictions in general (Figure 6a) and then calculated substrate-specific feature importances to assess whether there is a difference in coordinating residues between A domains that recognise different substrates (Figure 6b-f). Using our sequence-based PARAS model for feature inference, we visualised feature importance three-dimensionally by highlighting the residues in AlphaFold models of representative A domains with manually docked substrates.

**Figure 6.**
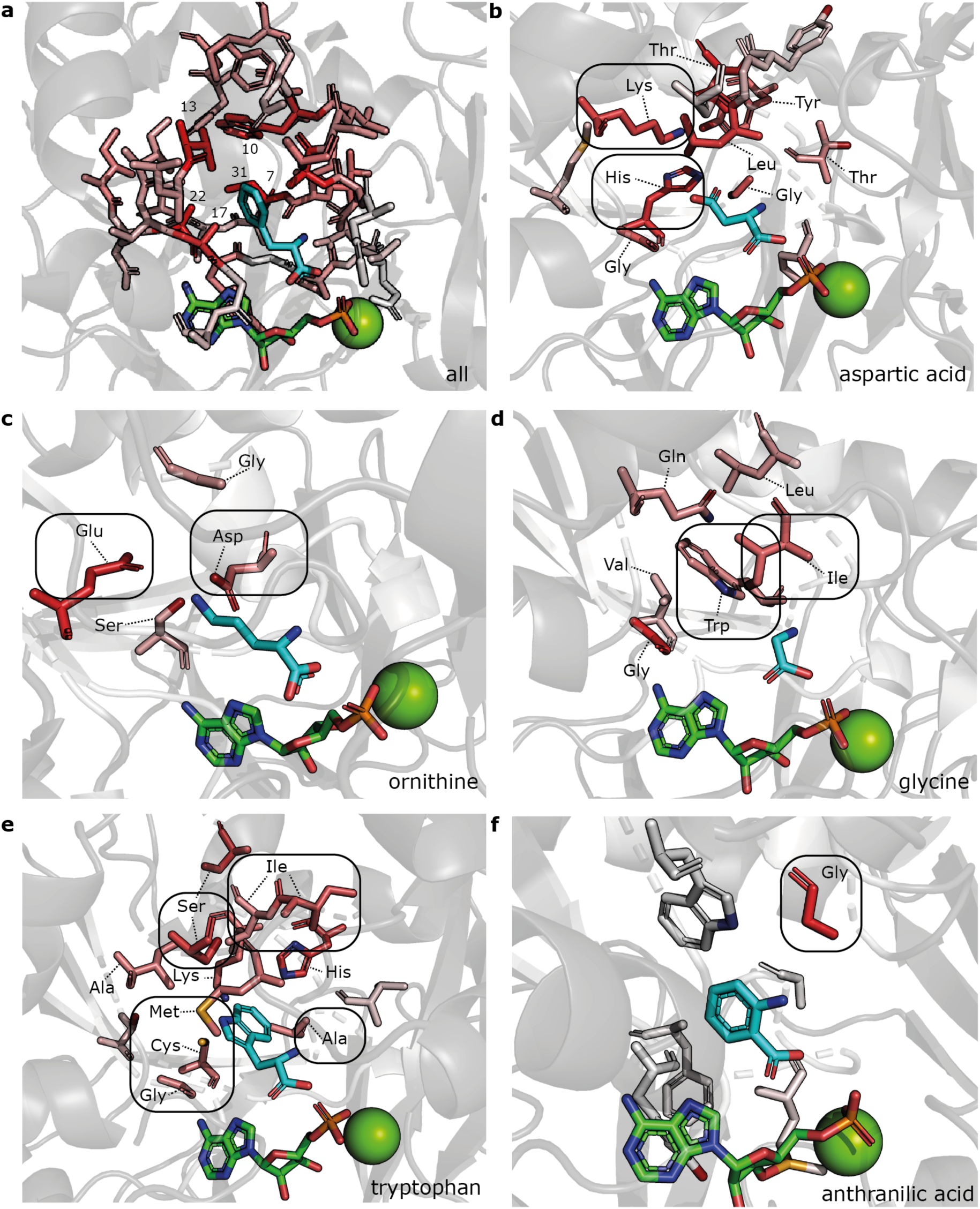
Three-dimensional visualisation of PARAS feature importance analysis. Feature importances are visualised on AlphaFold models of representative A domains. Residues with an importance greater than zero are shown. The redder the residue, the more informative the features that describe that residue. Substrate carbon atoms are shown in cyan; AMP carbon atoms and Mg^2+^ are shown in green. **a.** Overall perresidue feature importance. The six most informative residues are labelled with their extended active site residue number. **b.** Feature importance for aspartic acid-selecting A domains, with positively charged lysine and histidine residues indicated. **c.** Feature importance for ornithine-selecting A domains, with negatively charged aspartic acid and glutamic acid residues indicated. **d.** Feature importance for glycine-selecting A domains, with large hydrophobic isoleucine and tryptophan residues indicated. **e.** Feature importance for tryptophan-selecting A domains, with hydrophobic and aromatic residues indicated. **f.** Feature importance for anthranilic acid-selecting A domains, highlighting the highly conserved residue 6, which is an aspartic acid in most NRPS A domains.

Overall, we found that the residues close to the active site were the most informative (Figure 6a). The six most informative residues (residues 7, 10, 13, 17, 22, and 31 of the extended active site signature) are all also part of the 10 residue active site signature as defined by Stachelhaus *et al.*^11^, and all but one (residue 31) belong to the 9 residues described by Challis *et al.*^10^. These residues are positioned more closely to the active site pocket than other residues in our analysis, providing some evidence that the model is looking at biologically relevant features to make predictions. Still, the model uses many features derived from residues outside of the 10-residue active site signature, indicating that these residues are also informative. Possibly, the residues just outside the active site pocket may influence the position of residues that point inside of it. For example, an A domain that selects L-Glu may have aromatic ring stacking of one phenylalanine and two tryptophan residues. While only one of the tryptophan residues directly borders the active site pocket, all three residues are highly predictive for A domains that select glutamic acid (Figure S17). Also, residues not directly bordering the active site could carry a phylogenetic signal that the model uses for its predictions.

Substrate-specific feature importance analysis revealed that the residues that are important for making an A domain selectivity prediction depend on the substrate that an A domain recognises, suggesting that the residues involved in the active site are different for each specificity. Largely, the residues deemed important by the model are congruent with what one would expect biologically: basic lysine and histidine residues for aspartic acid (Figure 6b); negatively charged aspartic acid and glutamic acid residues for ornithine (Figure 6c); large hydrophobic residues restricting the size of the active site pocket for glycine (Figure 6d); small hydrophobic residues leaving open a large pocket for tryptophan (Figure 6e); and a non-aspartic acid residue at the highly conserved position 6 for anthranilic acid, which lacks an amino group in its backbone and therefore does not require aspartic acid for substrate stabilisation (Figure 6f).

Figures showing feature importances for all 34 different substrates can be found in the Supplementary Material (Figures S17j-S22). While some of the informative residues overlap between substrates, many do not: for example, the positive charge required in active site pockets selecting glutamic acid and aspartic acid is conferred by a different set of residues for each (Figure S17). As such, simple sequence similarity measures against a one-size-fits-all ‘specificity-conferring code’ as once hypothesised cannot accurately describe or predict A domain selectivity, demonstrating why machine learning is ideal for solving this challenge.

Our feature importance analysis also shed light on limitations of previous algorithms in predicting large substrates: to make predictions for A domains recognising large substrates, PARAS uses more features on average than for most other substrates (Figure 6e, Figure S19). The same was true for smaller substrates which do not clade together. This supports our hypothesis that A domain specificities have evolved multiple times, and explains why our expanded dataset greatly improved model performance for these substrates.

## DISCUSSION

A domain specificity prediction has historically been challenging for those A domains that do not clade according to the substrates that they recognise. In this work, we used large-scale analysis of over 3000 AlphaFold models to illustrate that this challenge is especially prominent for large substrates, likely due to a higher degree of rotational freedom in their respective A domain active sites.

To tackle this issue, we leveraged a larger and better-curated dataset, evaluating structure-based approaches and introducing substrate featurisation methods to build PARAS and PARASECT: two novel A domain selectivity predictors which offer substantial improvements compared to state-of-the-art tools, showing a performance increase of 14% for bacterial domains and 49% for fungal domains. A cross-comparison of our new models and currently available tools, which leverage a wide variety of featurisation methods and machine learning algorithms, respectively, clearly demonstrates that data are crucial. AdenPredictor and our best performing PARAS model use identical featurisation methods and highly similar machine learning models to make predictions, yet PARAS outperformed AdenPredictor by ∼28-49% on independent holdout data. By far the most substantial difference is the increased size and quality of the PARAS dataset, which is over three times as large as the AdenPredictor dataset and was extensively curated prior to model training. As such, we recommend focusing machine learning efforts in low-data regimes on data collation, acquisition, and curation, whenever this is feasible. This is why we launched a SQLite database for A domain data, alongside a A domain data annotation and correction portal. Combined with the easy retrainability of random forest models, this ensures that the A domain models available to the community are based on the latest available training data.

While the addition of substrate features to PARASECT did not lead to improved model performance, it has the benefit that the model accounts for substrate similarity in its decisions, leading to more informative secondary and tertiary predictions. This makes PARASECT especially useful for obtaining substrate approximations for A domains that are phylogenetically distant from A domains in the training set. Structure-based feature extraction methods have similar use cases, showing improved A domain active site extraction for clades of A domains underrepresented in the training data.

By combining computational and experimental methods, we identified a BGC that directs the production of tryptopeptin-related metabolites in two *Streptomyces* strains. Importantly, the tryptopeptin A scaffold, which includes a critical tryptophan substrate in position 3, was only accurately predicted by PARAS and PARASECT, and not by SANDPUMA, AdenPredictor, and NRPSPredictor. This, together with feature inference analysis of our models that shows a greater quantity of important features for large substrates, demonstrates that our models successfully address the challenge of large substrates in A domain selectivity prediction. Also, it is noteworthy that the L-Trp-incorporating A domain of TtpD was structurally different from all other L-Trp-incorporating A domains in our dataset, indicating that A domain diversity remains underexplored, even in well-represented phyla like the Actinomycetota.

To our knowledge, heterologous expression of A domains in *E. coli* BAP1 cells followed by MS to investigate *in vivo* A domain selectivity is a novel and important innovation over existing A domain assaying methods. Critically, this method assays both the adenylation and thiolation half-reactions that are required for A domain substrate selection, in stark contrast to the ATP-PPi exchange assays that are still a popular method for substrate determination. In addition, the bias that is introduced by researchers choosing which substrates to assay is removed; instead, the assayed A domain has access to the full diversity of substrates that are naturally available *in cellulo*, providing a means to perform large-scale competition assays including many substrates at once. Assaying A domains for substrates that are not naturally available to *E. coli* BAP1 could be achieved by altering the growth medium in which the cells are grown. We therefore recommend this approach to those in the natural products community studying NRPS systems. This will generate high-quality data to be integrated into future iterations of PARAS and particularly PARASECT, which can leverage these data to distinguish between substrate-tolerant A domains and truly promiscuous A domains that incorporate multiple substrates *in vivo*. Combined with feature inference approaches such as those demonstrated here, this can provide critical insights into molecular determinants of A domain promiscuity.

Already, PARAS and PARASECT have been used to discover natural products in the form of novel tridecaptin derivatives^68^. There, PARAS was used to identify two novel lipopeptide antibiotics, tridecaptin A_5_ and tridecaptin D, correctly predicting the substrates of all A domains in the NRPS enzymes TriD and TriE, while NRPSPredictor2 and AdenPredictor made various errors (Table S3). In addition, PARAS predictions and confidence scores have already been leveraged to compare and align peptaibol synthase architectures in fungi^51^, and aided in the discovery of the novel bacterial glycotetrapeptide biffamycin A^69^. By making both models freely available on an easily navigable web application, and by integrating an outlink to PARAS into antiSMASH 8.0, we hope that these models will be widely used by the natural products community to deorphan and discover novel compounds.

## EXPERIMENTAL SECTION

### Computational methods

#### Data collection and curation

To train PARAS and PARASECT, we collated a dataset of 3,653 unique A domains from different sources: the NRPSPredictor2 dataset^22^, the SANDPUMA dataset^25^, MIBiG 3.0^28^, MIBiG 4.0^29^, and recent literature^30–51^. A domains were labelled by their GenBank, Ref-Seq^70,71^ or UniProt protein ID^72^ and a number corresponding to the index of the A domain in the NRPS enzyme. For all A domains, we re-downloaded the sequence of their protein from GenBank, JGI, or their primary literature source, and extracted the sequences of N-terminal and C-terminal A subdomains with HMMer3^73^, using the AMP-binding (PF00501.29) and AMP-binding_C (PF13193.7) Pfam pHMMs^74^, respectively. We then deduplicated our dataset through pairwise identity searches, allowing for truncations on either end of the A domain (minimum overlap length: 0.8 x shortest sequence length). For each A domain, we kept all protein labels.

As errors are unavoidable in human-collated datasets, we set out to curate our dataset in three steps. First, during deduplication, we flagged any domains with identical sequences but different selectivities and consulted the literature to fix their annotations. We repeated this process for A domains with identical 10 amino acid active site signatures, asserting that any differences in specificity were backed up by literature, fixing annotations where they were incorrect or omitting data points when we suspected experimental data to be unreliable. For instance, the active site signatures reported in literasture for the first A domains in the proteins AAZ03554.1 and AAZ03552.1 were drastically different from the active site signatures we extracted^75^. Finally, we phylogenetically grouped the A domains based on their 34 amino acid extended active site signature with FastTree^76^ and similarly verified the specificity of obvious outliers.

#### Structural modelling

Prior to structural modelling, we aligned all adenylation domains with MAFFT v7.508^77^ (default settings). We used this alignment and the reference crystal structure of the GrsA A domain (1AMU^56^) to extract the N-terminal and C-terminal subdomains for each A domain. We then obtained predicted structures for each A subdomain domain using ColabFold^78^ in batch mode (default settings) without side chain relaxation. We only acquired structures of the C-terminal subdomain for 2617 domains in our dataset. As the C-terminal sub-domain does not play a large role in substrate recognition, with only the highly conserved K517 residue involved in stabilising the amino acid backbone of the substrate^10,11,22^, we decided to generate full structural models by combining the AlphaFold model of each native N-terminal subdomain with the C-terminal subdomain of 1AMU. We then aligned all full structure models to 1AMU in PyMol^79^ (open-source, v2.5.0) and saved all models in identical orientations in Protein Data Bank^80^ (PDB) format. Finally, we copied the Mg^2+^ and AMP cofactors from 1AMU to the PDB files of all A domains to reflect the state of the active site pocket without the amino acid substrate bound.

#### Sequence-based domain feature extraction

To obtain sequence-based features for our models, we first extracted the 34 residues that correspond to the 34 residues within 8Å of the phenylalanine substrate in the A domain of GrsA, as also done by the authors of NRPSPredictor2^22^. We tested three approaches: two used Muscle^81^ (v3.8.1551) profile alignments, using the sequence of the GrsA (BAA00406.1) A domain as a reference. As a guide alignment, we used either sequence-based or structure-based sequence alignments of a subset of our dataset (923 A domains). We obtained the sequence-based guide alignment with Muscle (v3.8.1551), and the structure-based guide alignment by modelling the N-terminal A subdomains with Modeller^82^ (v10.0, default: 250 models per sequence, no loop refinement) and aligning the resulting homology models with Caretta-shape^83^ (v1.0, default settings). As sequence extraction time scales exponentially with the size of the guide alignment, we chose to use a subset of our data rather than the full dataset. The final sequence extraction approach, and the one we ended up using for our published models, uses a previously used NRPS-specific AMP-binding domain pHMM^21^ with HMMer2^73^ (v.2.3.2). As some fungal domains are not detected by this pHMM, the final models also run more general AMP-binding HMMs (PF00501.29, PF13193.7) from PFAM^74^ for A domain detection, using structure-based profile alignment with Muscle for signature extraction by default when no matching NRPSPredictor2 pHMM hits are found.

Each of the resulting 34 amino acid residues was featurised as a list of 15 physicochemical properties describing hydrophobicity (WOLS870101^84^, NEU1, NEU2, NEU3 (Neumaier *et al.* accessed 2024), size (WOLS870102^84^, TSAJ990101^85^), electron state (WOLS870103^84^), hydrogen bond donors (FAUJ880109^86^), polarity (GRAR740102^87^, RADA880108 ^88^, ZIMJ680103 ^89^, occurrence in alpha-helices (CHOP780201), beta-sheets (CHOP780202) and beta-turns (CHOP780203)^90^, and isoelectric point (ZIMJ680104 ^89^)^91^, as previously described for NRPSPredictor and NRPSPredictor2^21,22^. These physicochemical features were then concatenated into feature vectors of length 510 (34 x 15; Figure 3b). Physicochemical properties and guide alignments can be found at https://github.com/BThe-DragonMaster/parasect/tree/master/src/parasect/data.

#### Comparing structure-based and sequence-based guide alignments

To compare structure-based and sequence-based guide alignments for feature extraction, we first split a subset of 923 domains into ten cross-validation sets, creating structure-based alignments (Caretta-shape^83^ v1.0) and sequence-based alignments (Muscle^81^ v3.8.1551) for each training set, stratifying on substrate class and using only the first reported substrate as response. Then, we featurised the 34 active site residues that we extracted from the structure-based and sequence-based alignments (as described above) and trained random forest models (scikit-learn^92^ v1.1.3, default settings) on those same cross-validation sets and compared the test accuracy of the resulting two models. To ensure fair comparison, sequence-informed and structure-informed models were trained on the same data points using the same random seed for each cross-validation set.

#### Structure-based domain feature extraction

To convert our AlphaFold models into feature lists, we first placed a cubic voxel grid of 20×20×20 voxels centred around the position of the β-carbon of the phenylalanine substrate in the 1AMU reference structure. Next, for each A domain, we determined how many atoms overlapped with each voxel, recording counts for 7 atom types separately: aromatic, hydrophobic, charged, and non-charged oxygens, charged and non-charged nitrogens, and sulphurs. We then concatenated these 7 numbers for all 8000 voxels into a single vector of length 56000 representing the A domain active site. As this is too large a number of features to work with considering our dataset size, we decided to reduce dimensionality through principal component analysis (PCA)^93^. To ensure that the resulting principal components captured variation between the active sites of highly similar substrates, we first assigned each data point to one of 9 substrate categories: aromatic amino acids, aromatic acids, Asx/Glx, cyclic aliphatic amino acids, substrates with nitrogen-containing side chains, phenylglycines, small amino acids, small hydrophobic amino acids, and small polar amino acids (Figure 3b). Data points that recognise substrates belonging to different groups were assigned to multiple groups. We only included substrates for which we have 11 or more examples in our dataset (Figure 2b). Then, we performed PCA on each of the 9 substrate groups as well as the full dataset using scikit-learn^92^ (v1.2.0) and stored the transformative models. For each PCA, we made a scree plot and determined the optimal number of components by using the kneedle package^94^ (v0.8.3) to automatically locate the knee in the plot. This yielded between 5-8 PCs for each substrate group and 11 PCs for the full dataset, cumulating to 68 PCs in total. We then applied each of the 10 transformative models on the full dataset to obtain all 68 PCs for all data points. The resulting PCs can be found in the ‘structure data’ folder of our online GitHub repository. These 68 PCs were consequently used as features for training PARAS and PARASECT (Figure 3b).

#### ESM2 embeddings

We generated ESM2 embeddings using the fair-esm Python implementation (v2.0.0, model: esm2_t33_650M_UR50D) on each complete A domain sequence. We extracted residue-specific embeddings (length 1280) for each residue of the 34-residue extended active site signature. As the resulting feature space (43520 features) was too large for training on directly, we reduced dimensionality with a PCA (scikit-learn v1.2.0), keeping the top 100 components as features for training.

##### Substrate feature extraction

As PARASECT predicts interaction between an A domain and a substrate based on features of both the domain and the substrate’s molecular structure, we also needed a way to featurise molecules. To this purpose, we collected isomeric SMILES strings^59^ for each substrate and used PIKAChU^95^ to extract ECFP-4 molecular fingerprints^55^ of length 1024 for each compound. These vectors were directly used for model training (Figure 3c, d).

#### Data stratification

Prior to model training, we stratified our data in two different ways to ensure we could properly assess how well our model will perform on unseen sequences afterwards. One split was done based on taxonomy such that each substrate was represented in both the training and test set with at least one example; and the other split was based on substrate class alone.

To split based on taxonomy, we obtained a taxonomic assignment for each A domain, either by fetching taxonomy from the Uniprot or GenPept ID, or from literature. A domains of unknown origin (from metagenomes or environmental DNA) were annotated as ‘None’. . We then assigned all A domains of the same taxonomic family to either the train or test set, first building coverage for all substrates (n >= 10) in both the train and test set (at least one example in each), then assigning families to train or test one at a time such that the ratio of train to test approximated 3:1 as closely as possible for each individual substrate. For PARAS train-test splits, we considered only the first substrate; for PARASECT train-test splits, we considered all substrates (n >=10). We later used this train-test split to assess how well our models are likely to perform on A domains that are phylogenetically very different from A domains in our training set.

To stratify based on substrate for PARASECT, we used an iterative stratification method that is suitable for multilabel datasets^96^. This strategy assures that data points are divided across train and test sets such that each substrate class is proportionally represented. We used the MultilabelStratifiedShuffleSplit module from the iterstrat package (v0.1.6; https://pypi.org/project/iterative-stratification/) to implement iterative stratification in Python, taking into consideration only those substrates for which there were at least 10 examples in our dataset. For PARAS, we stratified on the first-listed substrate using the train_test_split function in scikit-learn (v1.2.0).

#### Model training and evaluation

Models were trained and evaluated in four steps. 1) we selected the best model parameters and features through 3-fold cross-validation on the training sets; 2) we assessed overall performance and substrate-specific performance on the hold-out test sets; 3) we trained models with the best-performing parameters from step 2 on a) all data for bacterial benchmarking, and b) all data without the fungal A domains from the hold-out test set for fungal benchmarking to compare performance between PARAS/PARASECT and other A domain predictors; and 4) we trained our final models on all fungal data and all bacterial data minus the bacterial benchmarking set for publication. To enable meaningful validation on each substrate, we only included A domains recognising 37 common substrates for PARAS, and 38 for PARASECT (cutoff: at least 10 occurrences in the dataset, considering only the first label for PARAS and all labels for PARASECT) for steps 1 and 2. For steps 3 and 4, PARAS was trained on all 278 substrates. We used the StratifiedKFold module from the scikit-learn package (v.1.2.0; PARAS) and the MultiLabelStratifiedKFold module from the iterstrat package (v0.1.6; PARASECT) to perform 3-fold cross-validation on our training sets to assess intermediate model performance and tune model parameters. Both PARAS and PARASECT are random forest models implemented with the RandomForestClassifier module from scikit-learn^92^ (v1.2.0). For each train-test split, we trained four models for both PARAS and PARASECT, each using different approaches to featurising A domains: sequence features only, structure features only, a combination of both, or ESM2 features. As parameter tuning barely affected model performance, we decided to train all our models with 1000 trees, using default settings otherwise.

We trained PARAS as a single-label classifier, choosing the first listed label (which corresponds to the predominant substrate) as response for training purposes. PARAS output is therefore always a single substrate (Figure 3d). For assessing model performance, we designated a prediction as correct if the predicted substrate was among the substrates listed for that domain. We used balanced sampling to account for the differences in substrate counts for each class and prevent the model from prioritising the correct prediction of overrepresented substrates.

In contrast, PARASECT was trained on domain-substrate pairs, with a floating-point number as response, representing the probability of interaction between the domain and the substrate (Figure 3d). As this leads to highly imbalanced datasets, with a ratio of positive to negative data points around 1:30, we randomly under-sampled our data to achieve a 1:1 ratio using the RandomUnderSampler module from imblearn^97^ (v0.10.1). Model performance was subsequently assessed in two different ways: by looking at metrics that are commonly used for evaluating binary data, including precision, recall, and F1-score; and by determining if the substrate with the highest probability of interaction for a domain corresponds to one of the substrates that domain recognises. We additionally estimated a ‘cut-off’ interaction probability by obtaining and comparing mean and median interaction probabilities for true positives and false positives. For both PARAS and PARASECT, we also gauged model performance for each different substrate class.

Note that, for the comparison of sequence-based and structure-based featurisation methods, a dataset of only 3,254 data points was used.

#### Benchmarking

To benchmark PARAS and PARASECT against current state-of-the-art A domain predictors, we compared their performance against AdenPredictor^24^, SANDPUMA^25^, NRPSPredictor2^22^, DeepAden^61^, and NRPSTransformer^60^. To this purpose, we ran these predictors on a bacterial benchmarking set of 151 A domains published by the authors of NRPSTransformer, and a fungal benchmarking set of 130 A domains annotated by us. For a fair comparison, we trained separate benchmarking models for PARAS and PARASECT leaving out these benchmarking sequences.

If none of the true substrates are represented in the tool’s training set, the prediction was labeled “not in model.”. If no prediction is returned for a data point (no A domain was detected, or the confidence falls below the tool’s threshold), the prediction was labelled as “no call”. All other predictions were either labelled as “correct” or “incorrect”. In particular, for SANDPUMA, NRPSPredictor2, and AdenPredictor, which condense multiple labels into a single prediction, a prediction was labelled as “correct” if there was overlap between the true substrates of an A domain and the predicted substrates. For DeepAden, NRPSTransformer, PARAS and PARASECT, a prediction was only labelled “correct” if the top substrate was among the true substrates of an A domain.

To statistically compare the performance of different tools, we first generated a distribution of accuracy rates using 1000 bootstrap iterations. We then used the Kruskal-Wallis test to determine if there was a significant difference between the accuracy rates for at least two tools. When this was the case, we followed up with pairwise Mann-Whitney U tests to determine for each pair which tool significantly outperformed the other and whether the difference was significant.

For the speed benchmark, we used 100 sequences from the bacterial benchmark dataset. For fair comparison, we left out sequences for which no ‘correct’ or ‘incorrect’ prediction was given by some tools, as it may take no or little time, or additional computational time, for “no call” and “not in model” type inputs. For DeepAden and SANDPUMA, which are substantially slower than all other tools, we only speed-benchmarked on 1, 50, and 100 sequences. For other tools, we speed benchmarked on 1, 5, 10, 25, 50, 75, and 100 sequences. Time cost was recorded as the “real” time calculated by the “time” function in Shell. We repeated the speed benchmark 5 times for NRPSTransformer and 4 times for other tools, and removed one outlier for each to reduce random noise. We fitted the average of cost time by linear function and removed two outliers (AdenPredictor in 1 sequence and PARAS with “all substrates” mode in 25 sequences) in the fitting manually. Setup time (intercept) and per-domain computational cost (slope) were determined from the linear functions.

For NRPSTransformer, benchmarking and speed assessment were performed on a GPU server with 754G memory, 128 CPU (INTEL(R) XEON(R) GOLD 6548N) and NVIDIA L40S 48GB (2x). For other tools, benchmarking and speed assessment were performed on a CPU server with 3T memory, 256 CPU (AMD EPYC 9534 64-Core Processor) and NVIDIA L40S 48GB (2x).

Benchmarking sets are available at https://zenodo.org/records/17404295.

#### Structure modelling and substrate docking

To obtain structural models for the tryptophan-recognising A domains of Qui18-A1 and BreC-A3, we separately modelled their N-terminal and C-terminal subdomains with AlphaFold with amber relaxation^78,98^ (Google Colab v1.3.0, default settings). We then aligned both subdomains to the 1AMU reference structure to obtain full A domain models.

Next, we performed two dockings for each A domain: one with two separate ligands: tryptophan (PDB:TRP) and AMP; and the other with a reaction intermediate: tryptophanyl-adenylate (PDB:TYM). First, we prepared the ligands with the SDMolSupplier module of RDKit^99^ (v2022.03.5) by moving them into the rough vicinity of the active site, and subsequently converted them to .pdbqt format with the Meeko package (v0.2; https://pypi.org/project/meeko/). We converted the A domains to .pdbqt format with MGLTools. We manually added the Mg^2+^ atom into the .pdbqt files after. Finally, we performed the docking with Autodock Vina^100^ (v1.2.3; exhaustiveness: 64; number of poses: 40, Vina scoring function), saving the best 20 poses.

#### Feature inference

We performed two types of feature inference on our models: one that looks at overall feature importance in the entire model (PARAS and PARASECT); and one that assesses substrate-specific feature inference (PARAS only). For the former, feature importances were directly extracted from the RandomForestClassifier instances, and summed across all features describing the same residue to yield a measure of residue-level importance (Figure 6A). For substrate-specific feature inference, we iterated over the nodes of each individual tree in the forest, and determined the information gain per node for each substrate that was seen by that node. Information gain is defined as:

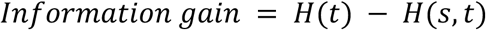

Where 𝐻(𝑡) is the entropy of the parent node, and 𝐻(𝑠, 𝑡) is the average entropy of the two child nodes. Entropy is calculated as follows:

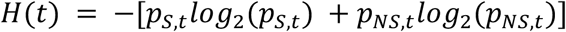

Where the probability of selecting substrate ‘S’ at node t, 𝑝_*s,t*_ = 𝑛(𝑡, 𝑆) / 𝑛(𝑡); and the probability of not selecting substrate ‘S’ at node t, 𝑝_*NS,t*_ = 𝑛(𝑡, 𝑁𝑆) / 𝑛(𝑡), where 𝑛(𝑡) is the total number of domains seen by node t, 𝑛(𝑡, 𝑆) is the number of domains recognising substrate S seen node t, and 𝑛(𝑡, 𝑁𝑆) is the number of domains not recognising substrate S seen by node t.

The average entropy of the two child nodes, 𝐻(𝑠, 𝑡), is given by:

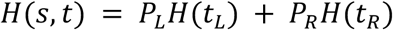

Where 𝐻(𝑡_*L*_) and 𝐻(𝑡_*R*_) are the entropy of the left and right child nodes of parent node t, calculated in the same way as 𝐻(𝑡); the probability of a domain at the left child node 𝑃_%_ = 𝑛(𝑡_*L*_)/𝑛(𝑡); and the probability of a domain at the right child node 𝑃_*L*_ = 𝑛(𝑡_*R*_)/𝑛(𝑡), where 𝑛(𝑡_*L*_) and 𝑛(𝑡_*R*_) are the number of domains seen by the left and right child nodes of node t, respectively.

As nodes are split on a single feature, information gain is a measure of entropy decrease for a specific substrate when the data is split on that feature. As such, the higher the information gain for a node, the more important the feature linked to that node.

To obtain an average substrate-specific information gain per feature, we can calculate the average gain of a feature across all its occurrences in the forest. An artefact of this method is that the information gain of features that are typically only used a few times in the forest but drastically decrease entropy when they occur is disproportionally inflated. This was especially problematic for features resolving leaf nodes for substrates that do not clade well together, such as tryptophan and alanine. Therefore, for each substrate we set the information gain for features that were used throughout the forest fewer than x times to 0, where we varied x from 0 to 500 at intervals of 50. We chose inclusion thresholds for each substrate individually, typically selecting higher inclusion thresholds for substrates whose recognising domains do not form monophyletic clades. Then, we obtained per-residue information gain by averaging across the information gain for all 15 features, Finally, we normalised the residue-level information gain across all substrates to highlight the differential contribution of certain residues to classifying specific substrates, and automatically visualised normalised per-residue information gain in Pymol^79^ for each substrate (Figure 6B-D, Figures S15-S20).

#### SQL database

We stored our dataset of 3,653 A domains in an SQLite3 database, available at https://github.com/BTheDragonMaster/para-sect/blob/master/app/src/server/parasect.db. This database stores domain, protein, substrate, and taxonomy data (Figure S12). The database is used by our webpage submission portal to check for duplicate user-submitted domain, substrate, and protein entries. Our GitHub contains Python scripts to add curated user entries to the database, and to train PARAS and PARASECT models directly from the database, facilitating easy retraining of both models.

##### Web portal development

To facilitate and promote the usage of PARAS and PARASECT, we have developed a web-based graphical user interface (v2.0.0). The web portal’s frontend is developed using JavaScript React (v18.1.0), which communicates with a Python Flask (v3.0.2) backend application. The web portal can be accessed at https://paras.bioinformatics.nl/. The user is able to upload a file containing protein sequences in FASTA format or a GenBank file, or input protein sequences in FASTA format directly. Settings for the user to customise are the model to use (PARAS substrate specificity prediction for all substrates, PARAS substrate specificity prediction for common substrates, PARASECT trained on all domains, PARASECT trained on bacterial domains only) and if to use profile-guided structure alignment for active site extraction. Additionally, if users pick PARASECT they are able to upload their own list of substrates to predict substrate specificity for. A short guide accompanied by screenshots from the v1.0.0 version of the web application can be found in Table S4 in the Supplementary Information.

The web portal also contains a data submission portal (https://paras.bioinformatics.nl/data_annotation), which relies on an SQLite3 database in the backend implemented using SQLAlchemy (v2.0.41)^101^.

Upon uploading one or more protein sequences through file upload (.gbk or .fasta), pasting fasta sequences directly, or pasting one or more NCBI GenPept IDs, the portal extracts all adenylation domains from the sequences. Sequences that already occur in the database (exact matches, or terminal overlaps of length 0.8 times the shortest sequence) are pre-annotated with their known substrates, and can be corrected. For new sequences, the PARAS all substrates model is run. Confident predictions should limit user error.

For both corrections and new annotations, users can annotate the domain with one or more substrates. These can either be selected from a drop-down list of known substrates, or input as a new substrate with a substrate name and a substrate SMILES, which are both checked against the PARASECT database to prevent duplicates (case-insensitive string matching for the substrate name; chemical fingerprints with PIKAChU^95^ for substrate matching. When one or more A domains have been annotated, the user can click ‘Submit’, upload literature references and optionally their ORCID ID, and confirm their submission, which will automate the creation of a GitHub issue. Entries in GitHub issues can subsequently be automatically downloaded and added to the database using scripts in our repository.

#### Experimental methods

##### Bacterial Strains, Plasmids, and Culture Conditions

*Streptomyces sparsogenes* DSM 40356, *Streptomyces species* maeda85, *Streptomyces coelicolor* M1154 and its mutants were grown on either TSB medium (Becton Dickinson) or SFM medium at 30°C for routine cultivation. For tryptopeptin production, 20µl spore suspension was inoculated on SY agar medium (starch 2.4%, glucose 0.1%, peptone 0.3%, beef extract 0.3%, yeast extract 0.5%, calcium carbonate 0.4%) and incubated at 30°C for 7 days.

Saccharomyces cerevisiae VL6−48N was grown on YPD medium (glucose 2%, yeast extract 1%, peptone 2%) supplemented with 100 mg/L adenine at 30 °C. For TAR cloning, tryptophan-deficient medium (sorbitol 18.2%, glucose 2.2%, yeast nitrogen base without amino acid and ammonium sulfate 0.17%, yeast synthetic dropout medium supplements without tryptophan 0.19%, ammonium sulfate 0.5%, and agar 2%) was used with 5-fluoroorotic acid (1mg/ml) (12).

#### TAR cloning and Streptomyces/*E. coli* conjugation

To construct capture vectors for cloning the two putative tryptopeptin BGCs, paired homologous arms were amplified from genomic DNA of corresponding microorganism with 20-30 bp primer-introduced overhangs for GeneArt™ Seamless Cloning and assembled into SpeI/XhoI-digested pCAP1000, together with the counterse-lectable cassette *pADH*-*URA3* amplified from pCAP1000 which is placed between the paired homologous arms. The integrity of resulting plasmids pCAP1000maedttp and pCAP1000spattp was confirmed by restriction digestion and sequencing.

To produce genomic DNA fragments containing tryptopeptin BGC region for TAR cloning, genomic DNA isolated from original hosts was digested with restriction enzymes that has at least one cutting site outside of target region (XbaI for *Streptomyces* sp. maeda85 and BswI for *Streptomyces sparsogenes*) and purified by ethanol precipitation. 0.5-1.5 µg of digested genomic DNA and 0.05-0.08 µg of the corresponding capture vector linearized by PmeI were used for yeast spheroplast transformation carried out using a previously reported protocol. The resulting yeast colonies were picked and screened by PCR (Table S5) using a previously reported protocol with primers designed to amplify fragments from the middle and each end of the target BGC^102^.

Plasmids were isolated from PCR-positive yeast colonies and used to transform *E. coli* Top10 for propagation. After verification by restriction digestion (XhoI and PstI for pCAP1000spattpBGC, SrfI and PstI for pCAP1000maedattpBGC), plasmids harboring tryptopeptin BGC were introduced into *S. coelicolor* M1154 by triparental mating from *E. coli* ET12567 with the helper strain *E. coli* ET12567/pUB307 using a standard protocol. Kanamycin resistance was used for exconjugant screening.

#### Growth of *Streptomyces* strains and analysis of epoxyketone production

After incubation, the agar was cut into small chunks and transferred into glass vials before adding 15 mL of ethyl acetate to each vial. After 2h, the organic extracts were decanted and concentrated by rotary evaporation. The residues were separately dissolved in 1 mL of methanol, and 2 µl of each sample was analysed by UHPLC-ESI-Q-TOF-MS/MS on a Bruker MaXis II mass spectrometer (or Bruker MaXis Impact mass spectrometer) coupled to a Dionex UltiMate 3000 UHPLC fitted with an Agilent Zorbax Eclipse Plus C18 column (100 × 2.1 mm, 1.8 μm). Using a flow rate of 0.2 ml/min, the column was eluted with a combination of water and acetonitrile as follows: 5% (v/v) acetonitrile for 5 min, 5−100% (v/v) acetonitrile over 21 min, 100% (v/v) acetonitrile for 3 min, 100−5% (v/v) acetonitrile over 2 min and 5% (v/v) acetonitrile for 3 min. We ran the mass spectrometer in positive ion mode (scan range: 200-3000 m/z). Source conditions were: end plate offset at −500 V; capillary at −4500 V; nebulizer gas (N2) at 1.6 bar; dry gas (N2) at 8 L min−1; dry temperature at 180 °C. Ion transfer conditions were: ion funnel RF at 200 Vpp; multiple RF at 200 Vpp; quadrupole low mass at 55 m/z; collision energy at 5.0 eV; collision RF at 600 Vpp; ion cooler RF at 50–350 Vpp; transfer time at 121 s; prepulse storage time at 1 s. Calibration was performed with 1 mM sodium formate through a loop injection of 20 µL at the start of each run. The resulting spectra were analysed with Bruker’s DataAnalysis software (v4.4).

#### A domain selectivity determination

To determine the substrate selectivity of the third A domain of TtpD (from here on referred to as TtpD-A3), we first transformed *Escherichia coli* with a plasmid containing DNA encoding the Histagged TtpD-A3-PCP didomain, and then overexpressed and purified the protein. We opted to express the didomain rather than the standalone A domain so that we could determine if both reactions that the adenylation domain catalyses took place: the adenylation reaction, which catalyses the conversion of ATP and the amino acid substrate to an amino-acyl-AMP intermediate; and the thiolation reaction, which transfers the amino acid to the phosphopantetheine arm on the PCP domain. After protein purification, we converted the didomain to its *holo* form *in vitro* by incubating it with CoA-SH and Sfp enzyme, a phosphopantetheinyl transferase which activates the didomain by transferring a phosphopantetheine arm onto the PCP domain from CoA. Then, we assessed the ability of *holo*-TtpD-A3-PCP to load various substrates by incubating the didomain with ATP and a tryptophan, phenylalanine, leucine, histidine and/or valine substrate, either separately for individual substrate assessment, or in tandem to observe which substrate was preferred. We also performed an *in vivo* competition assay by expressing the TtpD-A3-PCP didomain in *BAP1* cells^22^, a strain of *E. coli* that expresses Sfp, which allows *in vivo* domain activation and substrate loading. In all cases, substrate activation was measured with UHPLC-ESI-Q-TOF-MS analysis.

#### PCR amplification

As *Streptomyces sparsogenes* is very GC-rich, primer design with sufficiently long overlaps that still have reasonable annealing temperatures can be challenging, especially if the design requires overhangs with restriction sites for later incorporation of the DNA fragment into a plasmid. For this reason, we did two steps of PCR amplification to obtain PCR fragments containing the DNA encoding the TtpD-A3-PCP didomain: the first to amplify fragments from the GC-rich *Streptomyces sparsogenes* genomic DNA; and the second to amplify fragments with overhangs containing restriction sites (NdeI on the forward primer; EcoRI for the reverse). Detailed PCR protocols for each PCR can be found in Table S6. Primers were ordered from Merck. Resulting PCR reactions were run on an agarose gel (1g agarose in 100 ml 1x TBE buffer) for 70m at 130V, the bands were excised, and the DNA was extracted from the gel slices using the ThermoScientific Gel Extraction Kit.

#### Plasmid preparation and cloning

Next, we digested both our amplified PCR product (6 μl, ∼400 ng; 0.5 μl EcoRI; 0.5 μl NdeI; 11.7 μl dH2O; 2 μl 10X Buffer O) and the plasmid pET28A (28.3 μl, ∼2 μg; 1 μl EcoRI; 1 μl NdeI; 14.7 μl dH2O; 5 μl 10X Buffer O), which contains a T7 promoter and terminator, a kanamycin resistance cassette, an N-terminal Histag, an N-terminal thrombin cleavage site, and EcoRI and NdeI restriction sites. Enzymes and buffers were acquired from ThermoFisher scientific. Reactions were incubated for 2h at 37°C and subsequently inactivated for 20m at 65°C.

We then set up a ligation reaction to insert the digested PCR product into the digested pET28A plasmid (2 μl ligase buffer, 1 μl T4 ligase, 4 μl 5ng/μl digested pET28A, 13 μl 5ng/μl digested insert). The reaction was incubated for 5h at 16°C and subsequently stored at 4°C. Buffer and enzyme were acquired from ThermoFisher scientific. Next, we transformed our ligation reaction into chemically competent *E. coli* One Shot^TM^ TOP10 cells (ThermoFisher Scientific). We added 5 μl of our ligation reaction to one vial of competent cells and mixed gently, after which we placed the reaction on ice for 30m. Then, we heat-shocked the cells for 30s at 42°C and directly transferred them back to ice for 2m. We added 250 μl of LB medium to each vial and shook the vials horizontally at 37°C for 1h (225 rpm). Then, we spread 20 μl transformed cells onto LB plates with kanamycin (50 μg/ml).

To ensure that the plasmids had incorporated the sequence encoding the TtpD-A3-PCP didomain correctly, we inoculated 15 ml LB medium + kanamycin (50 μg/ml) with the resulting colonies, incubated them overnight at 37°C, and isolated the plasmids using ThermoFisher Scientific’s GeneJET Plasmid Miniprep Kit. We then checked the plasmids by restriction digestion and gel electrophoresis and sent plasmids that showed bands of the right sizes for sequencing. Cultures that harboured a correct plasmid were stored at -80°C.

Finally, we transformed 55.3 ng of plasmid into both chemically competent BL21* cells and chemically competent *E. coli* BAP1 cells, the latter of which express the Sfp protein responsible for activating the A-PCP didomain as described above. We incubated the reaction on ice for 15 minutes, heat-shocked the cells for 40s at 42°C and directly transferred them back to ice and added 250 μl of LB medium. We shook the vials horizontally at 37°C for 1h (225 rpm), plated 25 μl of our transformed cells onto LB plates with kanamycin (50 μg/ml), and incubated overnight at 37°C.

#### Protein overexpression and purification

To obtain sufficient protein for analysis, we inoculated 1L of LB + 50 μg/ml kanamycin with either transformed BL21* or BAP1 cells and left them to grow at 37°C to an OD of ∼1.0. Then, we added 1ml 0.5M IPTG to induce protein production. Cells were left to produce protein overnight at 15°C. Cells were spun down for 20m at 5000 rpm, 4°C. Pellets were resuspended in ∼15ml Tris washing buffer (20mM Imidazole, 20 mM Tris-HCl, 100 mM NaCl, 1-% glycerol). Next, we lysed the cells in a cell disruptor. The lysed cells were spun down for 15m at 17000 rpm, 4°C. Supernatant was transferred to a fresh tube and spun down for another 15m at 17000 rpm, 4°C. Then, the supernatant was filtered (0.45 μl filter) and loaded onto a 1ml Cytiva HisTrap-FF column. The column was washed twice with Tris washing buffer (20mM Imidazole, 20 mM Tris-HCl, 100 mM NaCl, 1-% glycerol), and the protein was eluted with elution buffers containing increasing concentrations of Imidazole (50 μM – 300 μM). Wash fractions were also collected. Fractions were denatured and run on an 8% SDS-PAGE gel for ∼30m at 180V. Fractions containing protein of the expected size were concentrated to 0.5ml (5000 MWCO, 4000 rpm, 4°C), the buffer was replaced with storage buffer (20 mM Tris-HCl, 100 mM NaCl, 1-% glycerol) four times. The resulting protein solutions had a concentration of 47.673 mg/ml for *apo*-TtpD-A3-PCP (expressed from TOP10 cells) and 32.05 mg/ml for *holo*-TtpD-A3-PCP (expressed from BAP1 cells). We analysed the proteins by UHPLC-ESI-Q-TOF-MS to assert that the proteins had the correct weights and to check which substrate was loaded by *holo*-TtpD-A3-PCP in BAP1 cells.

#### In vitro activity assay

To assess which substrate is loaded by TtpD-A3-PCP, we first converted *apo-*TtpD-A3-PCP purified from TOP10 cells to *holo-*TtpD-A3-PCP by incubating 42 μl 200 μM *apo-*TtpD-A3-PCP with 5 μl 100 mM MgCl2 (as Mg^2+^ is required by Sfp and by the A domain itself to stabilise the negative charge of ATP in the active site), 2 μl 20 mM CoA-SH (which provides the phosphopantetheine arm), and 1 μl 400 μM Sfp enzyme (isolated as described above). We incubated the reaction for 1h at room temperature, and then loaded the *holo*-enzyme with substrate by adding 50 μl 170 μM *holo*-TtpD-A3-PCP to 1 μl 100 mM ATP (required for the adenylation reaction) and 1 μl 50 μM substrate dissolved in dH2O. We tested the substrates tryptophan, phenylalanine, histidine, leucine, and valine. apo-TtpD-A3-PCP, holo-TtpD-A3-PCP, and loaded holo-TtpD-A3-PCP were analysed by UHPLC-ESI-Q-TOF-MS.

#### UHPLC-ESI-Q-TOF-MS analysis

We analysed intact apo-TtpD-A3-PCP, holo-TtpD-A3-PCP, and loaded holo-TtpD-A3-PCP on a Bruker MaXis II ESI-Q-TOF-MS connected to a Dionex 3000 RS UHPLC (equipped with an ACE C4-300 RP column (100 x 2.1 mm, 5 μm, 30°C); controlled using Bruker Otof control 4.0). The column was eluted with 0.1% formic acid and 5-100% MeCN in a linear gradient for 30m. We ran the mass spectrometer in positive ion mode (scan range: 200-3000 m/z). We used the following source settings: end plate offset: −500 V; capillary: - 4500 V; nebulizer gas (N2): 1.8 bar; dry gas (N2): 9.0 L/min; dry temperature: 200 °C. The following ion transfer conditions were used: ion funnel RF: 400 Vpp; multiple RF: 200 Vpp; quadrupole low mass: 200 m/z; collision RF: 2000 Vpp; transfer time: 110.0 µs; prepulse storage time: 10.0 µs. The resulting spectra were analysed with Bruker’s DataAnalysis software (v4.4).

## ASSOCIATED CONTENT

### Supporting Information

supporting_information.pdf (PDF) – supplementary figures, tables, and discussion.

## AUTHOR INFORMATION

### Author Contributions

The manuscript was written through contributions of all authors. All authors have given approval to the final version of the manuscript.

### Notes

G.L.C. is non-executive director, consultant and shareholder of ErebaGen Ltd. M.H.M. is a member of the scientific advisory board of Hexagon Bio.

## ABBREVIATIONS

A domain: adenylation domain
NRPS: nonribosomal peptide synthetase
BGC: biosynthetic gene cluster
PARAS: predictive algorithm for resolving adenylation domain specificity
PARASECT: predictive algorithm for resolving adenylation domain specificity by featurising enzyme and compound in tandem
Ppant: phosphopantetheine
PCP domain: peptidyl carrier protein domain
C domain: condensation domain
PCA: principal component analysis

