## supporting_information.pdf for "PARAS: high-accuracy machine learning of substrate specificities in nonribosomal peptide synthetases"

### Supplementary Discussion

#### Qui18-A1 and BreC-A3 recognise tryptophan in different orientations

We compared the 3D architectures of the active sites of Qui18-A1 and BreC-A3 to determine how two domains with highly divergent active site sequences might recognise the same substrate. To this purpose, we built AlphaFold models[19] for both domains and subsequently performed molecular docking using tryptophanyl-adenylate (TYM) as the ligand. While the docking of ligands onto static structures only allows estimating ligand-protein interactions, our results suggest that it is improbable that the two domains recognise Trp in the same orientation (Figure S20). The tryptophan moiety of the TYM ligand docked to Qui18-A1 (Figure S20a) is rotated ~90° compared to BreC-A3-docked TYM (Figure S20b), shifting the side chain slightly towards the adenyl moiety of TYM such that the aromatic nitrogen is positioned 8.9Å away from its corresponding position in BreC-A3 (Figure S20d). Also, clear differences can be seen in the lobes of the active site pocket. In contrast, the channels that bind the adenyl moiety of the substrate are far better conserved, as also reflected by the similar way in which this portion of the TYM substrate is docked in both domains.

#### Structure-based extraction of the A domain active site improves the performance of predictive models

As structural modelling is time-intensive, we wanted to examine structure-informed methods that do not require the modelling of domains that the user wants to query. For this reason, we first focussed our attention to structure-based extraction of the 34 amino acid active site signatures as also employed by NRPSPredictor2. Current active site extraction methods rely on profile alignments, which use a sequence alignment as guide and append a query sequence to the alignment such that residues of interest can be identified and extracted. It can be challenging to obtain a sequence alignment for A domains that adequately aligns the active site: this region of the domain is highly variable, especially the residues of interest, and it can therefore be difficult to determine from sequence alone which residues truly belong to the active site. In particular, we observed that active site extractions made using sequence-guided alignments contained relatively many gaps, while typically one would not expect gaps to occur in an active site. We reasoned that a structure alignment might resolve this, as it uses the three-dimensional position of residues to align sequences rather than residue identity to construct alignments. Indeed, extended active site signatures extracted using a structure-based alignment contained ten times fewer gaps, with an average of 0.06 gaps per active site signature for structure-guided extraction compared to 0.58 gaps per active site signature for sequence-guided extraction. For 10-aa active site signatures, this difference was much smaller: 0.03 gaps per active site for structure-guided extraction compared to 0.04 gaps for sequence-guided extraction. We also investigated mismatches, and found that on average, 1.01 residues were different for each active site signature (1.65 including gaps), and 0.12 for each Stachelhaus code (0.20 including gaps). Most of these mismatches were accounted for by alignments to tryptophan (Figure S3a,c), which were often aligned against gaps or different residues. This makes sense, considering that the BLOSUM62 matrix, which is typically used for local sequence alignments, highly penalises mismatches with tryptophan. We also examined how often each residue of the Stachelhaus code and active site signature contained a mismatch. We observed that residues 278 and 331 (indexed according to the reference structure 1AMU) accounted for most mismatches in the Stachelhaus code, and residues 213 and 214 contributed most towards mismatches in the active site signatures (Figure S3b,d).

To see if these differences led to differences in predictive power, we built homology models for a random subset of our dataset – a subset as profile alignment scales poorly with the number of sequences – and built ten different structure alignments for cross-validation purposes, each using 90% of the data. Then we trained basic predictive models to assess if there was a difference in performance when the active sites were extracted using sequence alignments or structure alignments as guide. On average, models trained on active sites that were extracted using structure-based guide alignments outperformed those that were trained on residues extracted from sequence-based guide alignments by about 1.7% (paired t-test: $p=0.03$). This demonstrates that it is still possible to leverage structural information to improve model performance, even when structural data of the queried domain is not directly used.

While active site extraction using sequence- or structure-guided profile alignments is fast compared to structural model computation or phylogenetic placement methods, it still takes ~2 seconds per A domain. For this reason, we also explored pHMM extraction methods, comparing both the HMMER2 pHMMs that were used by Rausch *et al.* for the first version of NRPSPredictor, and the AMP-binding pHMMs from the PFAM database (PF00501, PF13193).

#### Identification of tryptopeptin congeners in *S. coelicolor* M1154, *S.coelicolor* M1154 containing pCAP1000maedttpBGC, and pCAP1000spattpBGC

One metabolite yielded $[M+H]^+$ ions with m/z =559.3483 (calculated for $C_{30}H_{47}N_4O_6^+$: 559.3490) was proposed as tryptopeptin congener **1** with variation on the N-acyl group and decrease of one degree of unsaturation on the epoxyketone moiety compared with tryptopeptin A. Another metabolite yielded $[M+H]^+$ ions with m/z =517.3390 (calculated for $C_{28}H_{45}N_4O_5^+$: 517.3384) was proposed as tryptopeptin congener **2** with decrease of one degree of unsaturation on the ethyl ketone group compared with tryptopeptin B. The other two metabolites yielded with $[M+H]^+$ ions with m/z= 531.3543 (calculated for $C_{29}H_{47}N_4O_5^+$: 531.3541) and m/z= 545.3698 (calculated for $C_{30}H_{49}N_4O_5^+$: 545.3697) were proposed to be tryptopeptin congener **3**, **4** that contain further variation on N-acyl compared with tryptopeptin congener **2.** The metabolite yield with $[M+H]^+$ ions with m/z= 543.3536 (calculated for $C_{30}H_{47}N_4O_5^+$: 543.3541) was proposed as tryptopeptin congener **5** with alternation to the N-acyl group compared with tryptopeptin B. The proposed planar structures were consistent with the MS/MS analysis (Figure S11).

#### Supplementary Figures.

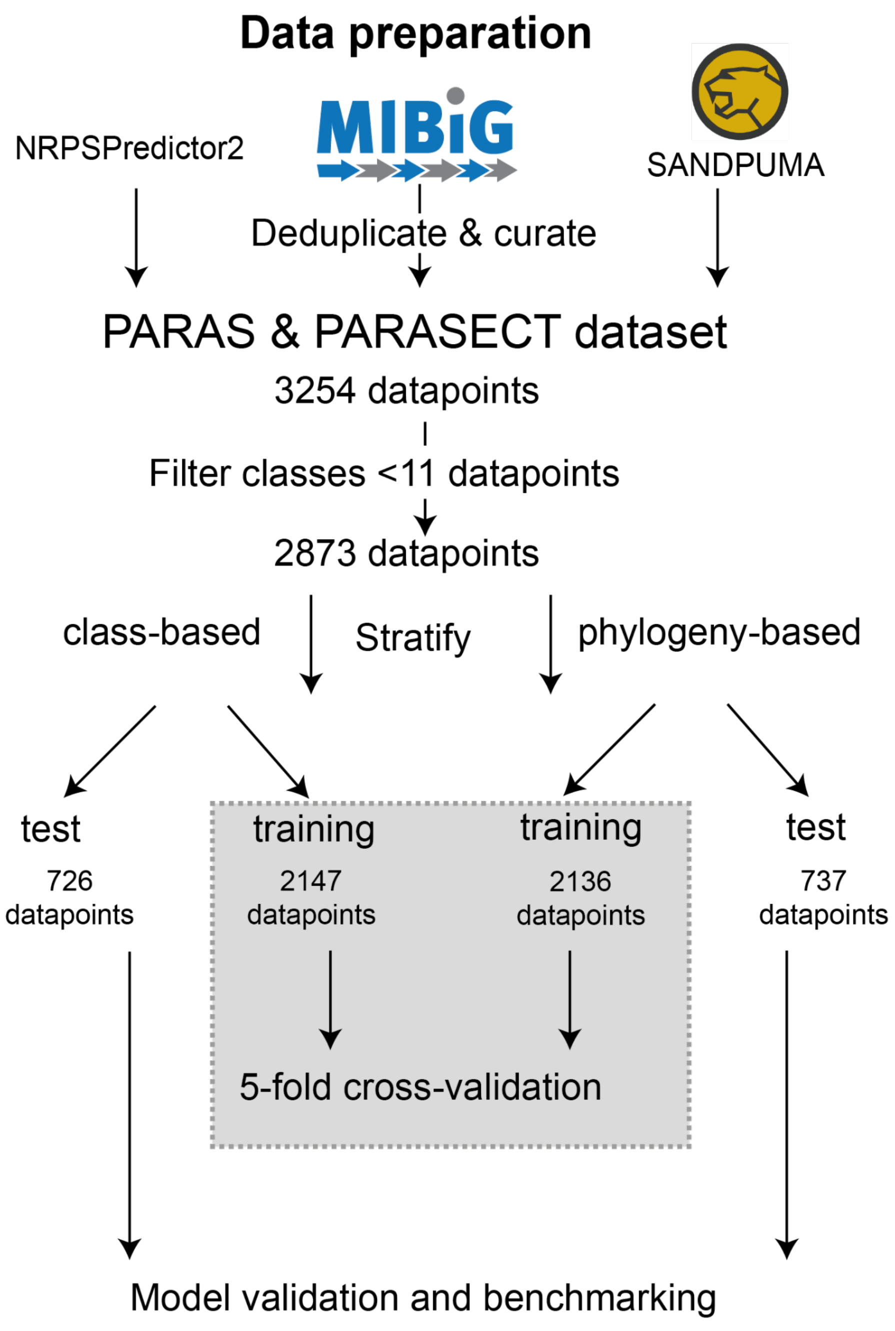

**Figure S1.** Overview of data selection, deduplication, filtering, and division across train-test sets.

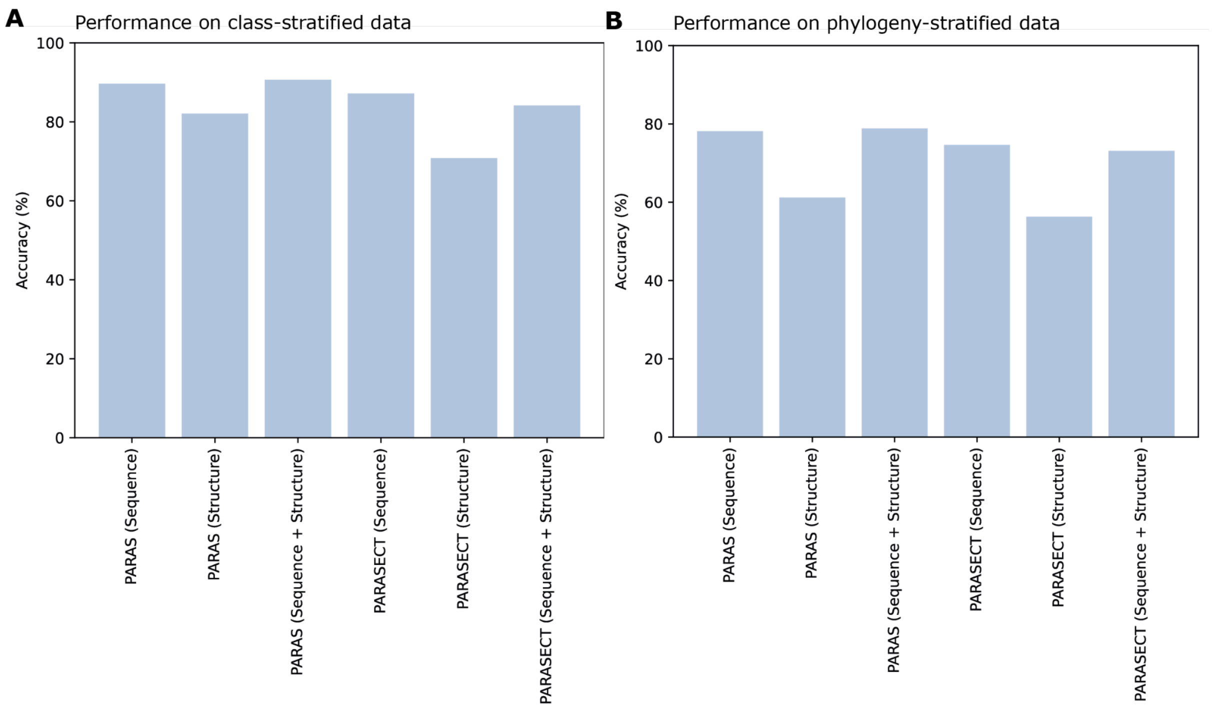

**Figure S2. Influence of featurisation method on model performance.** a) Model performance on a hold-out test set stratified on substrate class. b) Model performance on a hold-out test set stratified on A domain taxonomy. Models that were trained on sequence features perform substantially better than those trained only on structural features, and adding structural features to the model does not significantly improve model performance.

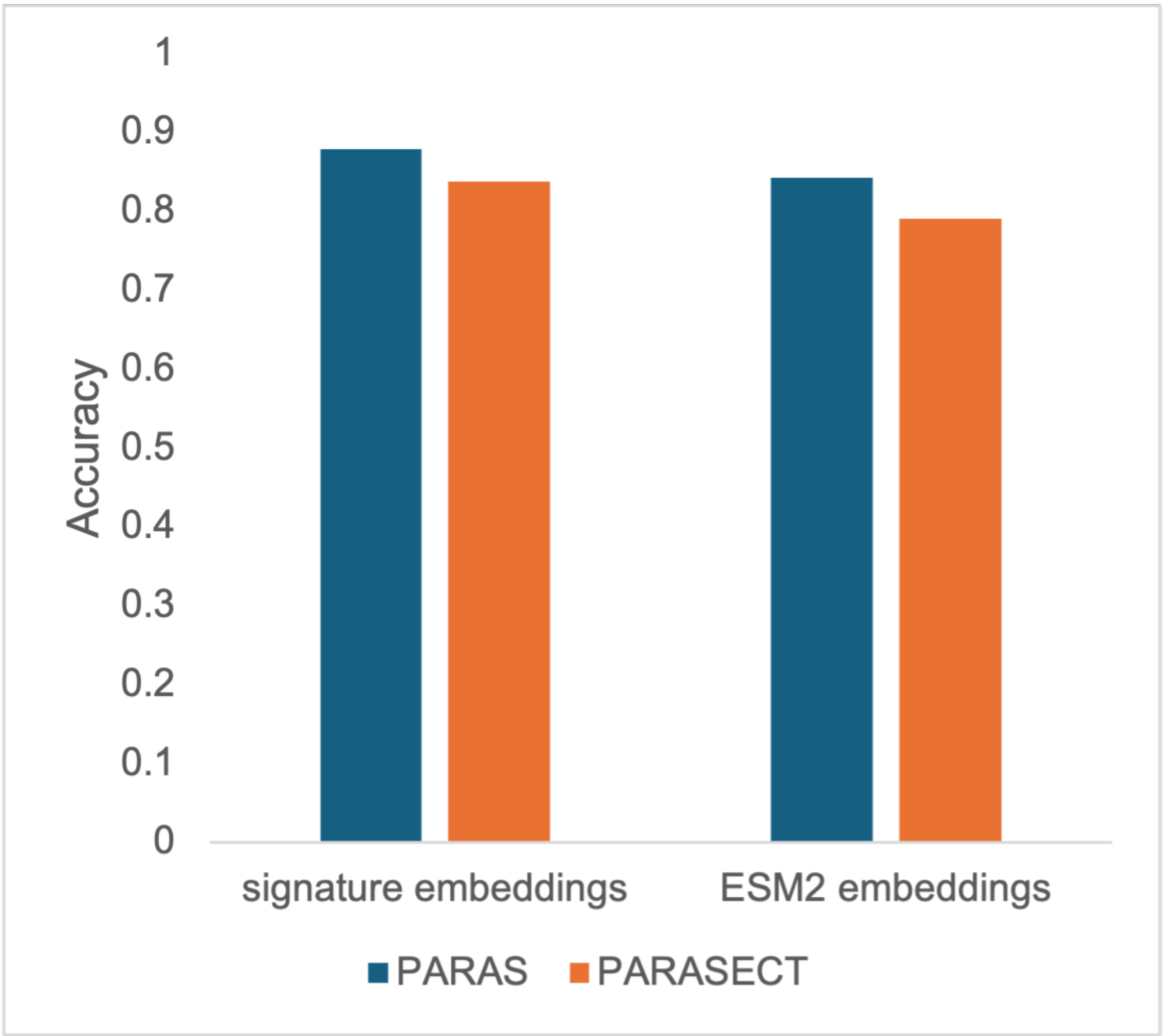

**Figure S3. Signature embeddings outperform ESM2 embeddings.** Signature embeddings comprise 15 physicochemical properties per residue of the 34 amino acid active site signature. ESM2 embeddings comprise the first 100 principal components of a 34 x 1280 = 43530 residue-specific ESM embedding space, embedding the 34 amino acids of the active site.

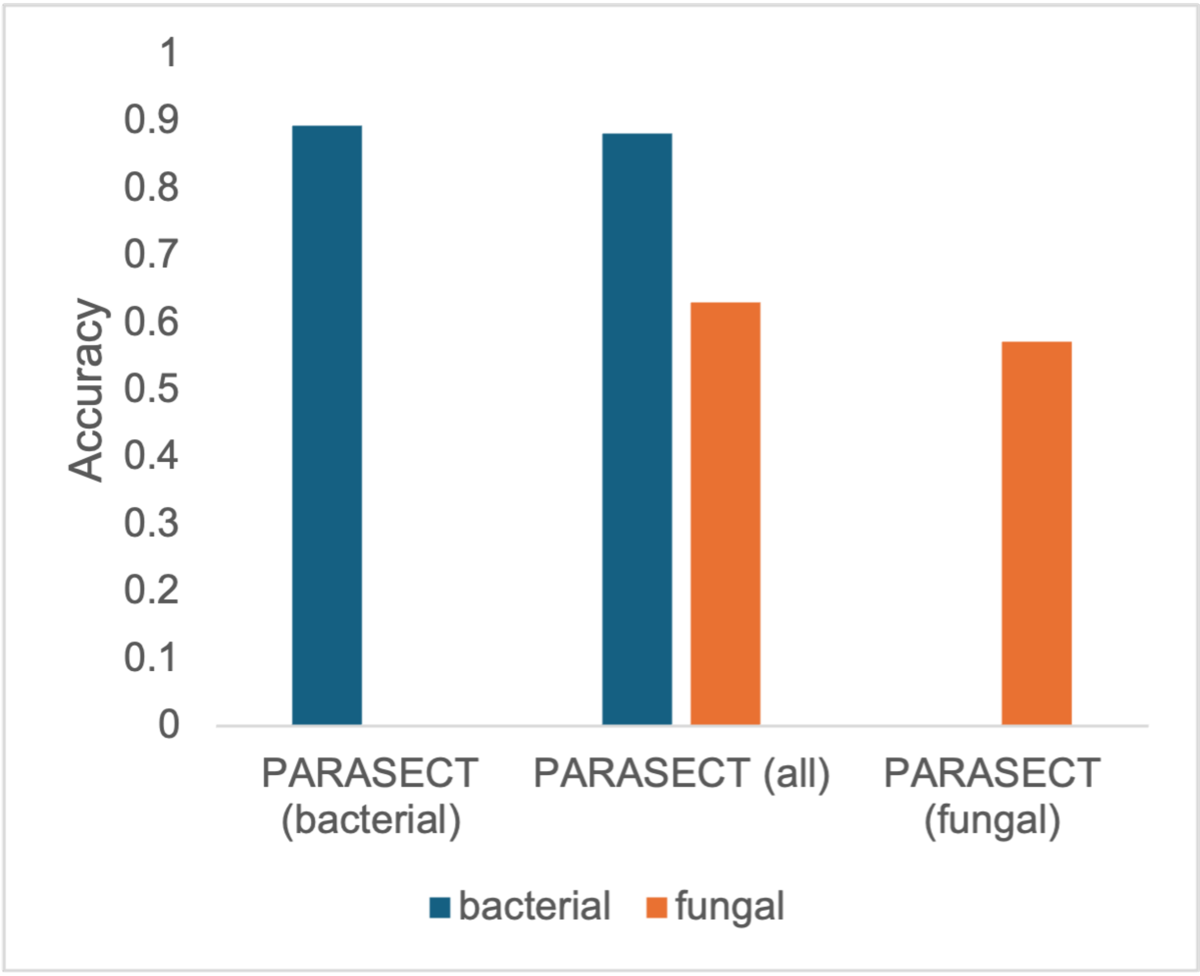

**Figure S4. Performance of PARASECT models trained on bacterial and fungal data subsets.** Inclusion of bacterial data boosts PARASECT accuracy on fungal sequences, but inclusion of fungal data hurts PARASECT accuracy on bacterial sequences.

.

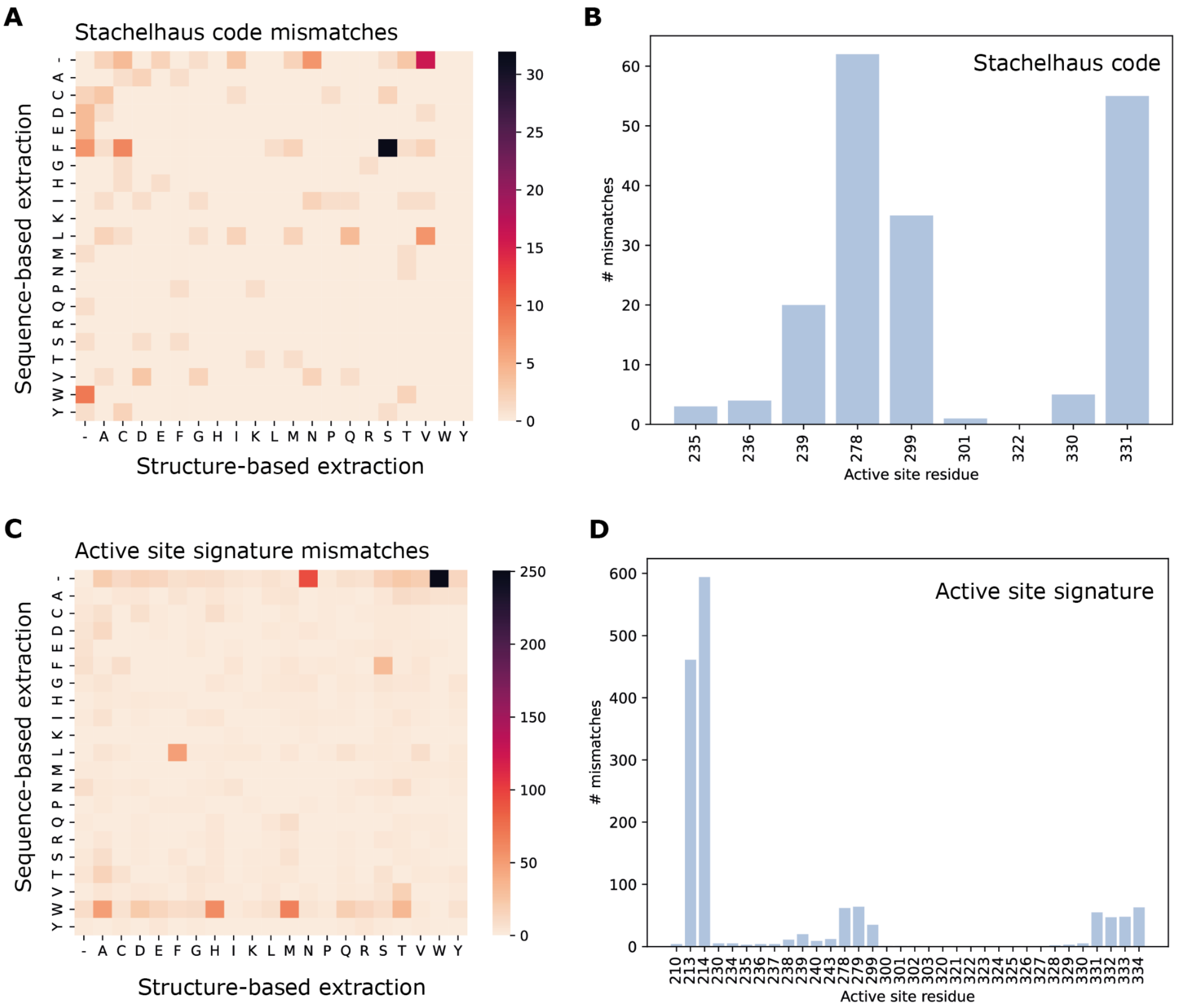

**Figure S5. Differences between active sites extracted using sequence-based and structure-based approaches.** a) Active site mismatches between sequence-based and structure-based extraction per residue type. b) Active site mismatches per active site position. c) Extended active site signature mismatches between sequence-based and structure-based extraction per residue type. d) Extended active site signature mismatches per active site position. Active site positions are labelled according to positions in the reference structure 1AMU.

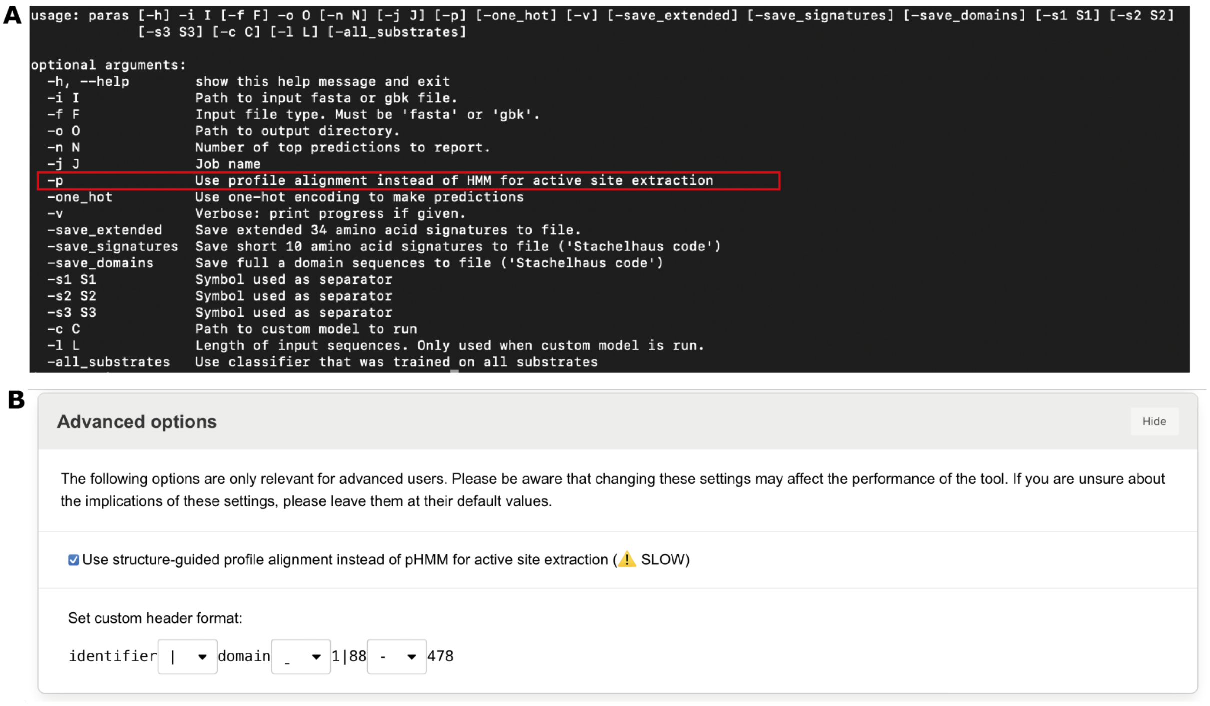

**Figure S6. User-toggleable options for selecting the active site extraction method.** pHMM-based active site extraction is selected by default. a) PARAS command-line interface. The red box indicates which option to toggle for structure-guided profile alignment-based active site extraction. b) Structure-guided profile alignment-based active site extraction can be selected in the advanced options on the PARAS website.

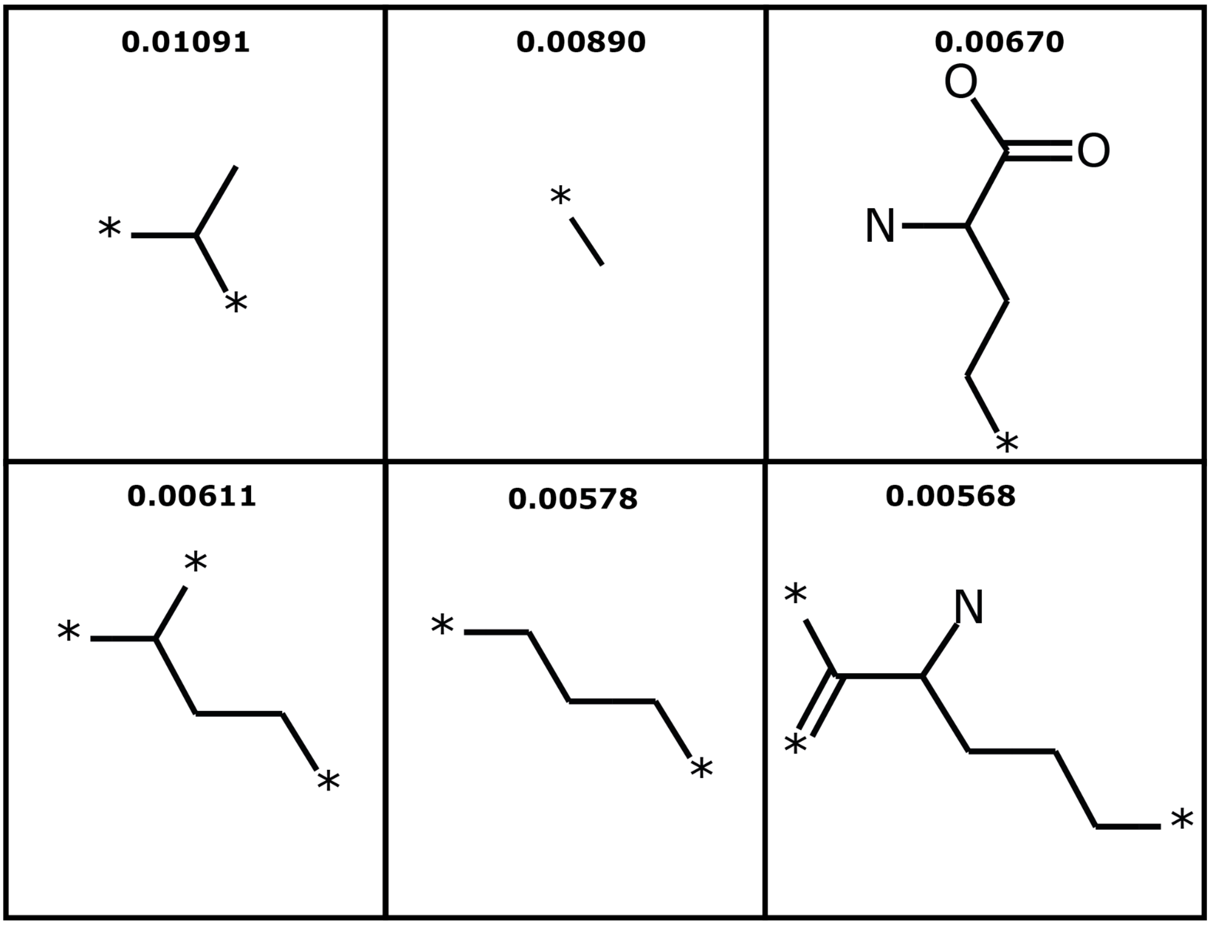

**Figure S7. Six most informative features used in sequence-based PARASECT model.** All six most informative features are substrate features. Stars indicate rest groups. Numbers at the top of panels show feature importances (with a feature importance of 1, the random forest would exclusively use that feature to make classifications). Fingerprints representing methyl groups and long unbranched chains are the most important features in PARASECT. Feature importances were obtained by averaging across 5 sequence-based cross-validation models.

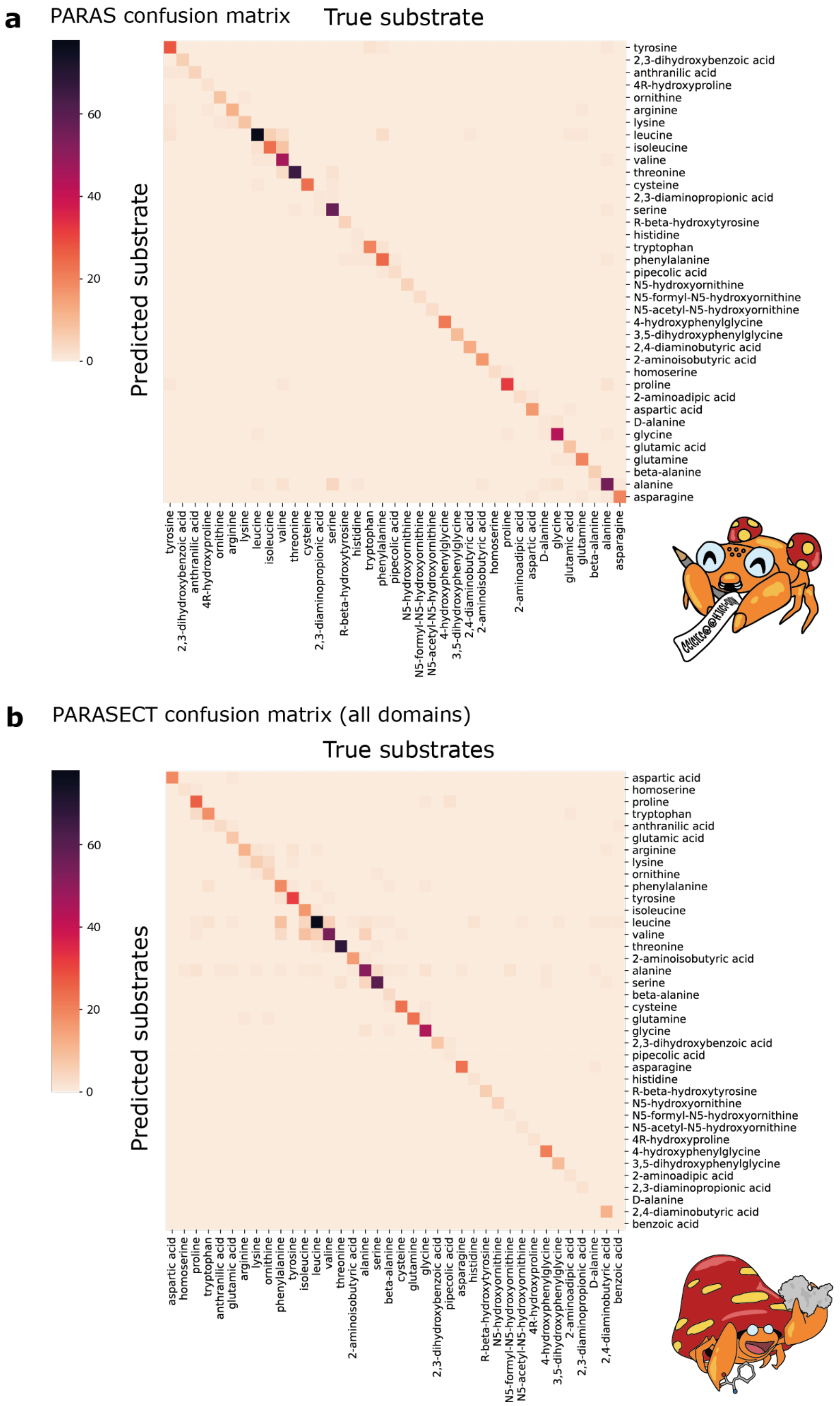

**Figure S8. Confusion matrices showing PARAS and PARASECT predictions per substrate.** Darker colours indicate a higher number of predictions. a) Confusion matrix for the multi-class predictor PARAS**.** All predictions are displayed in the matrix. b) Confusion matrix for the binary predictor PARASECT. For each domain, only predictions predicting an interaction probability >0.5 are displayed in the matrix.

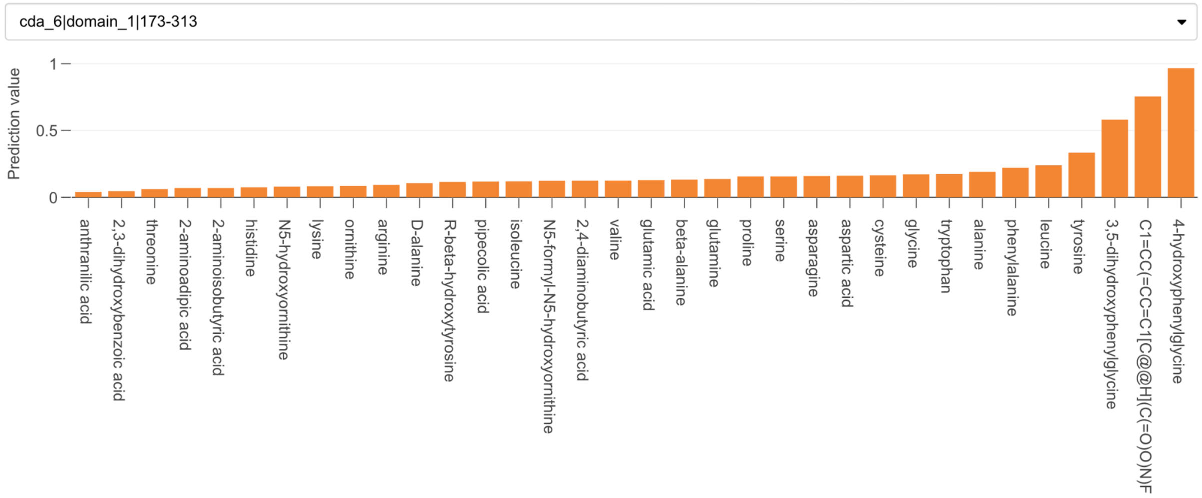

**Figure S9. Results of a PARASECT run on the 6th A domain of the CDA BGC.** The A domain was queried with all 34 standard substrates as well as 4-fluorophenylglycine, which was provided as a SMILES string.

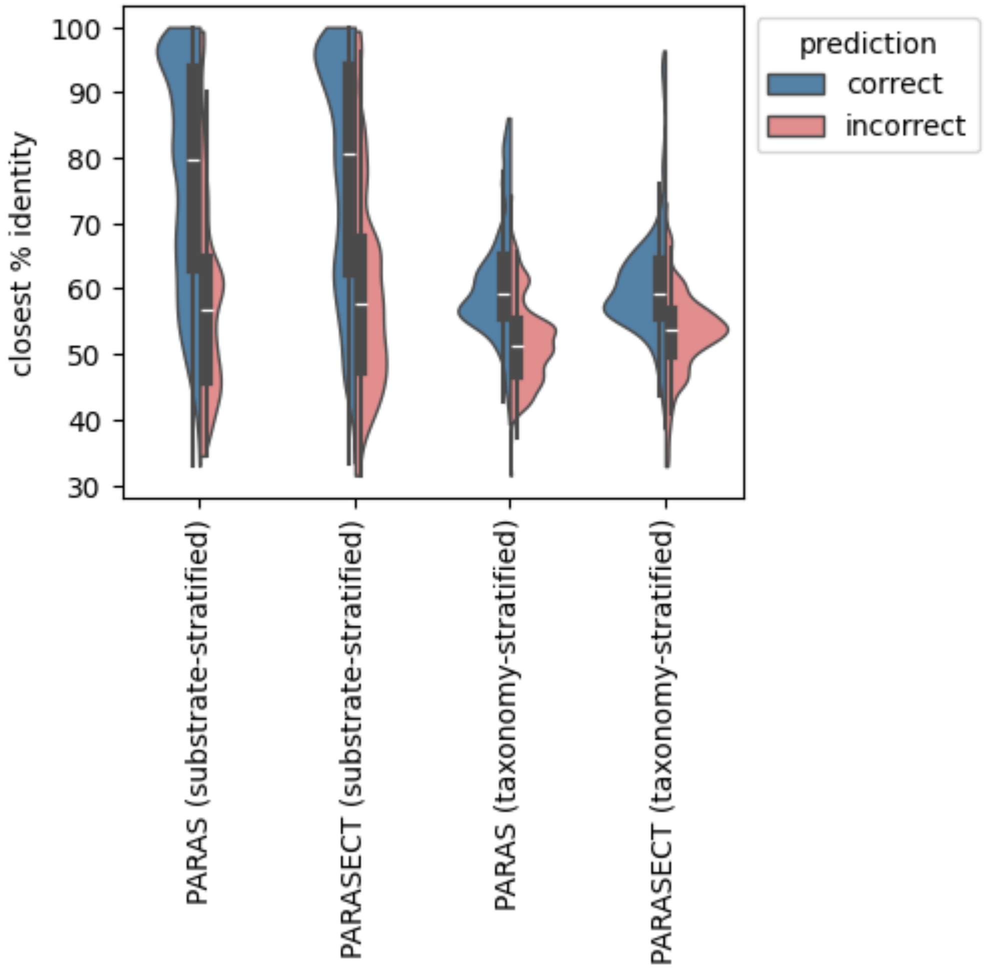

**Figure S10. Violin plots of PARAS/PARASECT test data.** Prediction accuracy increases with % identity to the most similar sequence in the training data.

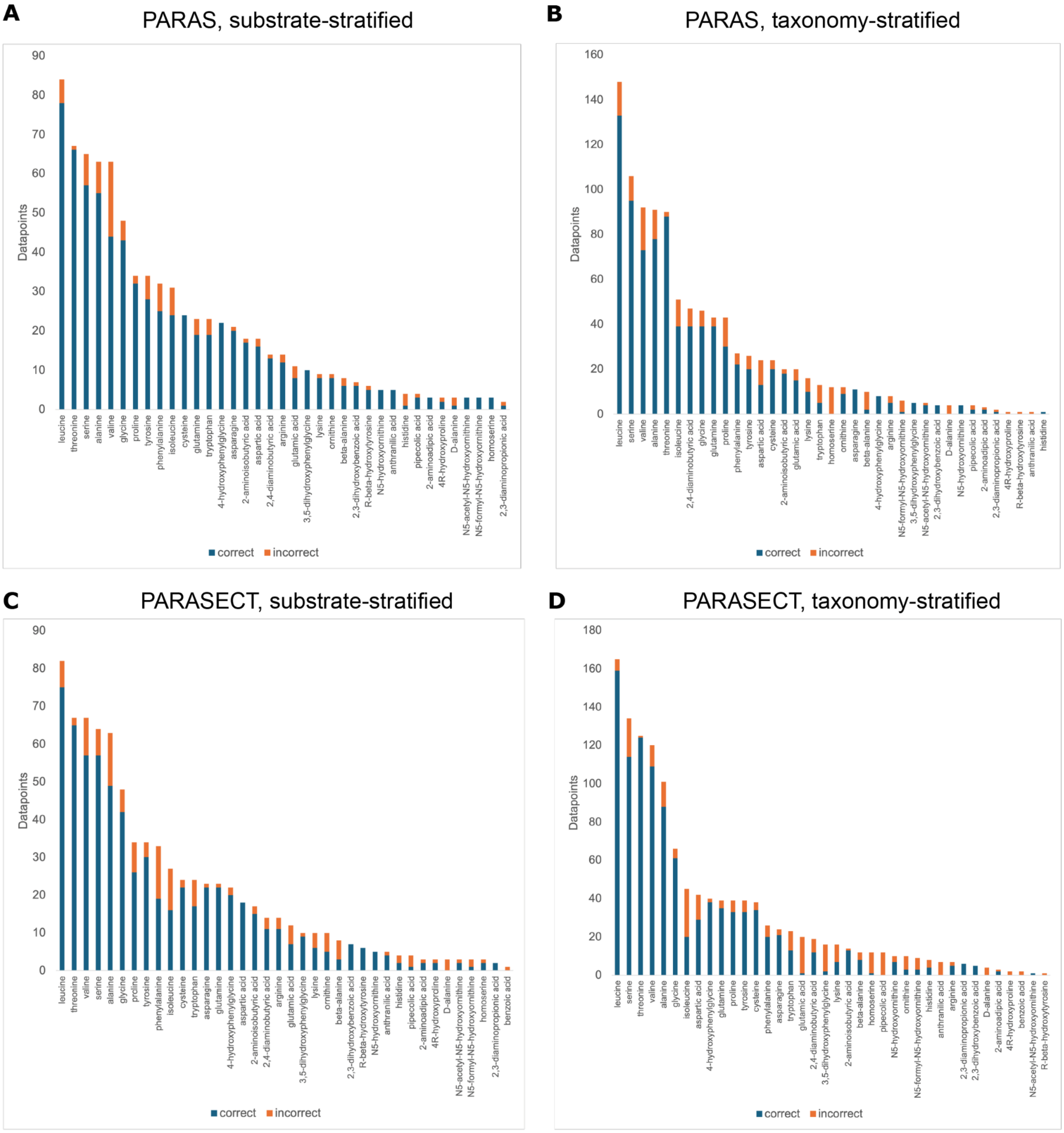

**Figure S11. Substrate specific performance of PARAS and PARASECT models.** a) Sequence-based PARAS performance on a hold-out test set stratified on substrate. b) Sequence-based PARAS performance on a hold-out test set stratified on A domain taxonomy. c) Sequence-based PARASECT performance on a hold-out test set stratified on substrate. d) Sequence-based PARASECT performance on a hold-out test set stratified on A domain taxonomy. Large and uncommon substrates perform substantially worse using taxonomy-based stratification methods, indicating the need for a phylogenetically diverse training set. For exact accuracy values, see Table S7.

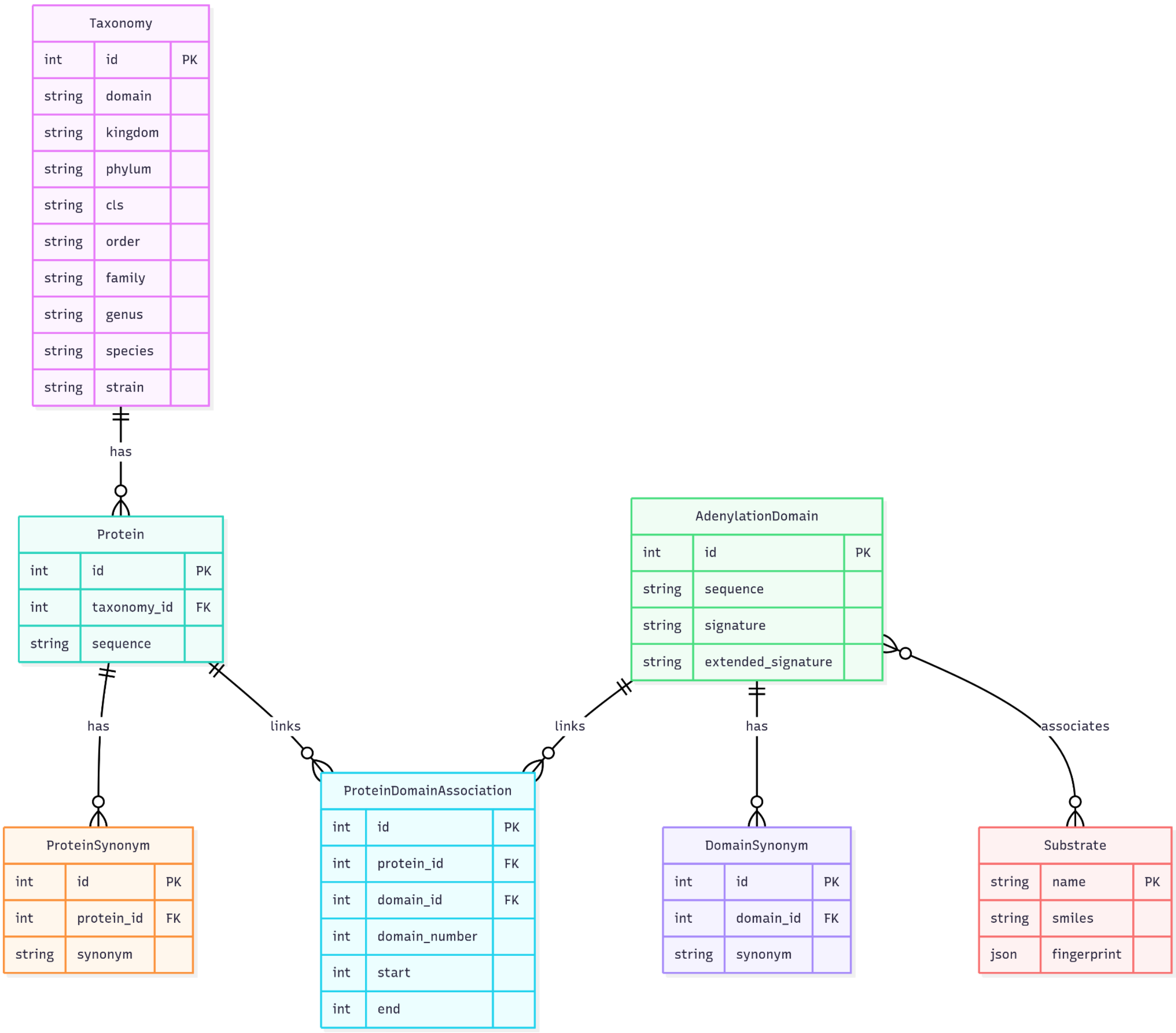

**Figure S12. PARASECT SQL database architecture.** The PARASECT database stores the following data. Adenylation domains store information on domain sequence, 10 amino acid active site signature, and 34 amino acid active site signature. This table is linked to a domain synonym table, which stores different synonyms that the domain is known as (domain synonyms are formatted as protein_identifier.A[domain index in protein], e.g. ACI30655.1.A2); to a substrate table, which stores the name, SMILES string, and molecular fingerprint of the substrate; and to a protein-domain association table, which stores the start and end coordinates of the domain within the protein, and the domain index within the protein. The protein-domain association table in turn links to a protein table, where full protein sequences are stored. Protein entries also link to the taxonomy table, which stores the taxonomic lineage of a protein; and to a protein synonym table, which stores different synonyms for the protein (e.g. uniprot ID, protein name, GenPept ID).

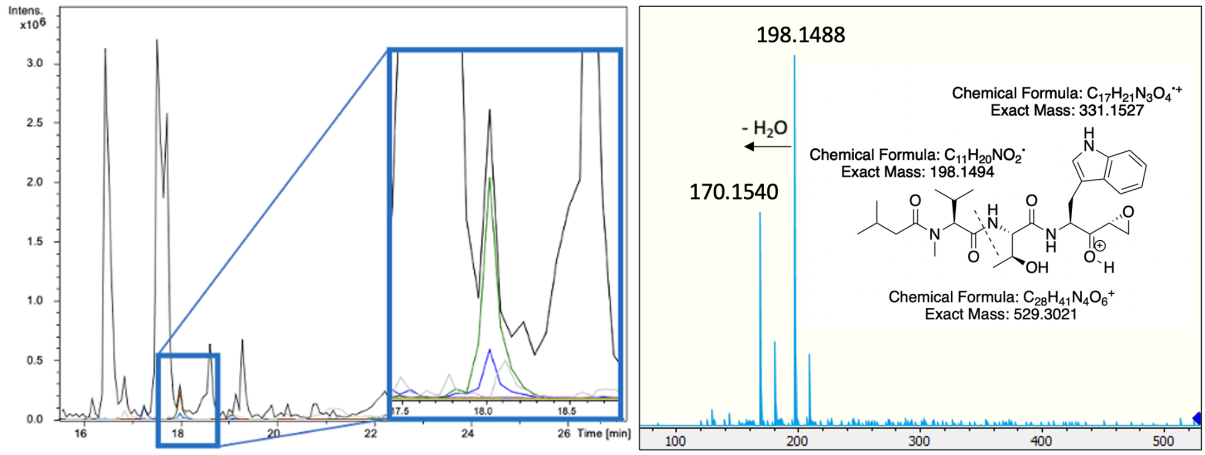

**Figure S13. LCMS analysis of metabolite extracts from S. sp. maeda85 after 5 days of growth on plates.** a) Base peak chromatogram (black), and extracted chromatograms for m/z 529.3021 (blue) and m/z 551.2840 (green) corresponding to [M+H]+ and [M+Na]+ adduct of tryptopeptin A respectively. b) MS2 fragments of m/z 529.3021 corresponding to [M+H]+ with corresponding structure.

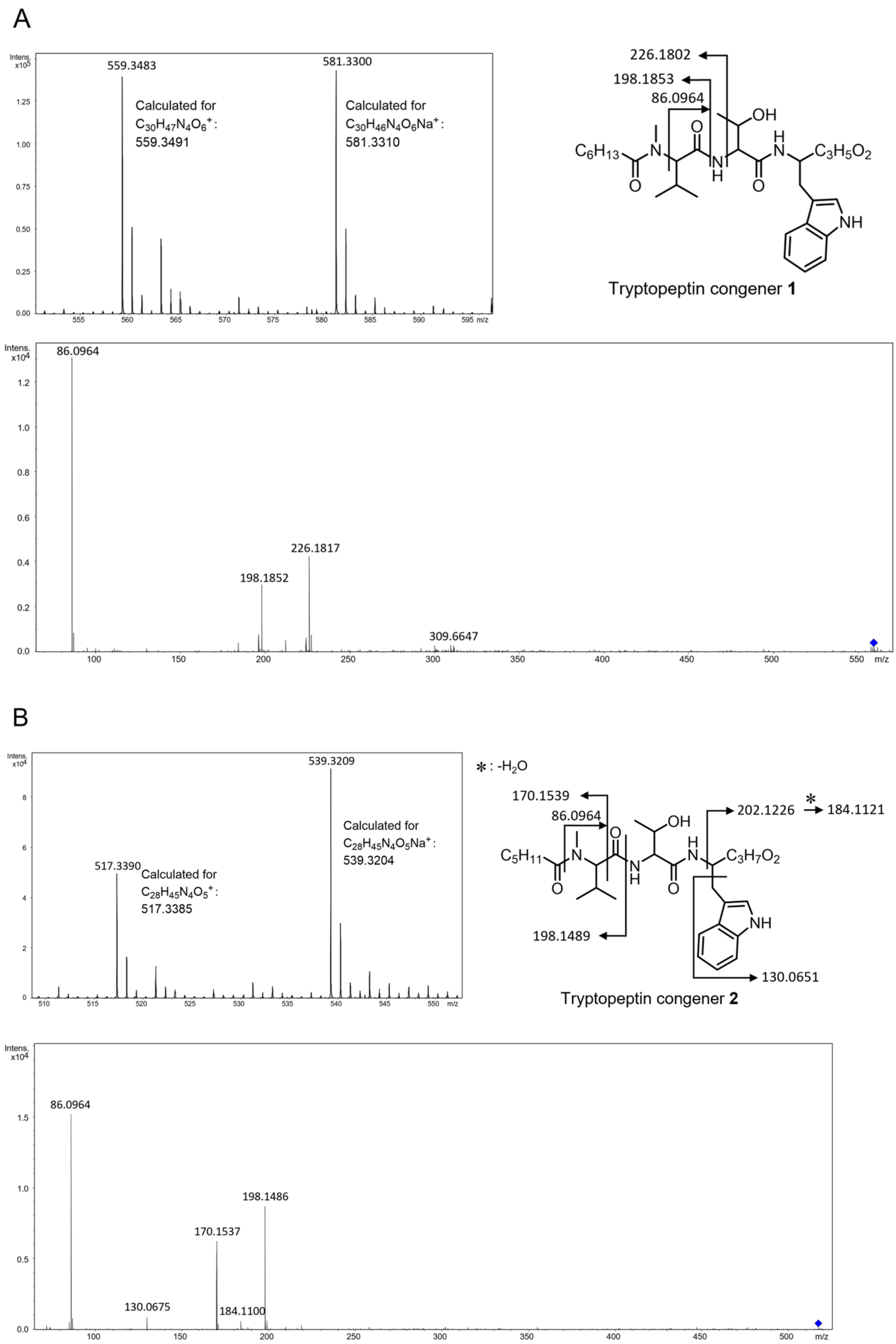

A
Intens. x10^5
559.3483
581.3300
Calculated for $C_{30}H_{47}N_4O_6^+$: 559.3491
Calculated for $C_{30}H_{46}N_4O_6Na^+$: 581.3310
226.1802
198.1853
86.0964
OH
$C_6H_{13}$
$C_3H_5O_2$
NH
Tryptopeptin congener **1**
Intens. x10^4
86.0964
226.1817
198.1852
309.6647
m/z
B
Intens. x10^4
539.3209
517.3390
Calculated for $C_{28}H_{45}N_4O_5^+$: 517.3385
Calculated for $C_{28}H_{45}N_4O_5Na^+$: 539.3204
*: $-H_2O$
170.1539
86.0964
202.1226
184.1121
OH
$C_5H_{11}$
$C_3H_7O_2$
198.1489
NH
130.0651
Tryptopeptin congener **2**
Intens. x10^4
86.0964
198.1486
170.1537
130.0675
184.1100
m/z

C

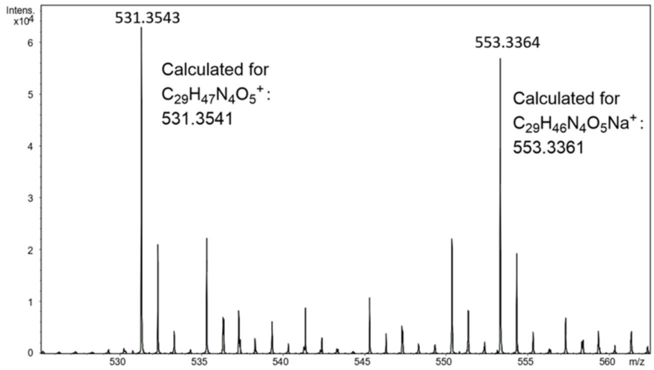

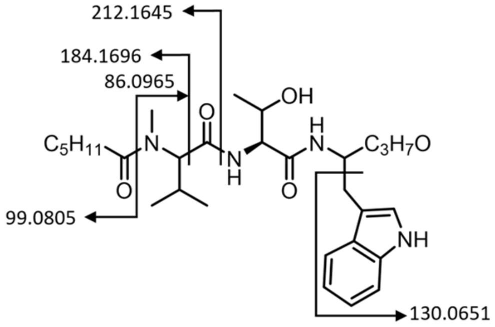

Tryptopeptin congener **3**

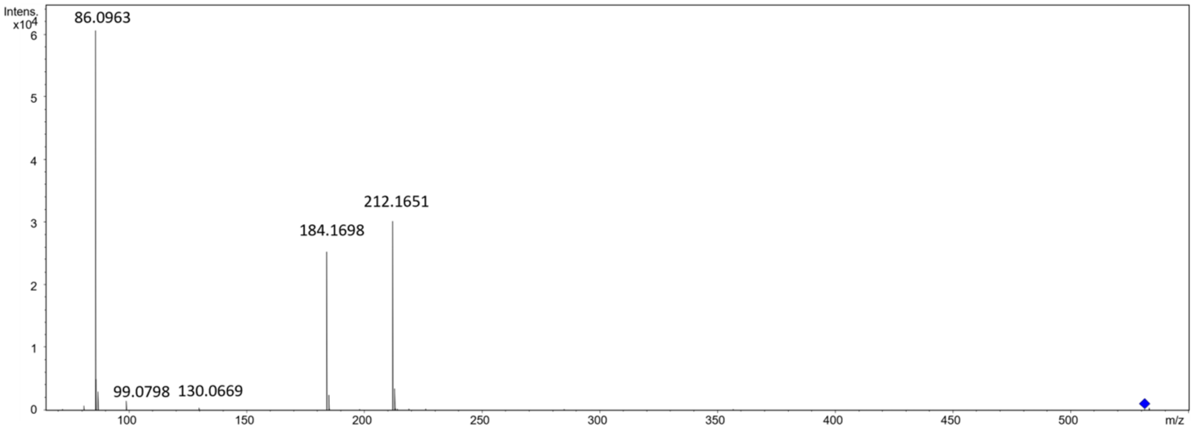

D

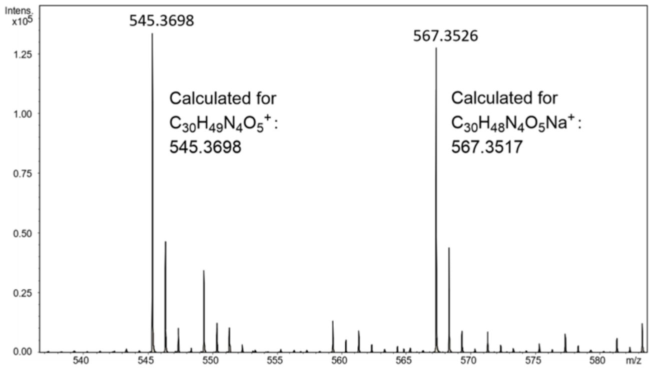

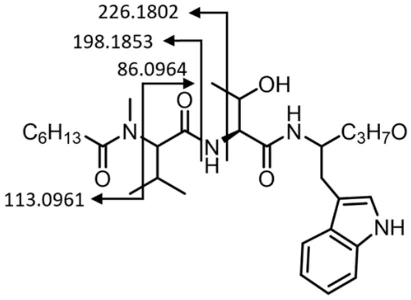

Tryptopeptin congener **4**

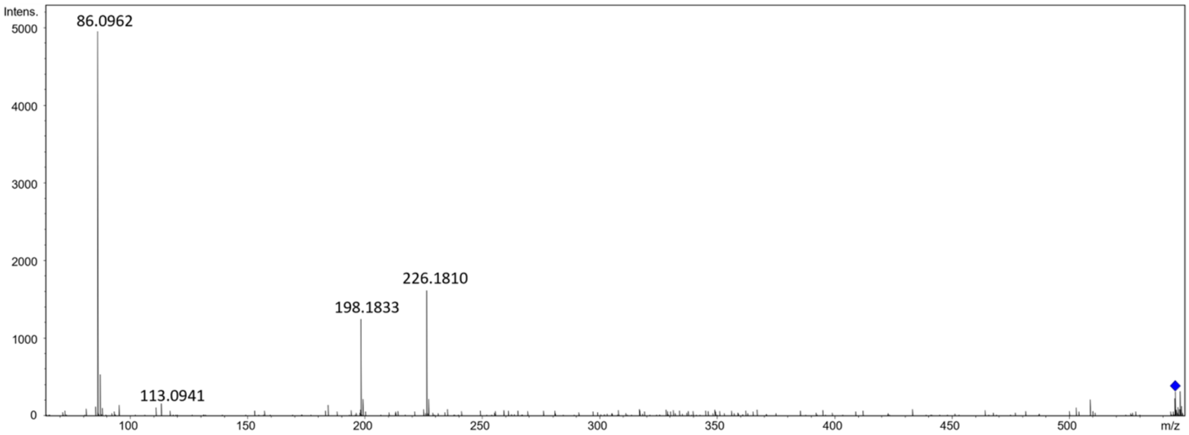

E

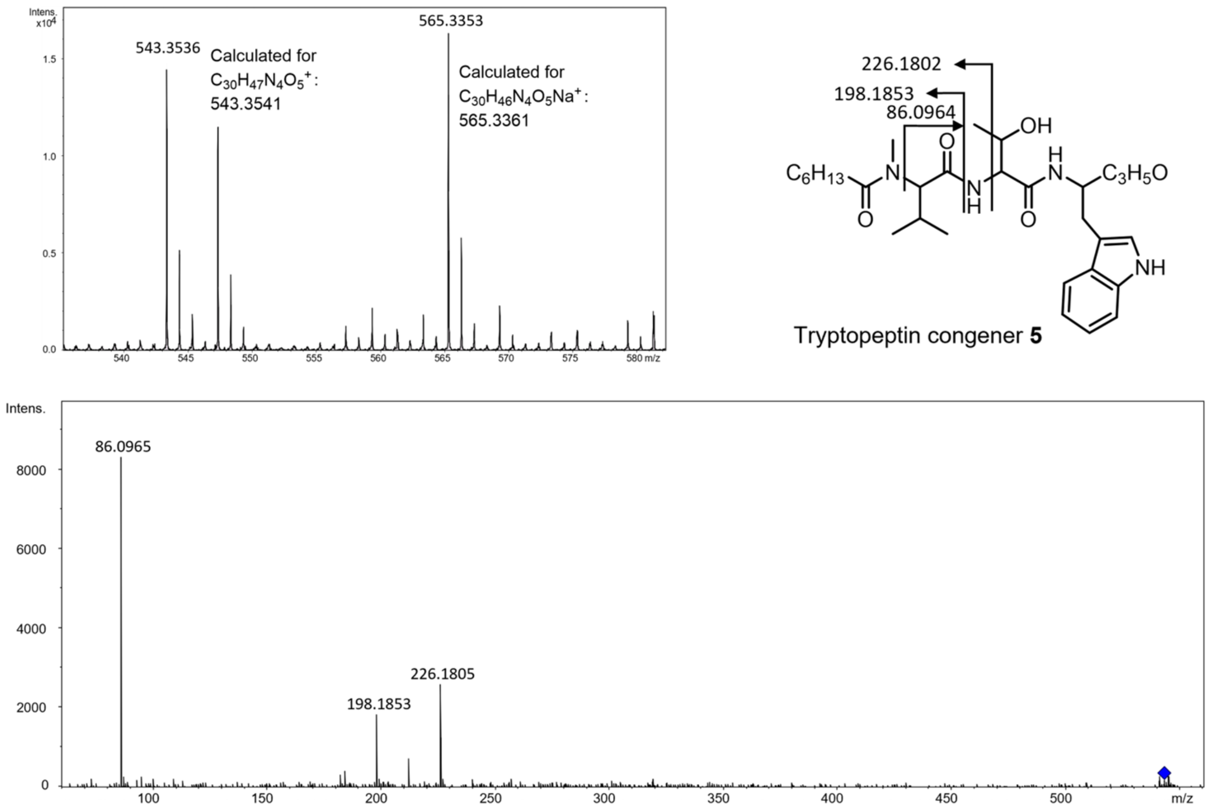

**Figure S14. High resolution and tandem mass spectra from UHPLC-ESI-Q-TOF-MS/MS analyses of culture extracts from *S. coelicolor* M1154 containing pCAP1000spattpBGC.** a) Tryptopeptin congener 1. b) Tryptopeptin congener 2. c) Tryptopeptin congener 3. d) Tryptopeptin congener 4. e) Tryptopeptin congener 5.

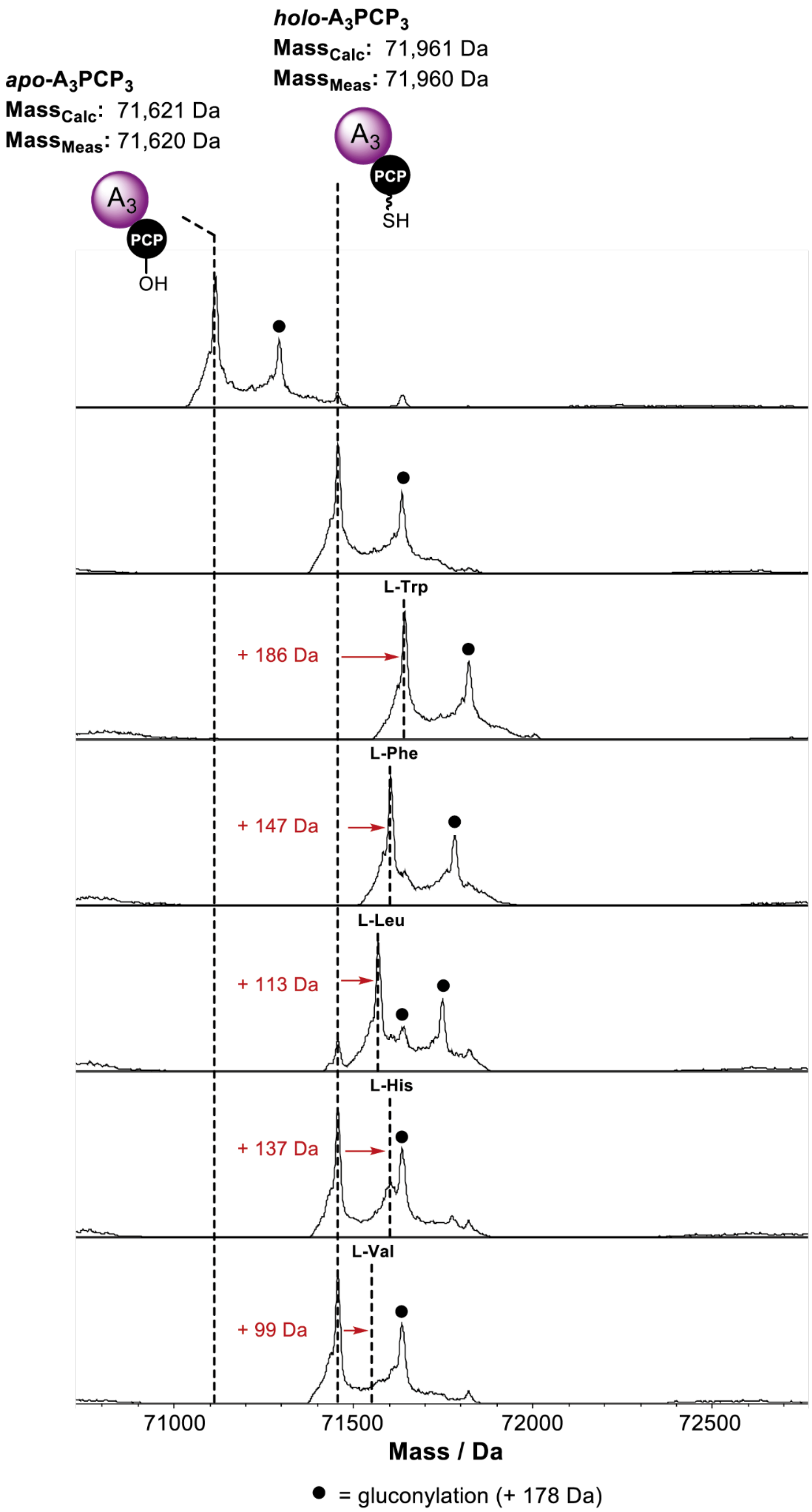

**Figure S15. Spectra of apo-TtpD-A$_3$PCP$_3$ and in vitro activated holo-TtpD-A$_3$PCP$_3$ incubated with various amino acids.** Expected mass shifts upon incubation of holo-TtpD-A$_3$PCP$_3$ with different amino acid substrates are indicated with red arrows and text. Peaks at these mass shifts indicate complete activation for tryptophan and phenylalanine; partial activation for leucine; minimal activation for histidine; and no activation for valine.

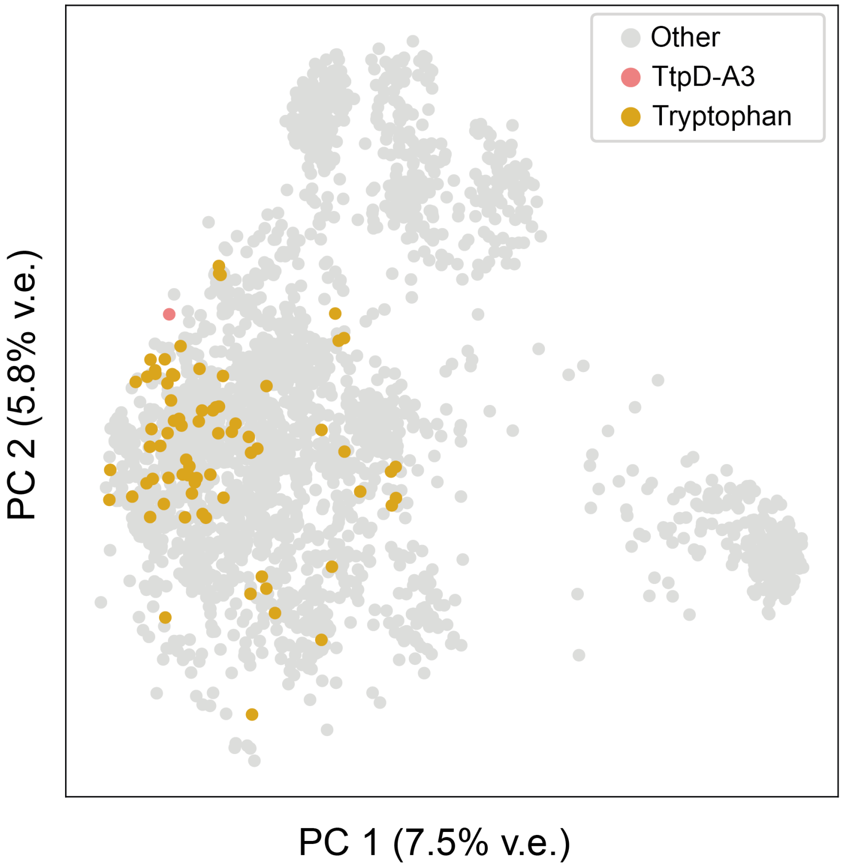

**Figure S16. PCA of 3D A domain active sites based on 3254 AlphaFold models.** The TtpD-A3 domain and A domains recognising L-Trp are highlighted.

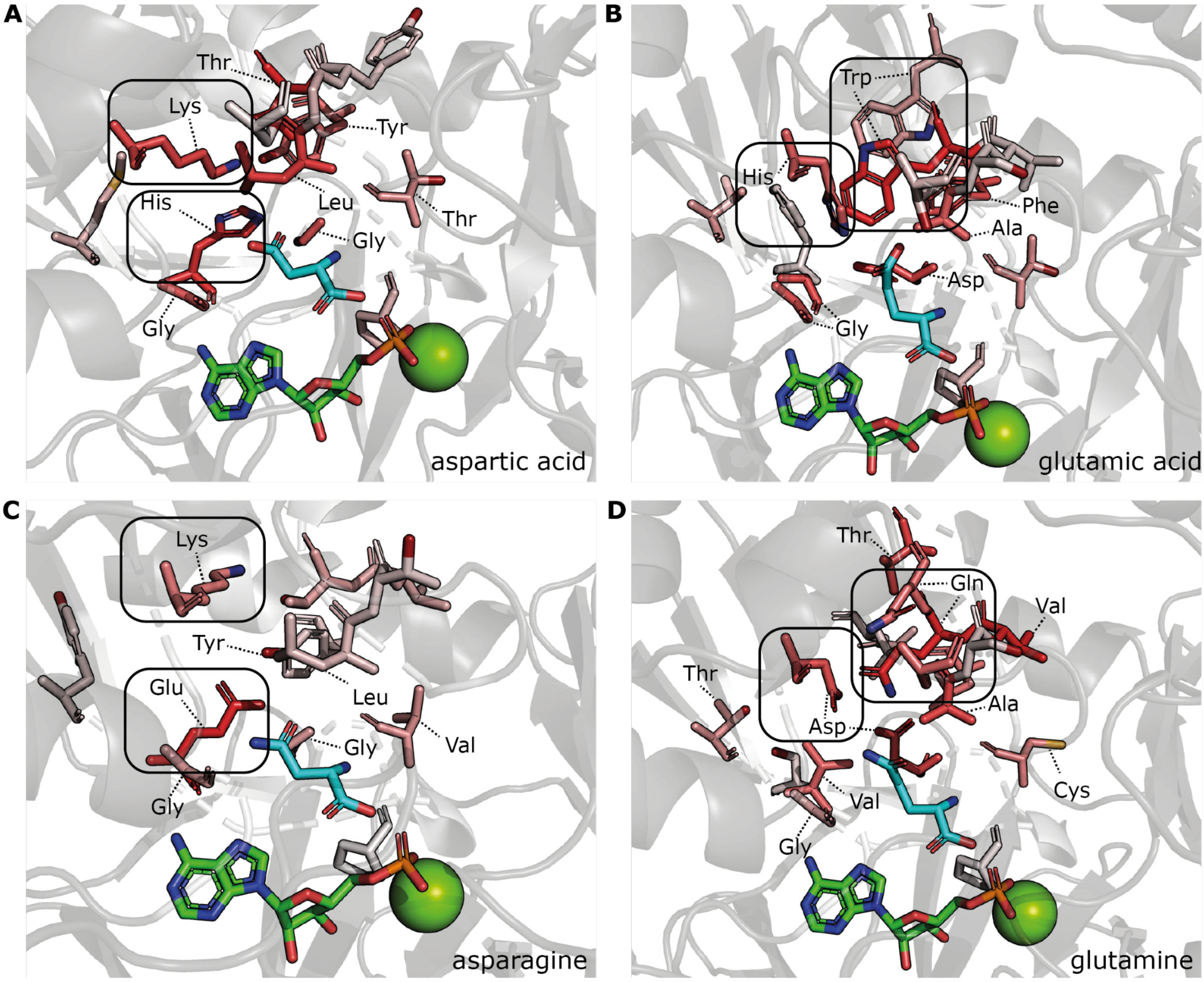

**Figure S17. Most predictive residues for Asx/Glx substrates.** a) Aspartic acid. b) Glutamic acid. c) Asparagine. d) Glutamine. A domains selecting Asx and Glx substrates use different residues to coordinate the charges of the side chain: residues 13 (top box) and 22 (bottom box) for Asx, and residues 10 (right box) and 13 (left box) for Glx. This is to accommodate the difference in side chain length between these substrates: residue 10 lies deeper in the pocket than residue 22, and therefore can interact with glutamine and glutamic acid side chains, while residue 22 is optimally positioned to interact with aspartic acid and asparagine. A domains recognising substrates with acidic side chains have more negatively charged pockets than those recognising asparagine and glutamine, containing a highly informative histidine residue at position 22 for aspartic acid and at position 13 for glutamic acid.

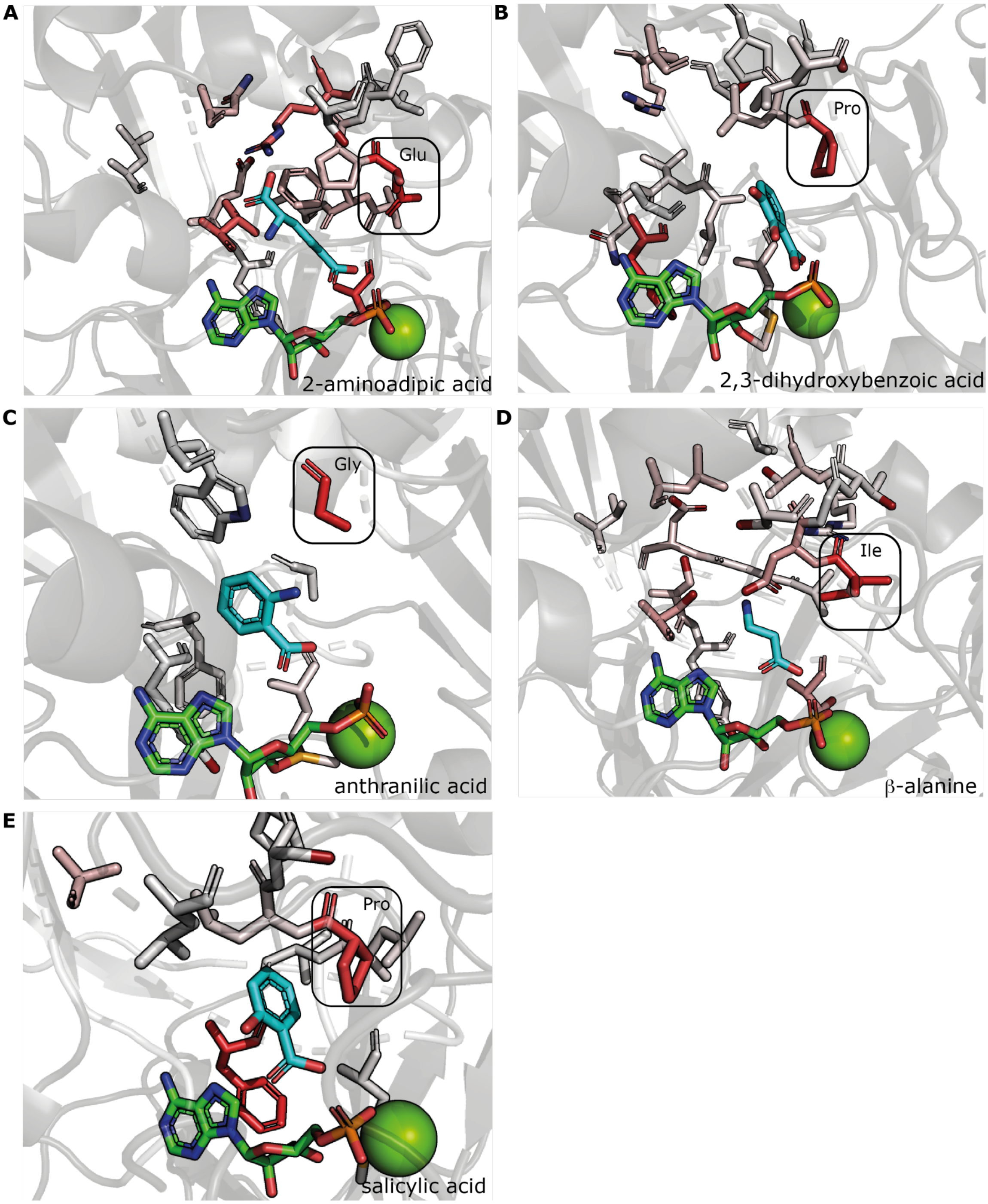

**Figure S18. Most predictive residues for non-amino acid substrates.** a) 2-aminoadipic acid. Note that this substrate is an amino acid; however it does not bind in the active site pocket like canonical amino acids, with the amino acid backbone sticking deep into the active site, its positive and negative charges coordinated by multiple polar and charged residues. b) 2,3-dihydroxybenzoic acid. c) Anthranilic acid. d) β-alanine. The backbones of β-amino acids are 1 atom longer than canonical amino acids, leading to a different mode of recognition. The amino group is stabilised by multiple positively charged and polar residues in the active site pocket. e) Salicylic acid. Predictive residues are visualised as sticks. The redder the residue, the more it contributes to predicting the indicated substrate.The 6th residue of the extended active site signature, which is usually the highly conserved aspartic acid residue that stabilises the amino acid backbone, has been indicated with a black box. For all 5 substrates, this residue is not an aspartic acid and is highly predictive.

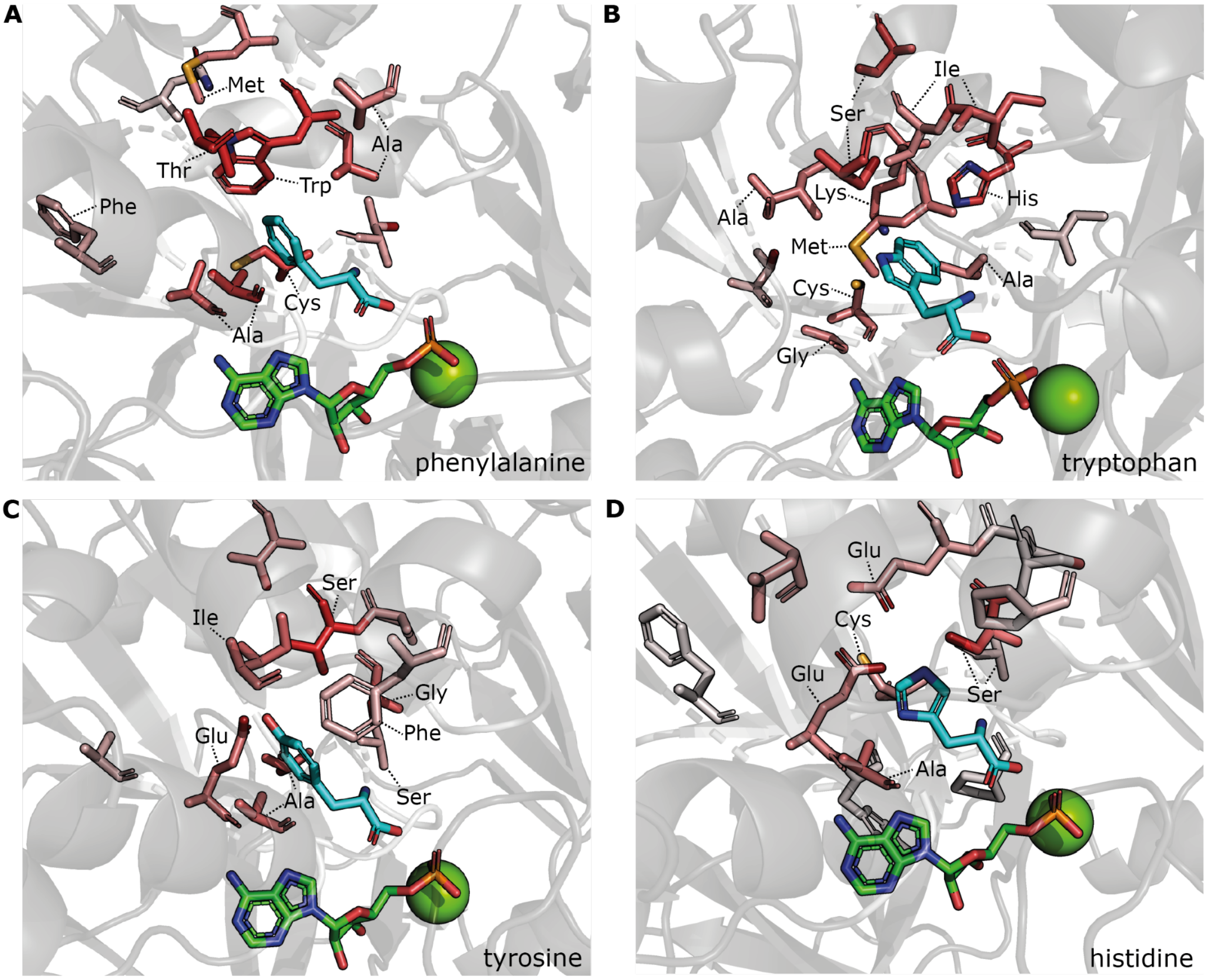

**Figure S19. Most predictive residues for proteinogenic aromatic substrates.** a) Phenylalanine. In this A domain, a large hydrophobic tryptophan residue caps the roof of the active site. Smaller hydrophobic alanine and cysteine residues line the rest of the active site. b) Tryptophan. A combination of hydrophobic (alanine, cysteine, methionine, isoleucine) and polar/charged (serine, lysine, histidine) residues form the active site pocket, coordinating the polar pyrrole and hydrophobic benzene of the indole side chain of the tryptophan substrate. c) Tyrosine. A combination of hydrophobic (isoleucine, phenylalanine, alanine) and polar/charged (glutamic acid, serine) residues form the active site pocket, coordinating the polar hydroxyl group and hydrophobic benzene ring of the tyrosine side chain. d) Histidine. Various negatively charged and polar residues stabilise the positively charged imidazole side chain of the histidine substrate. Predictive residues are visualised as sticks. The redder the residue, the more it contributes to predicting the indicated substrate. These large aromatic amino acids, particularly tryptophan, phenylalanine, and tyrosine, have a relatively high number of highly predictive residues, in contrast to some other substrates. This can be explained by the large number of residues involved in recognising the active site for large substrates, in addition to substrates from different phylogenetic clades recognising these substrates in different orientations, hence requiring a set of features describing a broader set of residues to make correct predictions.

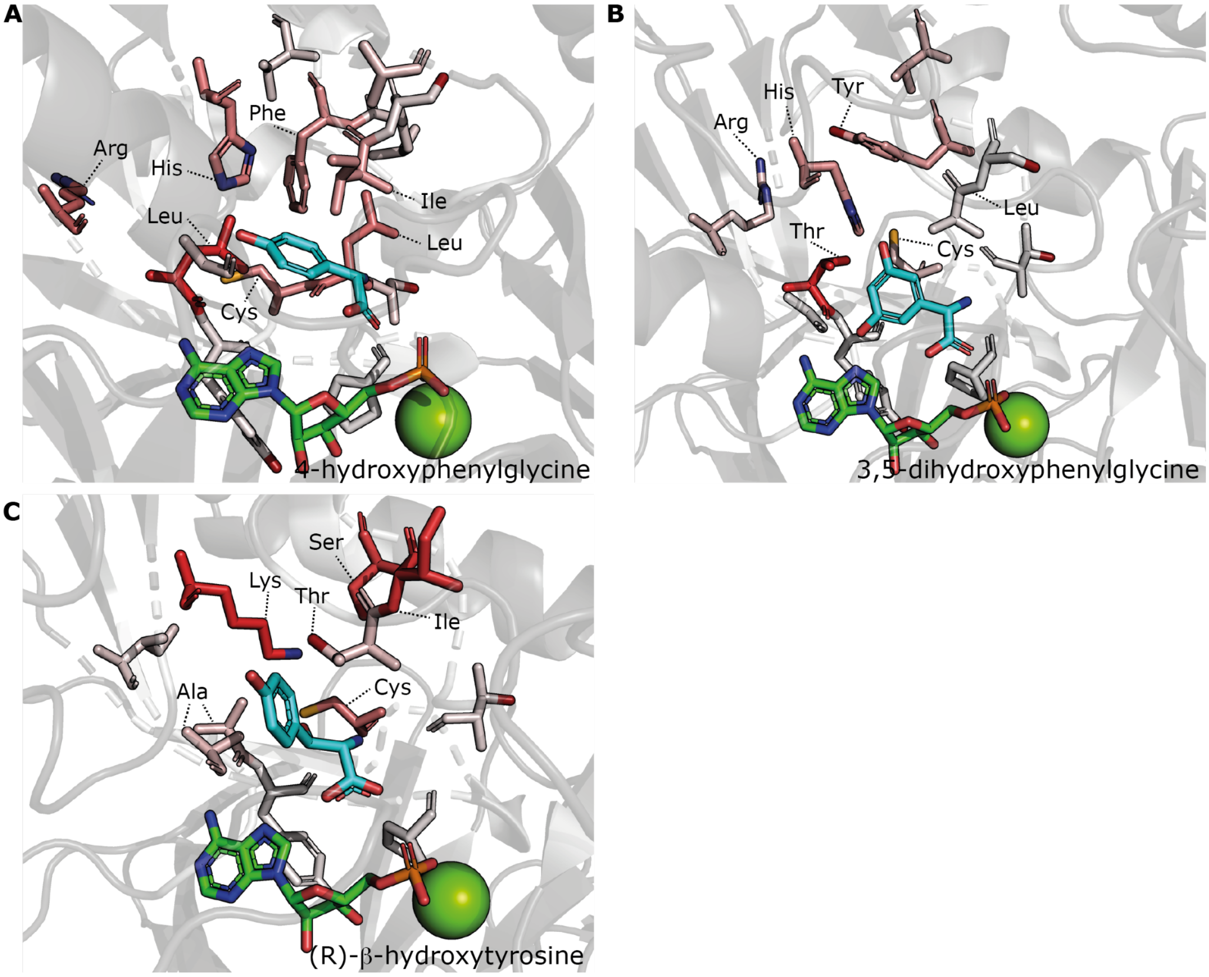

**Figure S20. Most predictive residues for nonproteinogenic aromatic substrates.** a) 4-hydroxyphenylglycine. The benzene ring is coordinated by hydrophobic residues (phenylalanine, isoleucine, leucine, cysteine), and two positively charged residues (arginine, histidine) create a polar pocket for the hydroxyl group. b) 3,5-hydroxyphenylglycine. This substrate is recognised similarly to 4-hydroxyphenylglycine, with the important difference that some hydrophobic residues have been replaced with structurally similar polar residues (leucine->threonine, phenylalanine->tyrosine) to accommodate the extra polar hydroxyl group in the substrate side chain. c) (R)-β-hydroxytyrosine. The polar and charged lysine, threonine and serine residues form a polar pocket for the β-hydroxyl and 4-hydroxyl groups. Small hydrophobic amino acids (alanine, cysteine) stabilise the hydrophobic benzene ring.

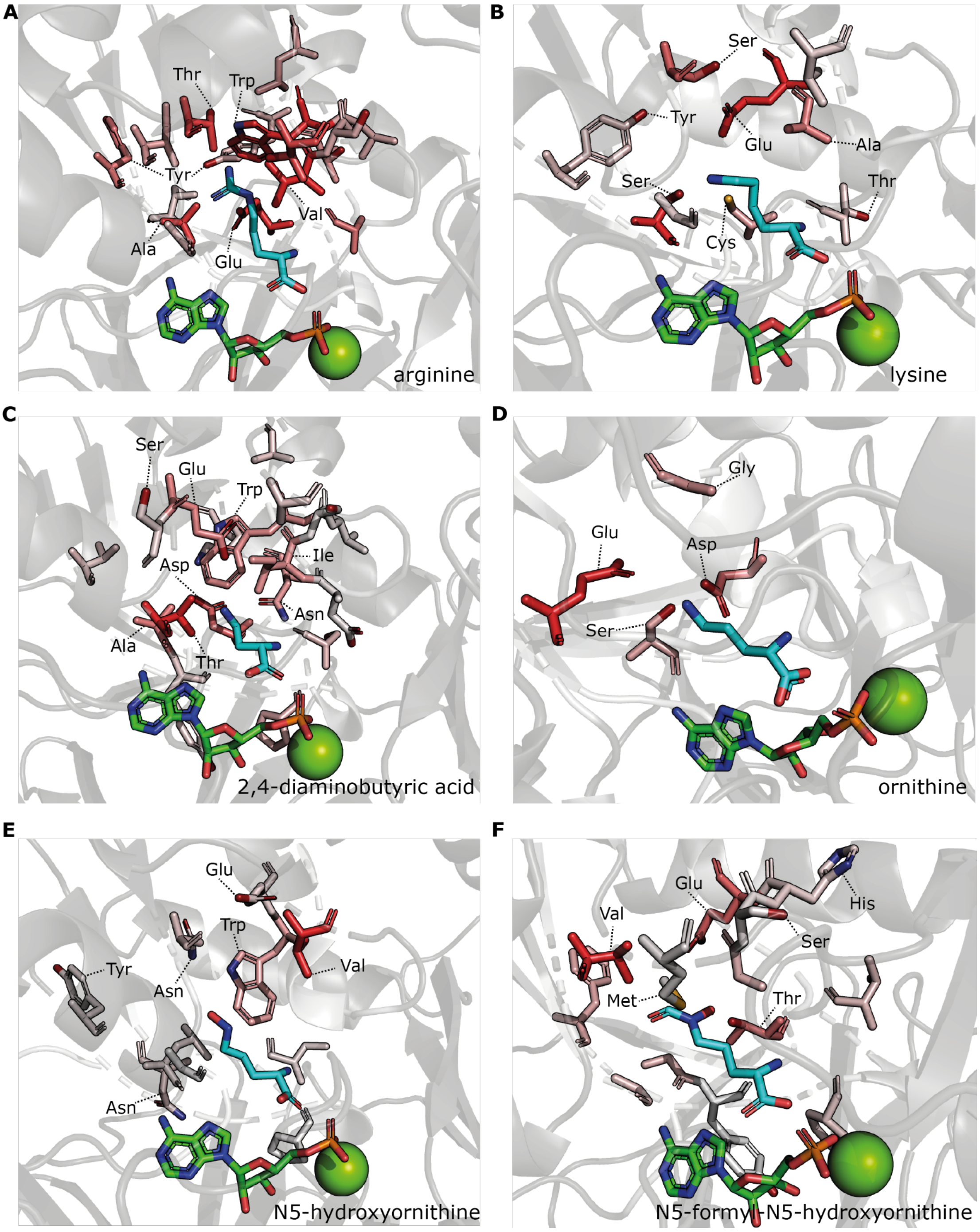

**Figure S21. Most predictive residues for negatively charged and polar substrates.** a) Arginine. The negative charge is stabilised by negatively charged (glutamic acid) and polar (tyrosine, threonine) residues. Small hydrophobic residues (valine, alanine) line the active site channel coordinating the carbon chain. A tryptophan residue caps the active site. b) Lysine, stabilised by positively charged (glutamic acid) and polar (serine, tyrosine, threonine) residues. c) 2,4-diaminobutyric acid. Polar (asparagine, threonine) and charged (glutamic acid) form a comparatively shallow polar active site pocket to accommodate the small, nitrogen-containing substrate. d) The negative charge of the ornithine substrate is neutralised by polar (serine) and charged (glutamic acid, aspartic acid) residues. e) N5-hydroxyornithine, which is slightly more neutral than its unhydroxylated variant, prefers asparagine and tryptophan to line its selecting A domain's active site pocket. f) This A domain, which selects N5-formyl-N5-hydroxyornithine, uses a combination of positively (glutamic acid) and negatively (histidine) charged residues as well as various polar (threonine, serine) and hydrophobic (valine, methionine) to coordinate its substrate.

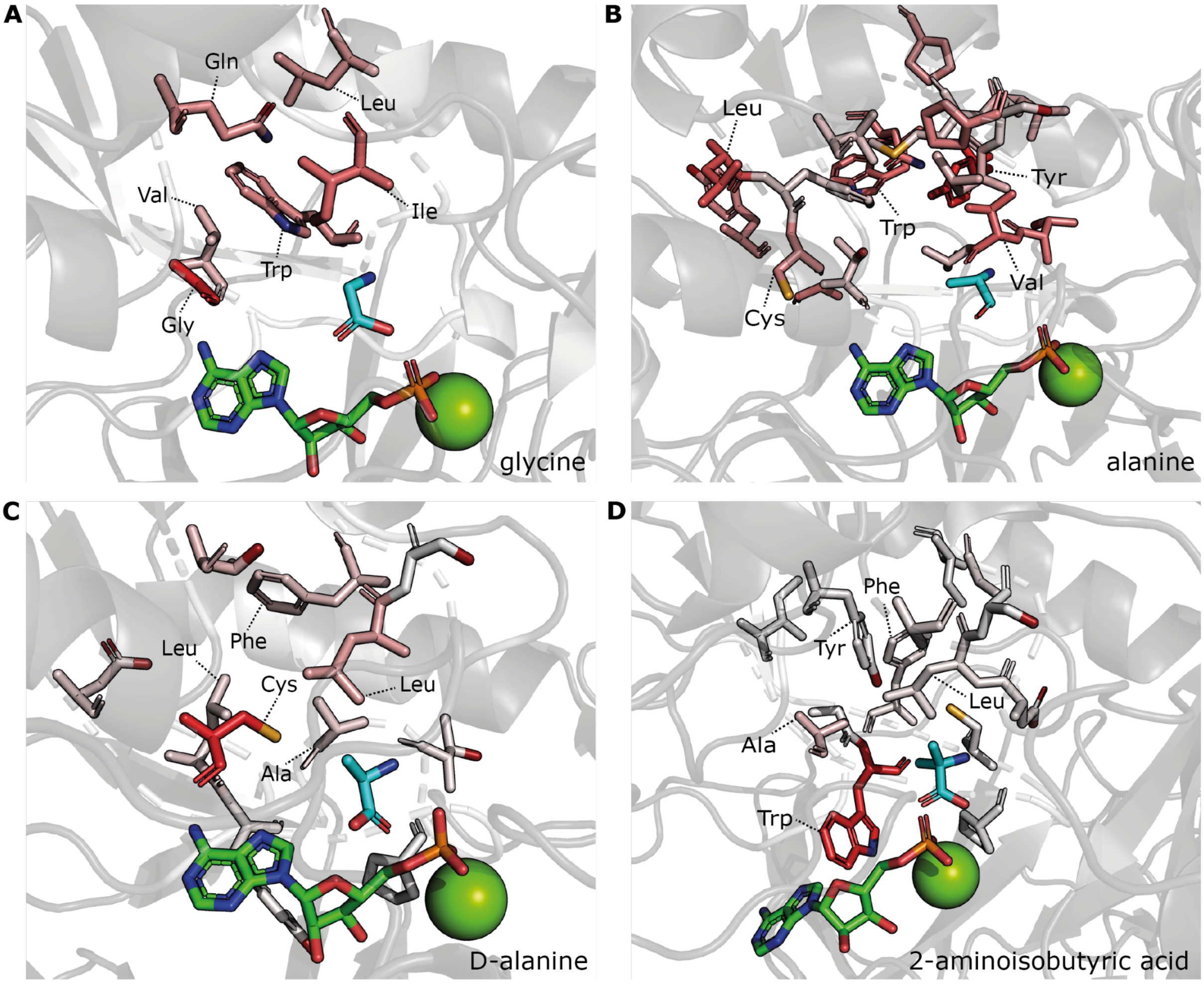

**Figure S22. Most predictive residues for small substrates.** a) Glycine. b) Alanine. c) D-alanine. d) 2-aminoisobutyric acid. For all substrates, large hydrophobic residues such as tryptophan, phenylalanine, isoleucine and leucine limit the size of the active site pocket.

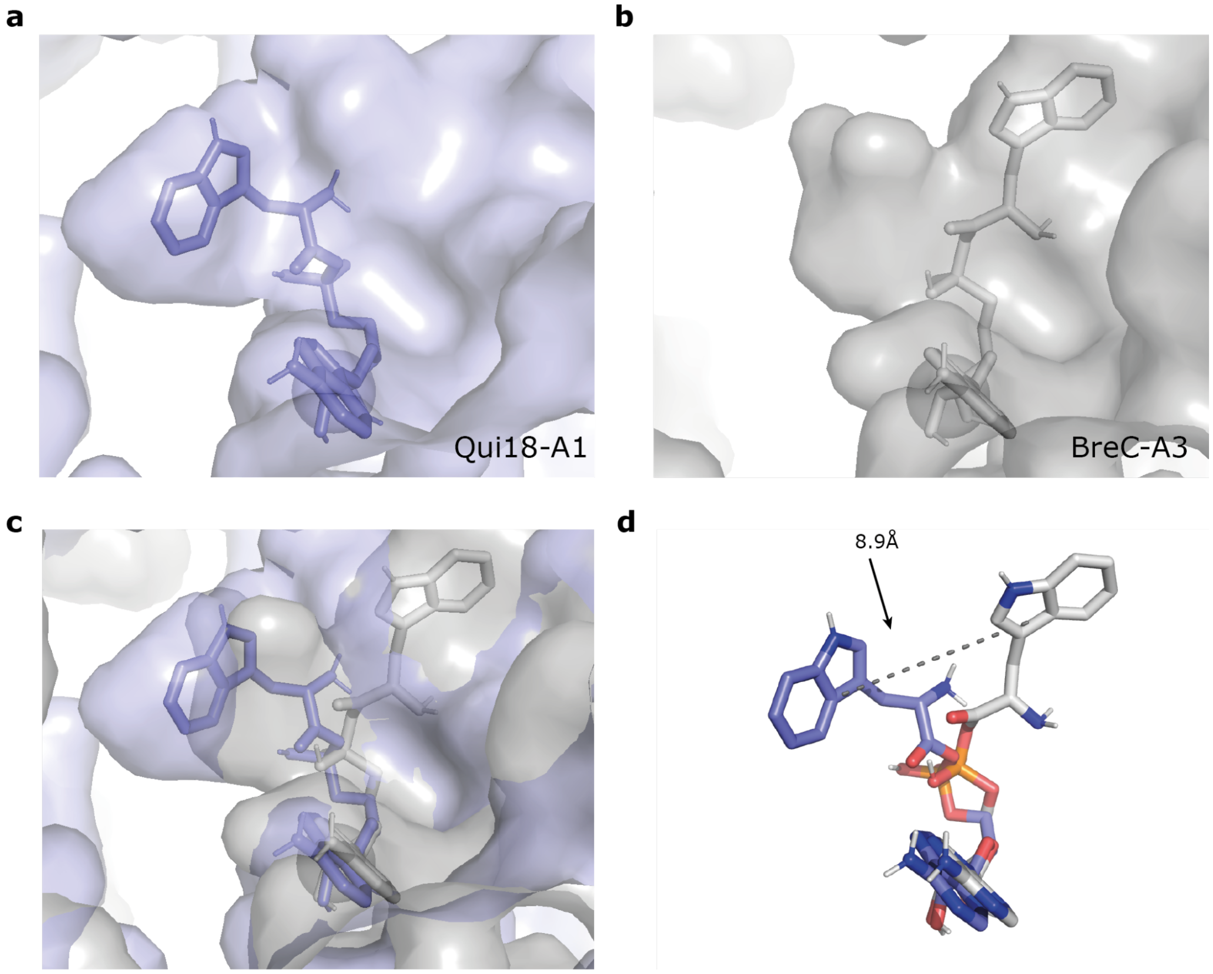

**Figure S23. TYM-docking to AlphaFold models of two phylogenetically and structurally divergent Trp-selecting A domains.** a) Tryptophan-adenylate (TYM) docked to Qui18-A1. b) TYM docked to BreC-A3. c) Comparison of TYM docked to Qui18-A1 (blue) and BreC-A3 (grey). The left pocket is smaller in BreC-A3 (grey) than in Qui18-A1 (blue), making it impossible for the tryptophan substrate to fit into this pocket in BreC-A3. Instead, the tryptophan moiety of TYM docked to BreC-A3 localises to a pocket on the right, leading to a 8.9Å shift (d; dashed line) of the central atom of the aromatic ring system within the tryptophan substrate with respect to the Qui18-A1 docking.

### Supplementary Tables

| Validation set | Structure-guided alignment | Sequence-guided alignment |
|---|---|---|
| 1 | 0.656 | 0.640 |
| 2 | 0.785 | 0.710 |
| 3 | 0.790 | 0.800 |
| 4 | 0.763 | 0.742 |
| 5 | 0.833 | 0.811 |
| 6 | 0.874 | 0.862 |
| 7 | 0.894 | 0.859 |
| 8 | 0.841 | 0.854 |
| 9 | 0.924 | 0.911 |
| 10 | 0.821 | 0.821 |
| Average | 0.818 (standard deviation = 0.076) | 0.801 (standard deviation = 0.0818) |

**Table S1. Structure-guided active site extraction boosts model performance.** We assessed the performance of ten random forest models following structure-guided active site extraction or sequence-guided active site extraction. Per substrate class, datapoints were randomised and subsequently distributed across the validation sets in order. As a result, validation sets with low indices contain a disproportionate number of datapoints for which there were few or no examples in the training set. This explains the drop in model performance in these sets.

| Tool | Setup time (s) | Time per domain (s) |
|---|---|---|
| **NRPSpredictor2** | 0.2355 | 0.0023 |
| **PARAS (all substrates)** | 9.826 | 0.0095 |
| **PARAS** | 1.8996 | 0.0118 |
| **AdenPredictor** | 1.7412 | 0.0131 |
| **NRPSTransformer** | 18.896 | 0.0178 |
| **PARASECT** | 1.5481 | 0.0328 |
| **PARAS (profile alignment)** | 1.8891 | 1.7337 |
| **PARASECT (profile alignment)** | 0.8633 | 1.7744 |
| **DeepAden** | 6.7505 | 2.2601 |
| **SANDPUMA** | 6.2682 | 61.192 |

**Table S2. Speed benchmark.** NRPSPredictor2, PARAS, PARASECT, AdenPredictor, DeepAden, and SANDPUMA were tested on a CPU-only server (256 CPUs) with no other processes running. NRPSTransformer was tested on a server with two L40S GPUs with no other processes running. For DeepAden and SANDPUMA, 1, 50, and 100 domains were run 4 times, removing outliers, to calculate the slope and intercept of the trendlines, which yielded setup time (intercept) and time per domain (slope). For all other tools, 1, 5, 10, 25, 50, 75, and 100 domains were run 5 times. Tools are sorted by per-domain computation time (ascending). Low speeds are indicated in green, high speeds in red.

| Domain | Actual substrate | PARAS | AdenPredictor | NRPSPredictor2 |
|---|---|---|---|---|
| **TriD.A1** | Val | Val | Val | Val |
| **TriD.A2** | Dab | Dab | Dab | Orn |
| **TriD.A3** | Gly | Gly | Gly | Gly |
| **TriD.A4** | Ser | Ser | Orn | Ser |
| **TriD.A5** | Trp | Trp | Tyr | No call |
| **TriD.A6** | Ser | Ser | Orn | Ser |
| **TriD.A7** | Dab | Dab | Dab | Orn |
| **TriD.A8** | Dab | Dab | Dab | Orn |
| **TriD.A9** | Trp | Trp | Tyr | No call |
| **TriD.A10** | Glu | Glu | Gln | No call |
| **TriE.A1** | Val | Val | Val | Val |
| **TriE.A2** | Ile | Ile | Leu | Ile |
| **TriE.A3** | Ala | Ala | Val | Val |

**Table S3. PARAS, AdenPredictor, and NRPSPredictor2 predictions of tridecaptin A5 BGC in *Paenibacillus* sp. JJ-21.** Red cells represent a wrongly predicted substrate. Grey cells indicate domains for which no substrate could be predicted. Dab: 2,4-diaminobutyric acid. Orn: Ornithine. Notably, PARAS made correct predictions for the two L-Trp-recognising A domains in TriD, which AdenPredictor and NRPSPredictor2 could not.

| Step | Screenshot | Description |
|---|---|---|
| 1 | 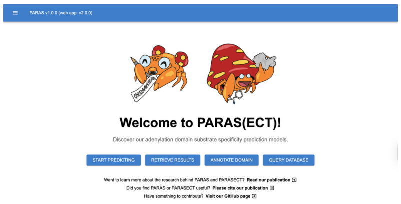 | The landing page of the PARAS and PARASECT web portal. Users can navigate between pages by pressing either the buttons for prediction, retrieval, domain annotation, or database querying, or using the drop-down menu from the navigation toolbar at the top. |
| 2 | 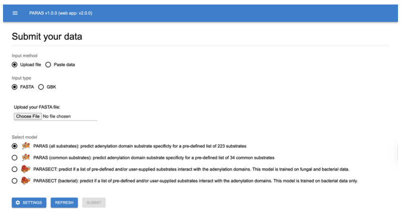 | Pressing the prediction button on the landing page will take the user to the submission page. The user can input FASTA or GenBank files and choose between the four PARAS and PARASECT models to make predictions. Additional settings are available upon clicking the settings button. Uploading a file or pasting the contents will prompt the submit button to be available. |
| 3 | 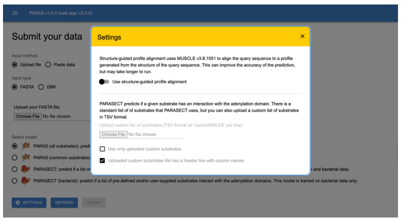 | Pressing the settings button will open a modal with additional settings for the user to set. Toggling the usage of the profile-guided structure alignment for active site extraction is available for all three PARAS and PARASECT models. Uploading a custom list of substrates to predict substrate specificity is only available for PARASECT. |
| 4 | 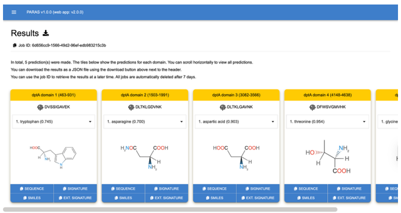 | After a few seconds to mine the input for A-domains and predict their substrate specificities, the results will be shown as a list of tiles. Every tile represents a single mined A-domain from the input with its predicted most likely substrate specificity. The user can toggle between all substrates. From this page, the user can download the results in JSON format. The provided job ID of the prediction task can be used to retrieve the same prediction results within seven days of generating them. |
| 5 | 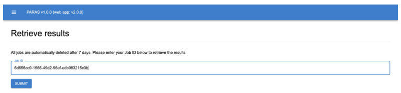 | Pressing the retrieval button on the landing page will take the user to the retrieval page. Here, the user can retrieve their previously generated result by submitting the job ID for that prediction job. Job results will automatically be deleted after seven days. |
| 6 | 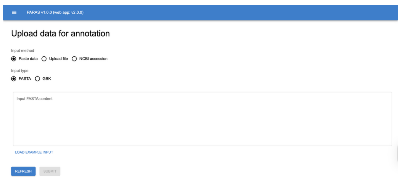 | Users can annotate and submit newly found A-domains, or re-annotate A-domains already present in the database using the user submission page. The annotation page uses the PARAS (all substrates) model by default. |

| | | |
|---|---|---|
| 7 | 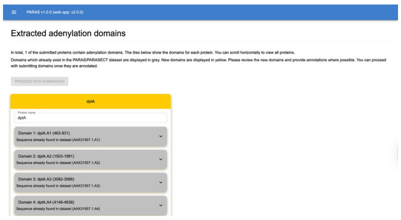 | After mining the user input for A-domains, the user is prompted to (re)annotate the domains found for each gene. |
| 8 | 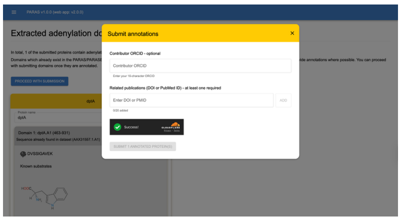 | After (re)annotating at least one domain, the user can proceed with the data submission. Providing an ORCID is optional, but at least one reference for the (re)annotation is required. Upon submission, a GitHub issue with the submission data is opened and publicly visible. |
| 9 | 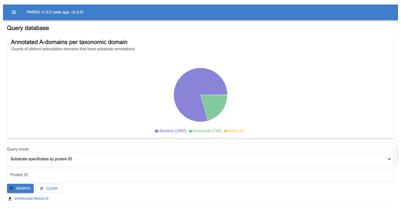 | Users can search the A-domain database using the database querying page. Several canned queries are available for the user to try, available in the query mode drop-down window. |
| 10 | 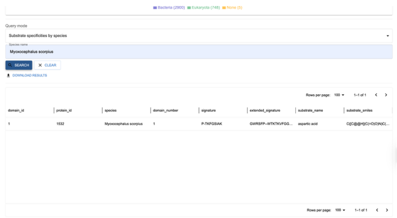 | An example result for retrieving all A-domains found in *Myoxocephalus scorpius*. The user can rearrange the data frame and download the results to TSV format. |

**Table S4.** Web portal (v2.0.0) instructions for PARAS and PARASECT.

| Primer | Oligonucleotide sequences (5' to 3') | Description |
|---|---|---|
| maedttp-L-for | gcgccgatggtttctacaaagatcgactagttcagctcacgtcggtcaccttccag | Left homologous arm for pCAP1000maedattp |
| maedttp-L-rev | tagccattttcttagttgttgtagggatggccaggacttccaggacatgcgcc | Left homologous arm for pCAP1000maedattp |
| maedttp-R-for | gagaagatgcggccagcaaaactaagatgtacgcggagaccgtcgtcttctggtgg | right homologous arm for pCAP1000maedattp |
| maedttp-R-rev | ttctaaatacaggtacctcaagtctcgagccagaacgaagtctccaagccccatgc | right homologous arm for pCAP1000maedattp |
| 1000-M-for | cctacaacaactaagaaaatggcta | Counter-selectable cassette from pCAP1000 |
| 1000-M-rev | ttagttttgctggccgcatcttctc | Counter-selectable cassette from pCAP1000 |
| spattp-L-for | gctgcgccgatggtttctacaaagatcgactagtcgccgtcggcgtagacgatccgca | Left homologous arm for pCAP1000spattp |
| spattp-L-rev | gcatgatagccattttcttagttgttgtaggaaggccgcctgggacccgaagaacat | Left homologous arm for pCAP1000spattp |
| spattp-R-for | tatttgagaagatgcggccagcaaaactaactcggaatggcccagcacatcgtc | right homologous arm for pCAP1000spattp |
| spattp-R-rev | ttctaaatacaggtacctcaagtctcgagcttgcgatcgcacgcaccagacgcc | right homologous arm for pCAP1000spattp |
| spattp-CK1-for | ctgtctgcaatacctcgacatcca | Check primers pair 1 for ttpBGC from *Streptomyces sparsogenes* |
| spattp-CK1-rev | ccaccacgacgtcctcacgg | Check primers pair 1 for ttpBGC from *Streptomyces sparsogenes* |
| spattp-CK2-for | gagcgcagaaaggtctttccg | Check primers pair 2 for ttpBGC from *Streptomyces sparsogenes* |
| spattp-CK2-rev | tcttccctcgcgacgagtga | Check primers pair 2 for ttpBGC from *Streptomyces sparsogenes* |
| spattp-CK3-for | accgacgactggtacaagggg | Check primers pair 3 for ttpBGC from *Streptomyces sparsogenes* |
| spattp-CK3-rev | ccacggagacgatgttgacgc | Check primers pair 3 for ttpBGC from *Streptomyces sp. maeda85* |
| maedttp-CK1-for | tcggagtagtcgtagatgttgtggc | Check primers pair 1 for ttpBGC from *Streptomyces sp. maeda85* |
| maedttp-CK1-rev | cagcgtcggcaagtcctatgtc | Check primers pair 1 for ttpBGC from *Streptomyces sp. maeda85* |
| maedttp-CK2-for | aaggccatgcgctgctgatg | Check primers pair 2 for ttpBGC from *Streptomyces sp. maeda85* |
| maedttp-CK2-rev | ctcgccacccgtctcgtctc | Check primers pair 2 for ttpBGC from *Streptomyces sp. maeda85* |
| maedttp-CK3-for | cagcgaacgggacctgttca | Check primers pair 3 for ttpBGC from *Streptomyces sp. maeda85* |
| maedttp-CK3-rev | ggggtgactccggtggacag | Check primers pair 3 for ttpBGC from *Streptomyces sp. maeda85* |

**Table S5.** Primer list for TAR cloning.

| Primers | |
|---|---|
| Forward primer PCR 1 | 5'-TGTCCTTCCAGGAACAGAACGGCAGTT-3' |
| Reverse Primer PCR 1 | 5'-TCATGCATCGTCGCCCTCCTTACGG-3' |
| Forward primer PCR 2 | 5'-AATAACATATGGCCGACGAGCGTGC-3' |
| Reverse primer PCR 2 | 5'-AATAAGAATTCTCATGCATCGCTGCCCTC-3' |

| Reaction setup PCR 1 | | |
|---|---|---|
| **Reactant** | **Volume** | **Concentration** |
| ThermoFisher Scientific HF Phusion mix | 12.5 µl | 2x |
| $dH_2O$ | 7.0 µl | |
| DMSO | 2.0 µl | |
| *S. sparsogenes* genomic DNA | 1.5 µl | 73.9 ng/µl |
| Forward primer PCR 1 | 1.0 µl | 10 µM |
| Reverse primer PCR 1 | 1.0 µl | 10 µM |
| **Program phase** | **Temperature** | **Time** |
| Start-up | 98°C | 1m |
| Cycles (30x) | | |
| Denaturation | 98°C | 30s |
| Annealing | 66.3°C | 30s |
| Extension | 72°C | 2m45s |
| Cooldown | 72°C | 10m |
| Store | 4°C | ∞ |

| Reaction setup PCR 2 | | |
|---|---|---|
| **Reactant** | **Volume** | **Concentration** |
| ThermoFisher Scientific HF Phusion mix | 12.5 µl | 2x |
| $dH_2O$ | 7.0 µl | |
| DMSO | 2.0 µl | |
| Amplified product PCR 1 | 1.5 µl | 39.7 ng/µl |
| Forward primer PCR 2 | 1.0 µl | 10 µM |
| Reverse primer PCR 2 | 1.0 µl | 10 µM |
| **Program phase** | **Temperature** | **Time** |
| Start-up | 98°C | 1m |
| Cycles (30x) | | |
| Denaturation | 98°C | 30s |
| Annealing | 65°C | 30s |
| Extension | 60°C | 2m30s |
| Cooldown | 72°C | 10m |
| Store | 4°C | ∞ |

**Table S6.** PCR primers and protocols for A domain cloning.

| | PARAS (substrate) | PARAS (taxonomy) | PARASECT (substrate) | PARASECT (taxonomy) | PARASECT bacterial (substrate) | PARASECT bacterial (taxonomy) |
|---|---|---|---|---|---|---|
| all | 0.88 | 0.83 | 0.83 | 0.80 | 0.89 | 0.84 |
| 2,3-diaminopropionic acid | 0.50 | 0.50 | 1.00 | 1.00 | 1.00 | 1.00 |
| 2,3-dihydroxybenzoic acid | 0.86 | 1.00 | 1.00 | 1.00 | 1.00 | 1.00 |
| 2,4-diaminobutyric acid | 0.93 | 0.83 | 0.79 | 0.63 | 0.93 | 0.63 |
| 2-aminoadipic acid | 1.00 | 0.67 | 0.67 | 0.67 | N/A | N/A |
| 2-aminoisobutyric acid | 0.94 | 0.90 | 0.88 | 0.93 | N/A | N/A |
| 3,5-dihydroxyphenylglycine | 1.00 | 1.00 | 0.90 | 0.13 | 1.00 | 0.13 |
| 4-hydroxyphenylglycine | 1.00 | 1.00 | 0.91 | 0.95 | 0.86 | 0.95 |
| 4R-hydroxyproline | 0.67 | 0.00 | 0.67 | 0.00 | N/A | N/A |
| D-alanine | 0.33 | 0.00 | 0.00 | 0.00 | N/A | N/A |
| N5-acetyl-N5-hydroxyornithine | 1.00 | 0.80 | 0.67 | 1.00 | N/A | N/A |
| N5-formyl-N5-hydroxyornithine | 1.00 | 0.17 | 0.33 | 0.33 | 1.00 | 0.22 |
| N5-hydroxyornithine | 1.00 | 1.00 | 1.00 | 0.70 | 1.00 | 0.91 |
| R-beta-hydroxytyrosine | 0.83 | 0.00 | 1.00 | 0.00 | 1.00 | 0.00 |
| alanine | 0.87 | 0.86 | 0.78 | 0.87 | 0.89 | 0.89 |
| anthranilic acid | 1.00 | 0.00 | 0.80 | 0.00 | N/A | N/A |
| **arginine** | 0.86 | 0.63 | **0.79** | **0.71** | 0.71 | 0.63 |
| asparagine | 0.96 | 1.00 | 0.96 | 0.88 | 0.96 | 1.00 |

| | | | | | | |
|---|---|---|---|---|---|---|
| aspartic acid | 0.89 | 0.54 | 1.00 | 0.69 | 0.94 | 0.57 |
| benzoic acid | N/A | N/A | 0.00 | 0.00 | N/A | N/A |
| beta-alanine | 0.75 | 0.20 | 0.38 | 0.67 | 0.60 | 0.60 |
| cysteine | 1.00 | 0.83 | 0.92 | 0.89 | 0.95 | 1.00 |
| glutamic acid | 0.73 | 0.76 | 0.58 | 0.05 | 0.82 | 0.11 |
| glutamine | 0.83 | 0.91 | 0.96 | 0.90 | 0.89 | 1.00 |
| glycine | 0.90 | 0.85 | 0.88 | 0.92 | 0.97 | 0.93 |
| histidine | 0.25 | 1.00 | 0.50 | 0.50 | 0.75 | 0.50 |
| homoserine | 1.00 | 0.00 | 0.67 | 0.08 | N/A | N/A |
| isoleucine | 0.79 | 0.76 | 0.59 | 0.44 | 0.61 | 0.65 |
| leucine | 0.93 | 0.90 | 0.91 | 0.96 | 0.94 | 0.98 |
| lysine | 0.90 | 0.63 | 0.60 | 0.44 | 0.73 | 0.36 |
| ornithine | 0.89 | 0.77 | 0.50 | 0.30 | 0.33 | 0.60 |
| phenylalanine | 0.78 | 0.82 | 0.58 | 0.77 | 0.82 | 0.75 |
| pipecolic acid | 0.75 | 0.40 | 0.25 | 0.00 | 0.33 | 0.00 |
| proline | 0.94 | 0.71 | 0.76 | 0.85 | 0.88 | 0.92 |
| serine | 0.88 | 0.90 | 0.89 | 0.85 | 0.95 | 0.90 |
| threonine | 0.99 | 0.98 | 0.97 | 0.99 | 0.97 | 0.99 |
| **tryptophan** | 0.83 | 0.36 | **0.71** | **0.57** | 0.77 | 0.52 |
| tyrosine | 0.82 | 0.77 | 0.88 | 0.85 | 0.89 | 0.90 |
| valine | 0.70 | 0.80 | 0.85 | 0.91 | 0.95 | 0.90 |

**Table S7.** Per-substrate accuracy of PARAS and PARASECT models on a substrate-stratified or taxonomy-stratified hold-out test set. PARASECT accuracy on tryptophan and arginine-selecting domains, discussed in the main text, are highlighted in bold.
